# Cognitive cartography of mammalian brains using meta-analysis of AI experts

**DOI:** 10.64898/2025.12.01.691701

**Authors:** Andrea I. Luppi, Hana Ali, Zhen-Qi Liu, Filip Milisav, Arianna Sala, Jitka Annen, Alessandro Gozzi, Danilo Bzdok, Bratislav Misic

## Abstract

No individual mind can fully grasp the complexity of modern neuroscientific literature. Where human experts flag, the latest AI models excel. Here we show how frontier large language models can systematically and quantitatively summarise literature-wide neuroscientific knowledge about the cognitive functions and dysfunctions associated with each brain region. Meta-analysis of these AI experts reveals structure-function mappings to which existing meta-analytic frameworks are blind, demonstrated by causal manipulations including lesions and intracranial stimulation. It also unlocks the possibility of extending quantitative literature meta-analysis and decoding of brain maps to other model organisms beyond human. As proof-of-concept, we integrate LLM meta-analysis with species-specific transcriptomics and connectomics in human, macaque, and mouse, discovering that the human molecular circuit for cognition is evolutionarily conserved. Altogether, literature synthesis with AI experts can fundamentally catalyze neuroscientific discovery by overcoming the barrier of data aggregation from heterogeneous studies, finally bringing together a scattered literature to identify latent insights across disparate subfields, modalities, and species.

## INTRODUCTION

A fundamental goal of neuroscience is to understand how the architecture of the brain enables cognition. Yet no single mind can grasp the deluge of findings about causal and correlational associations between brain structure, function, and dysfunction obtained from neuroimaging, brain stimulation, and brain lesions in humans and other animals (1–10). One way to make sense of this vast and tangled literature is through metaanalysis: the de facto strategy for evidence synthesis in science, offering a structured approach to integrate neuroimaging studies and summarise brain maps of the anatomical contributions to each cognitive operation (11–24).

Manually-curated meta-analysis (11, 25) is extremely laborious, and increasingly unfeasible given the exponential growth of the literature. However automated engines for coordinate-based meta-analysis (12, 13) are largely or entirely restricted to MRI or PET, and only available for human studies (19, 22–24, 26). These restrictions preclude translational discovery from key animal models, and from huge swaths of the literature using other methods for recording, imaging, and causally manipulating brain function.

Where human experts flag, the latest AI models excel: large language models (LLMs) based on the transformer architecture (27–32). Popular models such as OpenAI’s `Generative Pre-trained Transformer` (GPT) (33), Anthropic’s Claude (34), Google’s `Gemini and Gemma` (35, 36), Meta’s Llama (34), Alibaba’s Qwen (37), DeepSeek AI’s DeepSeek (38) and Mistral AI’s `Mistral`(39), are pre-trained on vast corpora of text including scientific literature, effectively making them ‘AI experts’ across multiple domains of knowledge. Indeed, LLMs demonstrate a remarkable capacity to summarise related technical information and high-level concepts across diverse domains of science – from genomics to medicine to cognitive science (40–49) – and across different techniques, model organisms, and sub-fields (42, 48, 50–52). In neuroscience, recent work revealed that popular pretrained LLMs exhibit remarkable domain-expertise, systematically outperforming human experts at predicting the outcome of neuroscience experiments (51).

Here we introduce NeuroNaut: fully automated meta-analysis of AI experts for neuroscience, combining key advantages of automation and domain-expertise. We use publicly accessible general-purpose LLMs as ‘artificial experts’ to systematically summarise literature-wide neuroscientific evidence about the cognitive functions and dysfunctions associated with each brain region. Rather than training a new model from scratch, this approach takes advantage of the vast literature corpus embedded in LLMs’ pre-training to automatically produce accurate mappings between brain regions and cognitive operations, without restrictions to specific techniques or modalities. We combine and compare 3 top-tier LLMs as automated neuroscience experts: OpenAI’s `GPT-5.4`, Google’s `Gemini 3.1 Pro`, and Anthropic’s `Claude-4.6-Opus`. We replicate the main results with flagship open-weight LLMs: OpenAI’s OSS-120b, Google’s `Gemma-4-31b-instruct`, and Alibaba’s `Qwen3-235b-a22b-2507`. Our approach allows users to choose from over 300 proprietary and open-weight LLMs. Among them, we benchmark cost and performance for a portfolio of 20 LLMs from the GPT, Claude, Gemini, OSS, Gemma, Qwen, Mistral, Llama, and DeepSeek model families, encompassing instruction-tuned, reasoning, dense, and mixture-of-experts models (33–39, 53, 54). We extensively validate brain maps generated by AI experts against functional, structural, metabolic, connectomic, transcriptomic, and chemoarchitectonic databases (1, 55). We show that brain maps of cognition assembled from AI experts’ summaries of the literature are consistent with the summaries provided by current meta-analytic engines (12–14). At the same time, they go beyond existing tools by providing additional insights to which current field-standard meta-analytic engines are blind, recapitulating lesion- and stimulation-based maps of human cognitive circuits (6, 56–58), and shedding light on brain-disorder and disorder-disorder associations that were latent in the scientific literature. Crucially, using AI experts to summarise the neuroscientific literature unlocks the possibility of extending meta-analysis into the domain of other model organisms where meta-analytic engines are not currently available, which we showcase here with macaque (59–61) and mouse (62–65).

The present approach does not involve any *de novo* model training, because it relies on at-scale querying of pre-trained LLMs that are publicly available. It is therefore fundamentally distinct from recent foundation models for fMRI, which are trained from scratch on the statistical structure of thousands of hours of fMRI scans and learn to simulate brain responses to specific stimuli, tasks, or conditions (66–68). We provide NeuroNaut as an openly available framework (https://github.com/andrealuppi/NeuroNaut) to bring together, summarise, and make accessible evidence that lay scattered across the literature used to pre-train frontier LLMs (47, 48, 51) – making it possible to ‘ask an expert’ systematically and at scale. We demonstrate that LLM experts can fundamentally catalyze neuroscientific discovery by automating data aggregation from heterogeneous studies, overcoming the barrier of terminological variations across research domains, and identifying emergent patterns and novel connections across disparate subfields, species, and modalities.

## RESULTS

The openly available NeuroNaut framework uses pretrained, general-purpose large language models as artificial experts, to systematically estimate the relative associations between brain regions and cognitive functions. LLMs’ linguistic competence and extensive neuroscientific evidence-base enables them to recognise different names for the same brain region, much like a human expert would (in the Supplementary Results we provide evidence in support of this claim). This enables automated, systematic mapping of function to anatomy at scale, without requiring specific coordinates or narrowly focusing only on the human literature, as required by currently used reference solutions for coordinate-based meta-analysis in the neuroscience community (e.g., NeuroSynth, NeuroQuery (12, 13)). Thus, a framework of LLMs as AI experts supports multiple species and brain atlases to enable comprehensive comparative analysis. Here we focus on three species: human, macaque (*Macaca mulatta*), and mouse (*Mus musculus*).

We here foreshadow the main results to highlight the guiding logic, before describing each in detail. To assess whether AI experts can provide good summaries of the literature about regions’ functional roles, we adopt a triangulation approach. First, we ask whether the AI experts’ summaries are consistent with the summaries provided by existing meta-analytic engines. We find very broad agreement, across 8 fundamental domains but also a broad set of 123 categories. Second, we ask whether the AI experts can provide a synthesis that goes beyond what is available from existing tools. We show that AI experts’ brain maps tend to reflect alternative sources of brain-mapping evidence such as lesions and stimulation, better than NeuroSynth and NeuroQuery which are primarily or exclusively based on fMRI (12, 13). Third, we obtain summaries for macaque and mouse, which are excluded by current meta-analytic engines. We validate the resulting cognitive circuits against species-specific transcriptomic, connectomic, and functional maps (9, 73).

For the main results, we access three frontier LLMs from leading AI companies as automated neuroscience experts: OpenAI’s GPT-5.4, Anthropic’s `Claude-4.6-Opus` and Google’s `Gemini 3.1` Pro. Models of the GPT-5, Claude-4, and Gemini-3-Pro families are consistently ranked as the current best-performing for biological and biomedical domain-knowledge (47, 74, 75). We also replicate the main results with three flagship open-weight LLMs: OpenAI’s OSS-120b, Google’s `Gemma-4-31b-instruct`, and Alibaba’s `Qwen3-235b-a22b-2507`. To ensure robust and diverse analytical perspectives and a broader range of training corpora, we further benchmark the metaanalytic performance and cost of a broader range of 20 proprietary and open-weight LLMs, spanning the GPT, Claude, Gemini, DeepSeek, Mistral, Llama, Qwen, Gemma, and OSS families. Models are accessed using OpenRouter’s direct API integration, but to democratise access and provide users with maximum flexibility, our framework also supports use of fully-local LLMs: free of charge, and without need for API keys or web access (see Methods for details of both approaches).

### Cognitive cartography with AI experts

We begin by using the NeuroNaut framework to query AI experts for the involvement of each region in specific functions representing broad domains of cognition. Each LLM is provided with a brain region *X*, a species *Y*, and a cognitive function *Z* as inputs; it is prompted to take on the persona of a neuroscience expert, and estimate the probability that brain region *X* of species *Y* is involved in function *Z*, summarising the frequency of association between region and function across the published literature (see *Methods* for details of the exact prompt). Responses are parsed to extract decimal scores between 0 and 1. We refer to these outputs as region-function association scores since they are not formal probabilities. Analyses rely on the relative ranking of regions, rather than on the absolute calibration of the reported values, because the goal is rather to estimate the *relative* importance of different brain regions for the same function.

We can then assemble quantitative brain maps for specific cognitive domains by querying each LLM about the same cognitive function systematically for each brain region (for the main analysis we use the 68 regions of the Desikan-Killiany atlas (70); results for alternative parcellations (multimodal and cytoarchitectonic) are provided in the Supplementary Results). After obtaining maps for each AI expert, we generate maps of meta-analytic consensus among the AI experts by (i) z-scoring each map to remove differences in scale between LLMs, which allows us to focus on the relative differences among regions; and then (ii) averaging the z-scored maps across all LLMs. In the Supplementary we show that the resulting maps of region-function association are highly correlated with maps obtained through an alternative approach: explicitly asking the LLMs to compare the relative importance of all possible pairs of regions for a specific function, and turning the resulting pairwise comparisons into region-wise maps of formal probabilities using the Bradley-Terry model (76).

Can automated neuroscience experts accurately map the relative importance of different brain regions for specific cognitive functions? As proof of concept we focus on eight fundamental cognitive domains, identified by a recent computational ontology as providing a representative data-driven synthesis of human cognition: emotion, cognitive control, listening, memory, language, vision, reward, and manipulation (69). Remarkably, even though LLMs are only ever asked about individual regions (i.e., no LLM is ever asked to produce an entire brain map), assembling LLMs’ opinions across regions does in fact produce brain maps of cognition that match well our own intuition as neuroscience experts (Fig. 1b) (including for example the left-lateralisation for the language map; Fig. S1). In other words, each AI expert provides region-function association scores that are internally calibrated for that expert, such that greater scores are assigned to regions with greater functional relevance. This is confirmed by the very high correlations observed between the rankings produced by different individual AI experts (Fig. S2), thereby supporting the validity of aggregating them (we also return to this point in the Supplementary Results).

**Figure 1.**
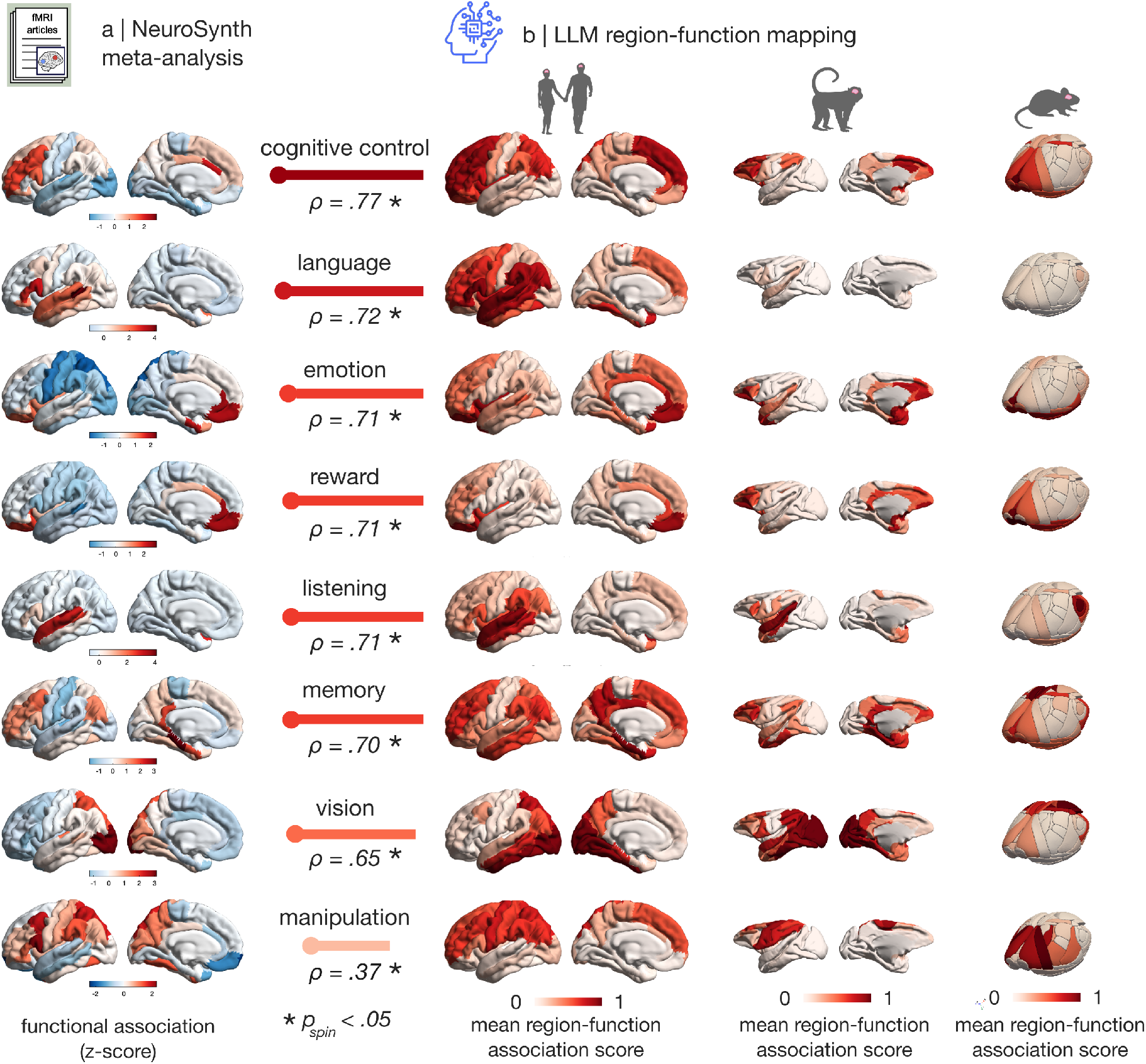
LLMs generate brain maps of cognition. (a) NeuroSynth meta-analytic maps term-based meta-analysis of the fMRI literature) for 8 representative domains of cognition identified through data-driven computational ontology (69)). Brain maps show, for each region, NeuroSynth’s measure of association strength between terms and voxels, quantified as the z-score from a twoway ANOVA testing for the presence of a non-zero association between term use and voxel activation (12). Note that NeuroSynth does not distinguish between areas that are activated or deactivated in relation to the term of interest, and a negative number does not imply deactivation or inhibition. To enable comparison, the voxelwise NeuroSynth maps are parcellated by averaging the values of the voxels belonging to the same region of the Desikan-Killiany atlas (see Methods). (b) AI experts generate plausible brain maps of cognition for human, macaque and mouse. Maps display, for each cortical region, the corresponding region-function association scores: LLM-estimated probabilities that each region is involved in the corresponding function (see Methods for the exact prompt, and see Supplementary Results for an alternative method of eliciting probabilities). A summary map is obtained by averaging the values assigned to the same region by different LLMs (here: `GPT 5.4; Gemini 3.1 Pro; and Claude 4.6 Opus`). Note that for analysis, each LLM map is z-scored prior to the averaging step, but here we show the average of the non-z-scored maps for ease of interpretation. See Fig. S7 for the same figure, but with LLM-generated brain maps z-scored prior to averaging; all 8 LLM-generated maps have Spearman’s *r >* 0.99 between the z-scored and non z-scored versions. Human: 68 cortical regions from the Desikan-Killiany atlas (70). Macaque: 82 cortical regions from the Regional Mapping atlas (71). Mouse: 72 cortical regions from the Allen common coordinate framework atlas (72). Horizontal bars indicate the Spearman correlation between the brain maps displayed at either side, with length proportional to the correlation magnitude. Scatter-plots for each correlation are shown in Fig. S8. Significance of correlations is assessed against a null population of surrogate maps with preserved spatial autocorrelation. See Fig. S9 for the corresponding null distributions. See Fig. S3 for replication of the human results with the NeuroQuery meta-analytic engine instead of NeuroSynth. For both (a) and (b), brain maps are displayed on the lateral and (for human and macaque) medial surface of the species-specific left cortical hemisphere.

To provide support for the plausibility of brain maps generated by automated neuroscience experts, we adopt a multi-pronged strategy. At the level of individual regions, we provide a `–justify` flag whereby users may request a 1-2 sentence textual justification alongside each LLM-provided score, to ensure transparency and provide insight into the automated experts’ rationale for the role of specific regions in specific functions (see Methods). Section (ii) of the Supplementary Results provides examples and quantitative analyses demonstrating that different AI experts’ explanations are consistent with each other and with our own neuroscientific expertise, as well as demonstrating their ability to recognise different names for the same brain region.

At the level of entire brain maps, we compare the maps of cognition generated by AI experts against corresponding meta-analytic maps generated by NeuroSynth (12) (Fig. 1a) and NeuroQuery (13) (Fig. S3). Indeed, all LLM-generated maps are significantly spatially correlated with NeuroSynth maps (beyond the effect of spatial autocorrelation), with all except “manipulation” exceeding a correlation magnitude of 0.60 (Fig. 1). In fact, in the Supplementary Results we demonstrate that the lower correspondence with the NeuroSynth map for “manipulation” actually militates *in favour* of the LLM experts, not against: the NeuroSynth map is contaminated by the broad meaning of the word, which is also used for example as “experimental manipulation” by researchers, alongside its meaning as the cognitive operation of study which refers to “object handling” or “symbol transformation” – resulting in paradoxically low values in the primary motor cortex of the NeuroSynth map. The results also replicate when comparing LLM-generated maps against an alternative meta-analytic engine, NeuroQuery (13) (Fig. S3). NeuroQuery employs techniques from (pre-LLM) natural language processing to infer semantic relationships among terms used in the literature, thereby combining the results of similar terms (13). Whereas NeuroSynth generates brain maps of the consistency of observations reported in the literature, the brain maps generated by NeuroQuery are *predictions of the probability* that a study on the cognitive operation of interest will report observations at each location (13). All correlations are again significant, with ‘manipulation’ now approaching Spearman’s *r* = 0.60. We also show that correspondence with NeuroSynth maps is not just due to aggregating the maps provided by different individual LLMs: individual LLMs also produce maps that are highly correlated with the corresponding NeuroSynth maps (Fig. S4) and with each other (Fig. S2). In the Supplementary Results, we show that LLMs’ ability to recapitulate maps of human cognition from NeuroSynth is not restricted to the broad categories of (69), but extends to a much more fine-grained ontology spanning 123 cognitive operations. We also provide a negative control, showing that AI experts consistently return all-zero maps when queried about regional importance for ‘telekinesis’ or ‘telepathy’, two putative mental faculties that have no documented scientific basis (Fig. S5).

Fig. 1 showcases the breadth of inquiry made possible by LLMs, in the sense that their neuroscientific expertise extends to other model organisms. We use the same procedure to generate cortical maps for the same eight cognitive domains in macaque and mouse, two widely used mammalian model organisms in neuroscience (77–79)(Fig. 1). Prima facie, the species-specific maps generated by consensus of AI experts recapitulate features of core functional neuroanatomy. For instance, in the macaque “vision” is associated with visual cortex, “listening” with superior temporal cortex and temporo-polar cortex (80), and “manipulation” with primary somatosensory and motor cortices, “reward” with orbitofrontal cortex, and “cognitive control” with dorsolateral prefrontal cortex. In mouse, the greatest associations with “vision” are likewise found in visual areas: VISal, VISam, VISl, VISp, VISpl. For “reward”, orbital, anterior cingulate (ACAd, ACAv), and limbic areas (PL, ILA) are prominent, and for “manipulation”, primary and secondary motor areas, as well as primary somatosensory areas. For “listening”, the auditory areas are most involved (AUDd, AUDp, AUDpo, AUDv).

The automated experts consistently attribute nearzero values for “language” to regions of the macaque and mouse brain: on average one order of magnitude smaller than other terms for the macaque, and two orders of magnitude smaller for the mouse (Fig. S6). However, if we consider the spatial pattern alone of relative association with “language”, the resulting maps are neither random nor trivial: in both species, the regions that are ranked as relatively (albeit to a very low extent) more associated with “language” display clear spatial overlap with the maps for “listening”, just as in the human (Fig. 1). In addition, these regions are in fact associated with species-specific vocalizations, including putative voice regions in superior temporal cortex and ventral-prefrontal and opercular cortex of the macaque brain that preferentially respond to species-specific vocalisations (80–82). In other words, the AI experts appropriately convey that finding language-associated regions in these non-human organisms is very unlikely, while also providing what may be regarded as sensible best guesses. In the next section, we assess the accuracy of the non-human maps in a more quantitative way.

### Cognitive cartography in non-human species

An important use of human meta-analytic databases such as NeuroSynth is to perform principled ‘cognitive decoding’: comparing a new brain map of interest against a pre-specified set of meta-analytic maps, to draw quantitative inferences about the underlying cognitive operations (83–86). Thanks to their advanced language-processing capacity, LLMs unlock the possibility of extending meta-analysis and cognitive decoding into the domain of other organisms. AI experts provide a twofold advantage: (i) they have access to a broader multimodal literature (e.g., not just human fMRI and PET); and (ii) they are not constrained by how studies report their results (e.g. with respect to a standardized coordinate system). Here we consider the capacity for AI experts to decode cortical maps for two fundamental model organisms for neuroscience: the macaque and the mouse.

For the mouse, we benchmark the generated maps against a recently-released flagship resource from the International Brain Laboratory consortium (9). Briefly, the dataset consists of cortical maps reconstructed using in vivo electrophysiological recordings of 621, 733 neurons recorded during a decision-making task with 699 Neuropixel probes in 139 mice (9). Importantly, the data were annotated with task components related to sensation, choice, action and reward, as well as internal cognitive states. Specifically, we use brain maps that report for each cortical region, the success (*R*^2^) of decoding. We then perform ‘cognitive decoding’ of each IBL map of task annotations, by spatially correlating it against the 8 representative cognitive maps for the mouse generated using consensus of AI experts. As Fig. 2a shows, we find that the meta-analytic maps display the greatest spatial correlation with the expected Neuropixel maps, such that “vision” is maximally correlated with the “stimulus” annotation, “manipulation” is maximally correlated with “choice” (behaviourally manifested by the animal using its paws to turn a wheel) as well as the “velocity” and “speed” of the wheel-turning, and “listening” is maximally correlated with the “feedback” annotation (feedback was provided as an aversive auditory tone, in case of incorrect choice)(Fig. 2a). Collectively, these results show that LLM maps can generate empirically realistic proxy maps, across domains of cognition and even across species, by summarising findings across the literature even when explicit meta-analytic results are not available.

**Figure 2.**
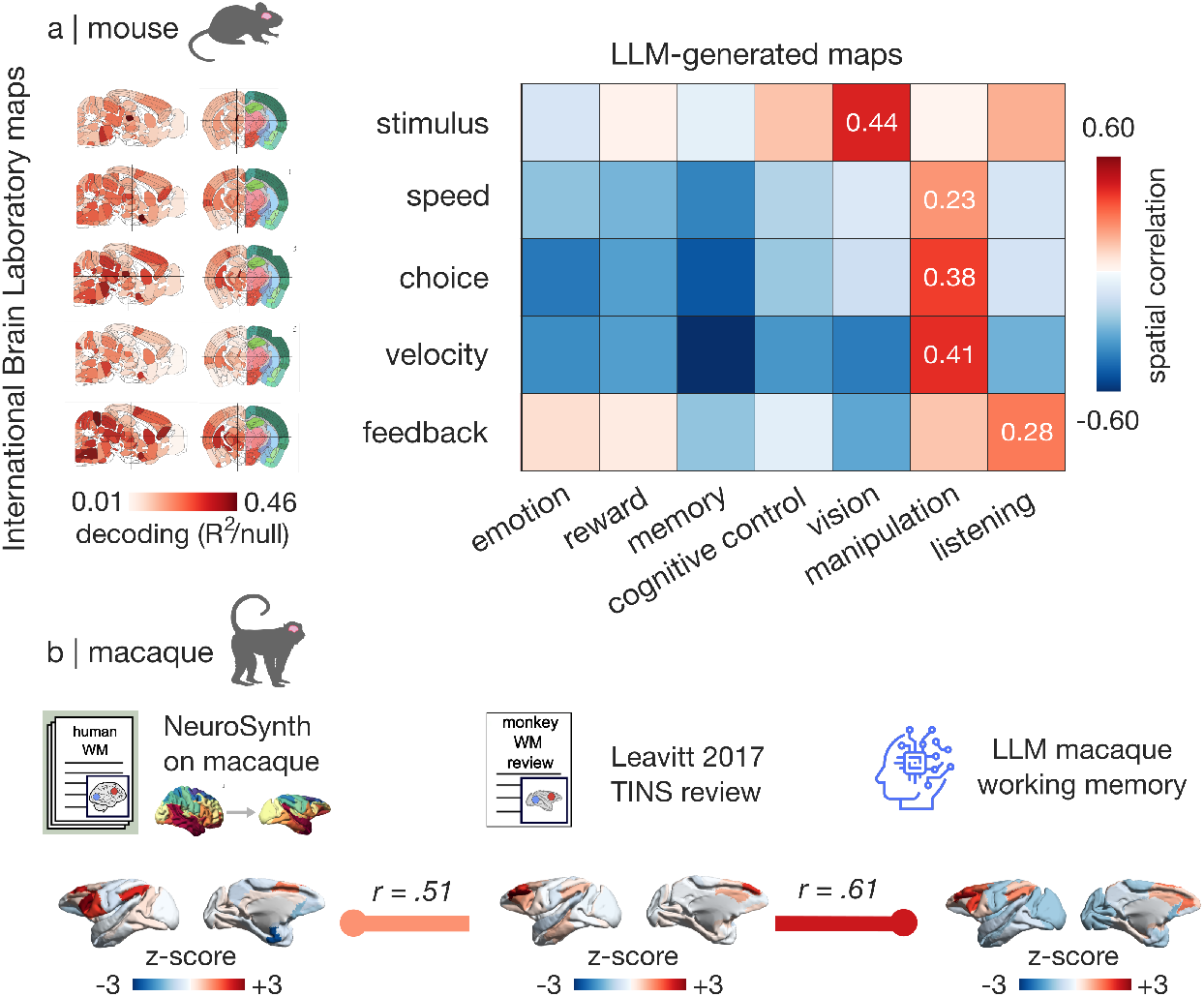
LLM decoding of mouse and macaque cortical maps. **(a)** Cortical maps of mouse cognition are obtained from recently released Neuropixels recordings from the International Brain Laboratory (9), corresponding to decoding success (*R*^2^ over null) for five separate components of a decision-making task: stimulus encoding; behavioural choice; velocity and speed of behaviour (turning of a wheel); and feedback (reward or aversive tone). Using LLM-generated maps for the mouse brain pertaining to 7 representative domains of cognition (see Fig. 1 for corresponding brain maps), we find that the best correlation with empirical Neuropixels maps are appropriate for each task stage. Mouse brain maps are displayed as coronal and sagittal slices, adapted from (9). **(b)** LLM meta-analysis of macaque working memory significantly recapitulates (*p <* 0.001 after accounting for spatial autocorrelation) available evidence about macaque working memory from Leavitt’s review (73), better than a NeuroSynth or NeuroQuery meta-analytic map of human working memory projected onto macaque brain. Horizontal bars indicate the Spearman correlation between the brain maps displayed at either side, with length proportional to the correlation magnitude (both *p <* 0.001, assessed against null distributions of surrogate maps with preserved spatial autocorrelation). See Fig. S11 for the corresponding null distributions. Macaque brain maps are displayed on the lateral and medial surface of the left cortical hemisphere. For ease of comparison, all maps are shown in units of z-score across regions (see Methods for the original units of each). Therefore a negative number simply means that the region in question has a lower value than the mean, but does not imply deactivation or inhibition.

For the macaque, there exists no comparable comprehensive dataset. The closest available is Leavitt’s (73) quantitative summary of the evidence of involvement of every macaque region in working memory (number of studies finding evidence in favour, or null, for each region). After resampling this map to the Regional Mapping atlas used for the macaque (see *Methods*), we show that it exhibits significant spatial correlation with the map of macaque working memory obtained from artificial experts (Fig. 2b). The phylogenetic proximity of the macaque to human allows us to ask an additional methodological question: is the map generated by AI experts macaque-specific, or is it simply evidence from the human literature projected to the macaque cortex? To address this question, we obtain a meta-analytic map of human working memory from NeuroSynth and parcellate it into the macaque Regional Mapping atlas (61, 71, 87), comparing it against (1) the LLM-derived map of macaque working memory; and (2) the summary of macaque working memory from Leavitt (73). We find that the macaque working memory map from Leavitt (which we treat as ground truth) is best matched by the LLM-derived map (Spearman’s *r* = 0.62, *p*_*Moran*_ *<* 0.001), outperforming the human NeuroSynth map (Spearman’s *r* = 0.51, *p*_*moran*_ *<* 0.001). Notably, the LLM map remains significantly correlated with the map of macaque working memory from Leavitt, even after partialling out the map of human working memory from NeuroSynth (Spearman’s *r* = 0.45, *p*_*moran*_ *<* 0.001). The same results are replicated upon using NeuroQuery instead of NeuroSynth (Leavitt-NeuroQuery correlation: Spearman’s *r* = 0.39; LLM-Leavitt correlation after partialling out the NeuroQuery map: *r* = 0.51, *p*_*moran*_ *<* 0.001). In other words, the summary of the literature produced by LLMs is species-specific, and it goes beyond meta-analyses of the human neuroimaging literature for representing the available evidence about macaque working memory. We also demonstrate a concrete application of cognitive decoding across all three species, for the distribution of the dopamine D1 receptor, obtained from in vivo Positron Emission Tomography in human (88), from in vitro autoradiography in the macaque (59), and from immunohistochemistry in the mouse (89) (Fig. S12). Up to this point, this analysis was only possible in human. By extending to macaque and mouse, we converge towards an evolutionarily conserved association with the neural circuit for ‘reward’ (Fig. S12). Collectively, application of AI experts to mouse and macaque literature is encouraging, demonstrating the added value of an LLM-powered approach for automated data integration that complements empirical databases.

### AI experts’ neuroscientific competence recapitulates functional similarity and structure-function relationships across species

So far, we used LLMs as automated neuroscience experts to map functions to brain areas, but this approach does not take into account how the network structure of the brain supports coordinated and distributed interactions (8, 55, 90, 91). To quantify similarity of regions’ functional profiles in an unbiased fashion, we ask automated experts to list the top *N* functions of each brain region (e.g. top 5). The resulting string of text is turned into a text embedding using the OpenAI API (effectively, the reverse of the process by which LLMs generate human-readable text as output). The embedding is a high-dimensional (here, 3072-dimensional) vector representation, such that text strings with more similar meaning will have closer positions in the highdimensional space of the linguistic embedding. Therefore we can use a measure of vector similarity in embedding space (e.g., correlation or cosine similarity) to quantify the similarity between regions’ functional profiles, as determined by AI experts. For each pair of regions, we generate a region×region matrix of functional similarity. This approach provides a quantitative way to ask whether, according to the literature available to LLMs, two regions perform similar functions. Fig. 3a shows this LLM functional similarity matrix. The matrix clearly encodes non-trivial network structure, which we explore next.

**Figure 3.**
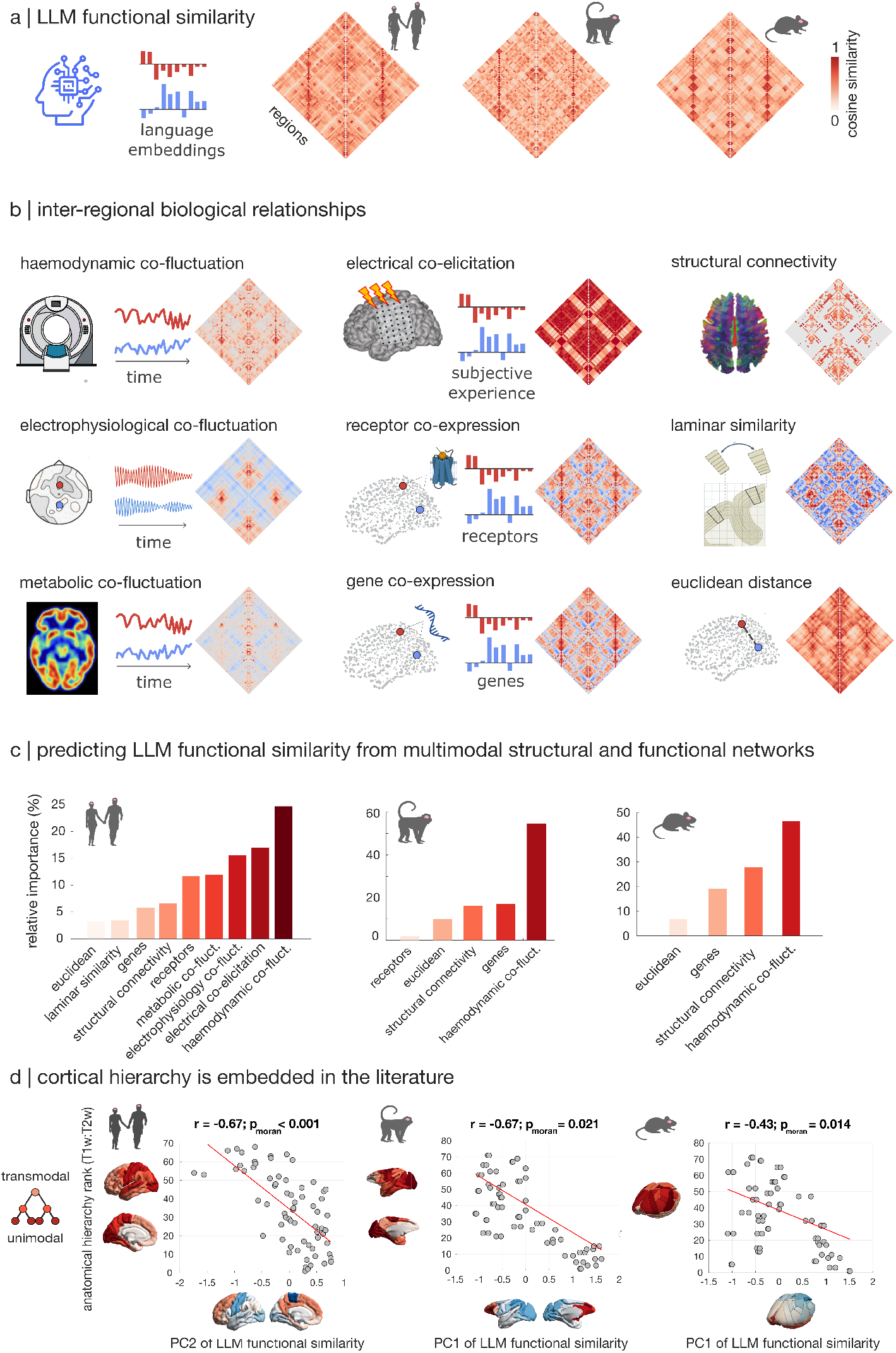
LLMs’ quantification of functional similarity recapitulates structural and functional interactions across species. **(a)** We obtain functional similarity from LLMs as the inter-regional cosine similarity of language embeddings when asked to describe the function of each region. **(b)** Multimodal quantifications of inter-regional interactions for the human brain. Functional interactions can be quantified as co-fluctuation (correlation over time) of spontaneous activity from fMRI haemodynamics (‘functional connectivity’), or electrophysiology from MEG, or metabolic activity from dynamic FDG-PET; or as similarity of subjective experiences upon intracranial electrical stimulation. Anatomical relationships can be quantified as similar patterns of gene and receptor expression, similar cytoarchitecture across layers, presence of direct white matter tracts between regions, or physical proximity in space. See Fig. S13 for corresponding networks available in macaque and mouse. **(c)** Dominance analysis reveals that functional interactions best account for LLM-derived functional similarity across species. Dominance analysis distributes the fit (variance explained) of a multiple regression model across predictors, such that the contribution of each predictor can be assessed and compared to other predictors, producing a relative importance score that reflects the proportion of the variance jointly explained by all predictors, that can be attributed to each predictor. A permutation-based null model (Methods) returns *p* = 0.025 for the probability under pure chance that in each of the three species, the top-ranked predictor will be a functional network. **(d)** We estimate the hierarchical position of each region using the T1w:T2w ratio, an in vivo MRI proxy of intracortical myelin (lower intracortical myelin indicates higher position on the hierarchy). In each species, we find a significant correlation with one of the two principal components of LLM-derived functional similarity (note that PC sign is arbitrary). Significance of correlations is assessed against a null distribution of spatial autocorrelation-preserving surrogate maps. See Fig. S14 for the corresponding null distributions. Brain maps are displayed on the lateral and (for human and macaque) medial surface of the species-specific left cortical hemisphere.

Similarity of function can arise from many forms of interaction between brain regions that can be quantified across multiple spatial and temporal scales using imaging, recording and stimulation (92–94). These include (i) similarity of spontaneous fMRI activity (haemodynamic co-fluctuation, also known as resting-state functional connectivity) (95), (ii) similarity of spontaneous magneto-encephalographic signals (electrophysiological co-fluctuation) (96–99), (iii) similarity of spontaneous FDG-PET activity (metabolic co-fluctuation) (100), and (iv) similarity of subjective experience upon electrical stimulation (electrical co-elicitation) (56) (Fig. 3b). The underlying principle is that if two regions have the propensity to activate and de-activate synchronously, it may be because they perform similar roles in the functioning of the brain (101). Ultimately, whether two brain regions perform similar or different function must arise from anatomy: their respective molecular and cytoarchitectonic make-up, and how they are physically connected with each other and with the rest of the brain (8, 92). We therefore also consider networks of (v) receptor co-expression (correlation across 19 in vivo PET maps) (55, 88); (vi) gene co-expression (correlation of transcriptomic fingerprints) (55, 102, 103); (vii) structural connectivity from diffusion MRI tractography (104); and (viii) laminar similarity (similarity of histologically derived cellular distributions across the cortical layers) (55, 105) (Fig. 3b). For completeness, we also consider (ix) the Euclidean distance between regions as an additional predictor, because the brain’s spatial embedding means that nearby regions may be more likely to perform similar functions than far-away ones (92).

To what extent does the LLMs’ quantification of functional similarity reflect other multimodal forms of interaction between brain regions? To address this question we use dominance analysis (see *Methods*; (106)), which quantifies the relative contribution of each network of inter-regional interaction for predicting the automated expert consensus of functional similarity. We repeat this analysis in macaque and mouse, using fMRI co-fluctuation (64, 78), structural connectivity (62, 107), gene co-expression (60, 61, 63), Euclidean distance, and receptor co-expression (macaque only; (59)) (Fig. S13).

Consistently across human, macaque, and mouse, we find that the fMRI-derived networks of spontaneous haemodynamic co-fluctuation display the highest relative importance for predicting similarity of function across the literature. They outperform anatomical connectivity, spatial proximity, and regions’ molecular similarity (Fig. 3c). In human, where multiple functional networks are available – fMRI co-fluctuation, co-elicitation from electrical stimulation, and electrophysiological and metabolic co-fluctuation – we find that they all rank above the anatomical and molecular ones. Thus, using LLM-based automated summary of the literature allows us to extend for the first time the human results of Smith et al. (101), Crossley et al. (108), Toro et al. (109) to macaque and mouse, revealing that synchronous haemodynamic activity predicts similarity of function. Crucially, these analyses would not be possible through currently available meta-analytic engines, which exclusively cover the human literature. Altogether, automated expert consensus of regions’ functional similarity from the literature is not simply redundant with meta-analytic similarity, but rather provides a valuable complement to the growing compendium of perspectives on inter-regional interactions.

### AI experts discover the latent hierarchical organisation of mammalian cortex

So far, we found that LLMs can recover biologically realistic brain topographies and networks. Does their internal representation also encode other, more complex features of mammalian brain organization that are latent in the literature? One such prominent feature is the hierarchical organization of the mammalian cortex, from unimodal cortices at one end, to transmodal association cortices at the apex (3, 61, 65, 110, 111).

To address this question, we estimate the hierarchical position of individual regions using the T1w:T2w ratio, an in vivo MRI proxy of intracortical myelin (3, 65, 110, 111). We then spatially correlate this empirical estimate of anatomical hierarchy with the principal axes of variation in the LLM functional similarity matrix, obtained from principal components analysis. In each of the human, macaque, and mouse, we find significant spatial correlations between the empirical anatomical hierarchy, and the putative LLM hierarchy (PC2 for human, and PC1 for macaque and mouse; Fig. 3d), suggesting that the LLM experts can discover low-dimensional latent features of brain organization in each species. This is particularly noteworthy because the LLMs are never asked about “hierarchy”, and are only ever queried about one region at a time, yet a distributed hierarchical organisation spontaneously emerges. Altogether, use of LLMs to perform automated structure-function mapping opens the path to comparative discoveries across species and across modalities.

### AI experts recapitulate effects of direct intracranial stimulation beyond meta-analytic associations

Do LLM-derived maps merely reflect meta-analytic statistical associations between cognitive functions and brain regions, or can they draw on a broader evidence-base to summarise the cognitive effects induced by direct experimental manipulations on brain regions? To address this question, we analyze a dataset of direct intracranial electrical stimulation (iES) in surgical patients (*N* = 1, 537 cortical sites in 67 participants) (56). Briefly, participants self-report their subjective experience upon direct electrical stimulation at different cortical locations. Integration of the first-person reports yields probabilistic maps of how direct stimulation elicits broad categories of cognitive processes, including ‘emotion’, ‘language’, ‘memory’, and ‘vision’, which we can use to benchmark the corresponding LLM-derived maps. Fig. 4b shows the iES-derived maps, alongside their spatial correlations with LLM-derived maps (Fig. 4c) and NeuroSynth meta-analytic maps (Fig. 4a). Two broad observations emerge. On one hand, LLM maps are positively and significantly correlated with all iES maps. On the other hand, the correlations between iES maps and LLM-generated maps are all greater in magnitude than the correlations between iES maps and NeuroSynth maps. The same results are replicated when using Neuro-Query instead of Neurosynth (Fig. S16). This key observation demonstrates that automated experts do not simply return existing brain imaging meta-analyses: in such case, their correspondence with iES maps should be on par with NeuroSynth/NeuroQuery. Instead, we provide evidence that in their capacity as neuroscience experts, LLMs consistently go beyond current correlation-based meta-analyses to embody additional insights from the literature on lesions and stimulation, about how individual regions shape ongoing cognition.

**Figure 4.**
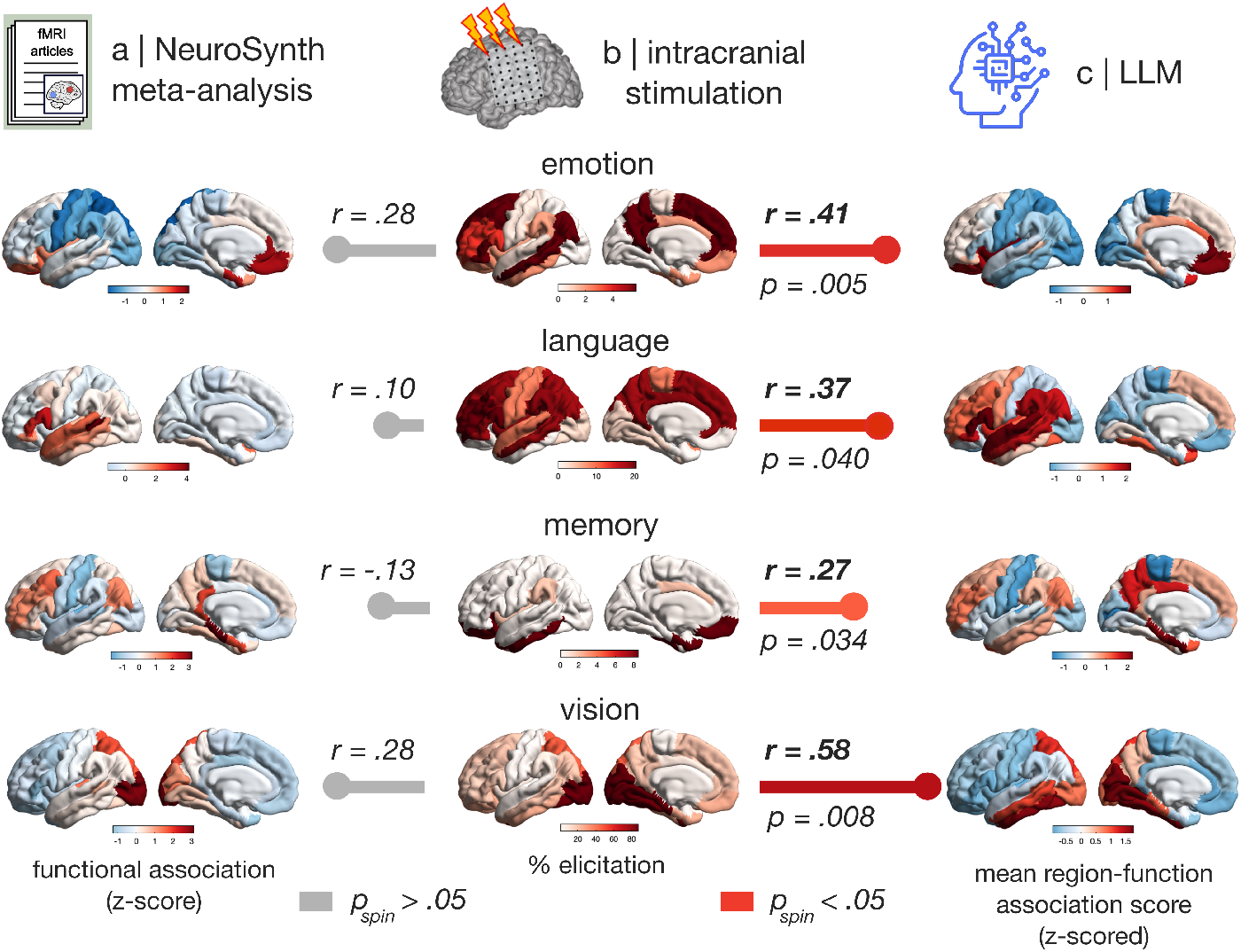
LLM-derived maps of region-function association recapitulate circuits obtained from direct intracranial stimulation. **(a)** NeuroSynth meta-analytic maps (regional z-score from association test; see Methods and (12)). **(b)** Prevalence of subjective experience pertaining to different domains of cognition, elicited by direct intracranial electrical stimulation, for regions belonging to each resting-state network (56). **(c)** LLM-generated maps of region-function association (z-scored across regions within each LLM, then averaged across LLMs; see Methods). Horizontal bars indicate the Spearman correlation between the brain maps displayed at either side, with length proportional to the correlation magnitude; grey bars indicate that the correlation is non-significant (*p >* 0.05). Significance of correlations is assessed against a population of null maps with preserved spatial autocorrelation. See Fig. S15 for the corresponding null distributions and all p-values, including non-significant ones. Brain maps are displayed on the lateral and medial surface of the left cortical hemisphere. See Fig. S16 for replication with NeuroQuery.

### Using AI experts to generate summary maps of brain disease

Up to this point, we used LLMs as AI experts to generate brain maps summarising healthy brain function. But evidence aggregation also provides an opportunity to obtain insight about dysfunction. In psychiatry, intense effort has been directed to understand the extent to which different brain regions are involved in specific syndromes, using numerous imaging, recording and stimulation methods. While individual techniques are increasingly powerful, psychiatric syndromes are increasingly conceptualized as multifaceted diseases that are expressed at multiple levels of brain organization (112). Here we explore whether AI experts, taking a broader view based on the large neuroscience literature included in their training corpora, may generate biologically insightful disease-specific topographic maps.

We systematically query LLMs for their neuroscientific expertise about each region’s involvement in five psychiatric diseases: autism, bipolar disorder, attention deficit hyperactivity disorder (ADHD), major depressive disorder, and schizophrenia. As before, the approach generates disorder-specific consensus maps of regional disease involvement, based on the extensive literature included in the training corpus of each LLM (Fig. 5a). At first blush, the maps recapitulate hot spots well known to human experts; for example, the involvement of subgenual anterior cingulate in depression (57, 113, 114), or the involvement of dorsolateral prefrontal cortex in schizophrenia (115). To contextualize the potential utility of these maps, we contrast them with two other sets of maps, one approximating a “gold standard” and the other approximating the current “field standard”. First, for each disease, we derive regional density of expression of GWAS-derived risk genes in the normative sample of the Allen human Brain Atlas (102) (Fig. 5a, middle column; see *Methods*). Transcriptomic patterns of genes associated with risk for a disease thus provide a molecular signature of the disease (112). Second, for each disease we derive meta-analytic maps of sMRI-estimated cortical thickness abnormality from the global ENIGMA (Enhancing Neuro Imaging Genetics through Meta-Analysis) consortium (Fig. 5a, left column) (14, 17, 18). The regional magnitude of case-control changes in cortical thickness provides an anatomical measure of each region’s involvement in each disorder. We find that all LLM maps exhibit significant spatial overlap with maps of risk-gene expression. In contrast, no significant correlation is observed between the maps of risk-gene expression density, and the corresponding ENIGMA meta-analytic maps of anatomical disease involvement from cortical morphometry. In addition, the magnitude of correlation between risk-gene expression maps is greater with every single LLM-generated map compared with ENIGMA maps. Replicating the analysis with the ENIGMA maps of case-control effect sizes, rather than their magnitude, changes the sign of some ENIGMA-GWAS correlations but produces otherwise virtually identical results, with the sole exception of depression which narrowly passes the threshold for significance (Fig. S20). Note that our interest is in the strength and significance of the statistical spatial association between disease-related maps, rather than the direction (sign) of the association. This is because the transcriptomic maps represent normative adult expression of GWAS-implicated genes. As such, they indicate whether each region’s expression is above or below the mean, rather than indicating disease-related differential expression or whether individual genes are up- or downregulated in patients. As one prominent example, dysregulation or duplication of autism-risk genes can equally produce autism-related developmental syndromes (116–118). A statistically significant association between transcriptomic maps and ENIGMA or LLM-generated maps therefore indicates the existence of a relationship that is unlikely to be due to random chance or spatial autocorrelation. While these statistical associations do not imply causal mechanisms, they are consistent with the neuroanatomical expression of key vulnerability genes associated with these conditions. Altogether, automated experts’ summary of the literature about regional involvement with specific neuropsychiatric disorders is more sensitive to regional genetic risk than manual meta-analysis of cortical morphometry.

**Figure 5.**
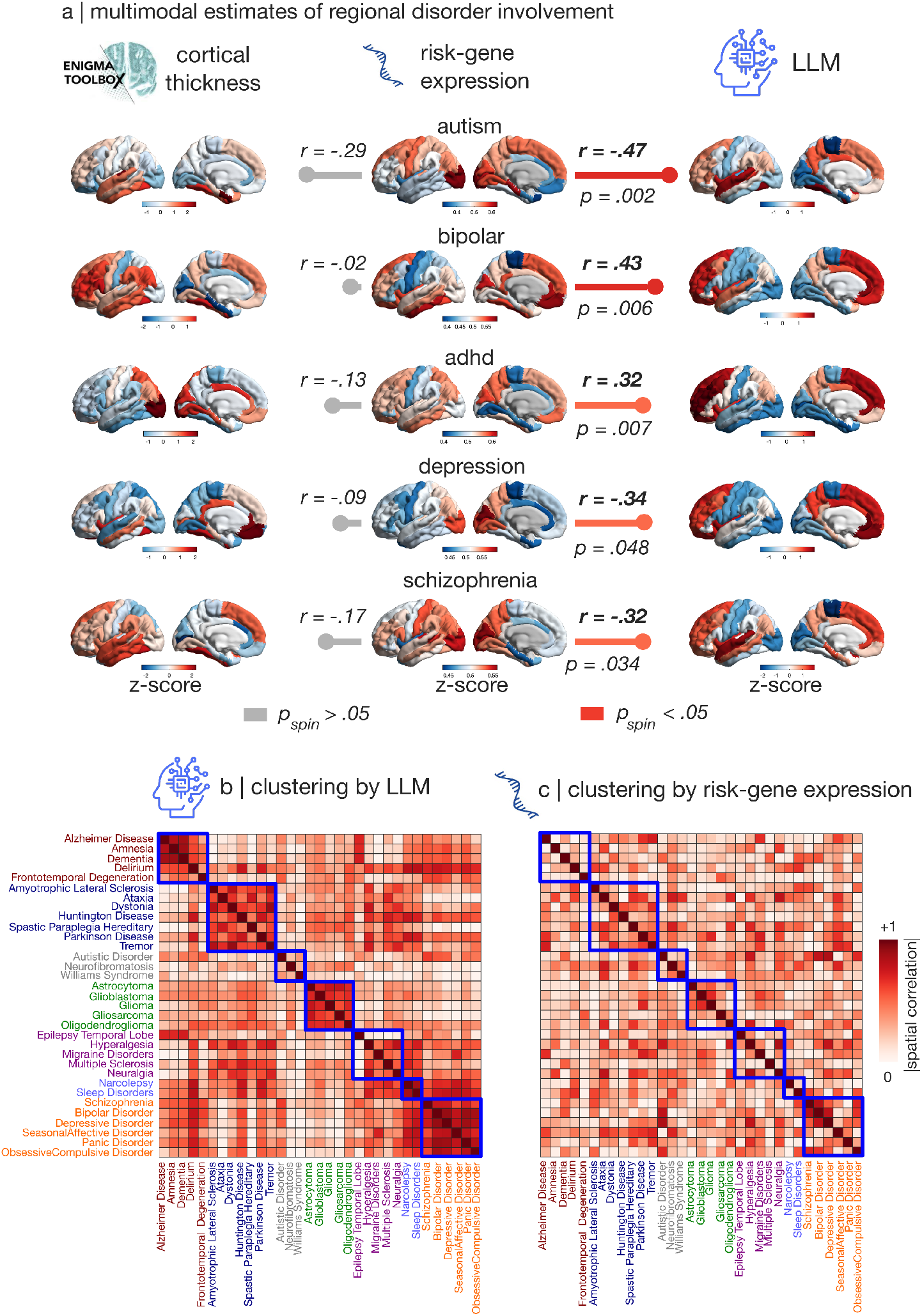
LLMs recapitulate brain circuits of cognitive dysfunction from risk genes and atrophy. **(a)** Multi-modal regional disorder involvement from ENIGMA meta-analysis of cortical thickness (left); from transcriptomic expression of risk-genes identified by genome-wide association studies (GWAS) (middle); and from consensus of AI experts (right). Horizontal bars indicate the Spearman correlation between the brain maps displayed at either side, with length proportional to the correlation magnitude; grey bars indicate that the correlation is non-significant (*p >* 0.05). Significance of correlations is assessed against null distributions of surrogate brain maps with preserved spatial autocorrelation. See Fig. S19 for the corresponding null distributions. All maps are z-scored across regions. For transcriptomic maps, a negative z-score therefore indicates below-average expression of the corresponding genes. For the ENIGMA and LLM maps, a negative z-score indicates below-average anatomical (for ENIGMA) and functional (for LLMs) involvement in the disorder, respectively. See Fig. S20 for replication using ENIGMA case-control effect sizes instead of their magnitude. Maps are displayed on the lateral and medial surface of the left cortical hemisphere. **(b**,**c)** Similarity (magnitude of spatial correlation) between disease maps from LLM-quantified regional involvement (b) aligns with symptom dimensions significantly better than similarity from transcriptomically derived maps of molecular risk (c). Modular structure is based on assignment into 7 disease categories from the International Classification of Diseases manual: Neurocognitive disorders and Dementias; Neurodegenerative and movement disorders; Neurodevelopmental disorders; Brain tumors; Neurological disorders; Sleep and Wake disorders; and Mood and Behaviour disorders. The difference in modular structure between LLM similarity and molecular similarity is significantly greater than chance, as assessed using a label-permutation null model whereby the difference in modularity score is recomputed after randomly rearranging the matrix (5,000 nulls). (Fig. S21).

Taking this idea further, we test the capacity of automated experts to recapitulate shared relationships and comorbidities among brain diseases. We first compile risk genes across multiple neurological, psychiatric, neurodevelopmental, sleep, and tumor-related syndromes, and generate brain-wide transcriptomic maps for each (112). Genes associated with risk for brain disease exhibit characteristic expression patterns. Fig. 5c shows the patterns of similarity (correlations) among these diseases based on their cortical risk gene expression. Fig. 5b instead shows the patterns of similarity among LLM-generated brain maps for the same diseases. Both heatmaps are organized according to the International Classification of Diseases (ICD-11), a global standard diagnostic manual published by the World Health Organization (119). To ask which of the two matrices (risk genes or LLM) better captures ICD classifications, we compute the modularity of each with respect to ICD categories. The LLM matrix is significantly more modular compared to the risk gene matrix (modularity Δ = 0.06, *p <* 0.001; Fig. S21; see Methods), suggesting that LLMs—even when only asked about a single region and a single disease at a time—find that diseases from the same family display similar cortical topography. Interestingly, AI experts also identify similar functional involvement between diseases that are not part of the same category but are nevertheless related, such as for dementia and delirium with many psychiatric disorders including schizophrenia (originally termed ‘dementia precox’ (120)). This set of associations does not emerge from patterns of risk-gene expression. Likewise, a relationship between sleep disturbances and psychiatric disorders is increasingly recognised (121, 122), as is a link between epilepsy and dementia (123, 124): both are correctly identified by AI experts, but not by similarity of risk-gene expression (Fig. 5b,c). Crucially, these links between disorders are an emergent property: the automated experts are never queried about more than one disorder at a time, and never asked about comorbidities, yet similar brain maps are produced for disorders that turn out to be co-morbid. The nosological insight of AI experts may potentially stem from the fact that they are directly queried about cognitive function (rather than genetic risk), which is presumably closer to the behavioural and cognitive symptoms that are used to classify each disease.

### AI experts recapitulate lesion-based circuits of cognitive dysfunction

Corroborating the causal evidence from direct intracranial stimulation, we demonstrate that LLM-generated maps of region-disorder association recapitulate causal insight about regional dysfunction from lesion network mapping (5, 6, 125) (for detailed discussion of this technique, we refer the reader to (126–129)). Namely, we use a psychosis circuit, derived from regions whose lesion induces psychosis symptoms (58)(*N* = 153 lesion cases); and a depression circuit, derived from the convergence of regions whose lesion induces depressive symptoms, and whose brain stimulation (using DBS or TMS) alleviates depressive symptoms (57)(14 studies; *N* = 461 lesion cases and *N* = 251 stimulation cases). We compare them with the LLM-derived and ENIGMA-derived maps of schizophrenia and depression, respectively; but also with the maps of bipolar disorder, which can involve both depressive and psychotic symptoms. We find that the bipolar map from cortical morphometry exclusively tracks the lesion circuit for depression but not psychosis. The opposite is observed for the bipolar map generated by AI experts, which unlike the ENIGMA map tracks risk-gene expression ( Fig. 5a) and which tracks the lesion network circuit for psychosis but not depression (Fig. 6) - possibly indicating a dissociation between anatomical and functional underpinnings of the depressive and manic phases of this complex disorder. For all other comparisons, the disease maps generated by AI experts exhibit stronger (and significant) associations with the causal maps from lesion network mapping (Fig. 6). Notably, we find that even though the LLM-derived maps of depression and schizophrenia do not correlate with the corresponding risk-gene maps, they do track causal circuits obtained from lesions and stimulation, which arguably offer a more direct link between regions and their functional roles. Collectively, these results demonstrate that automated neuroscience experts are sensitive to both (more distal) genetic and (more proximal) lesion circuit manifestations of cognitive dysfunction.

**Figure 6.**
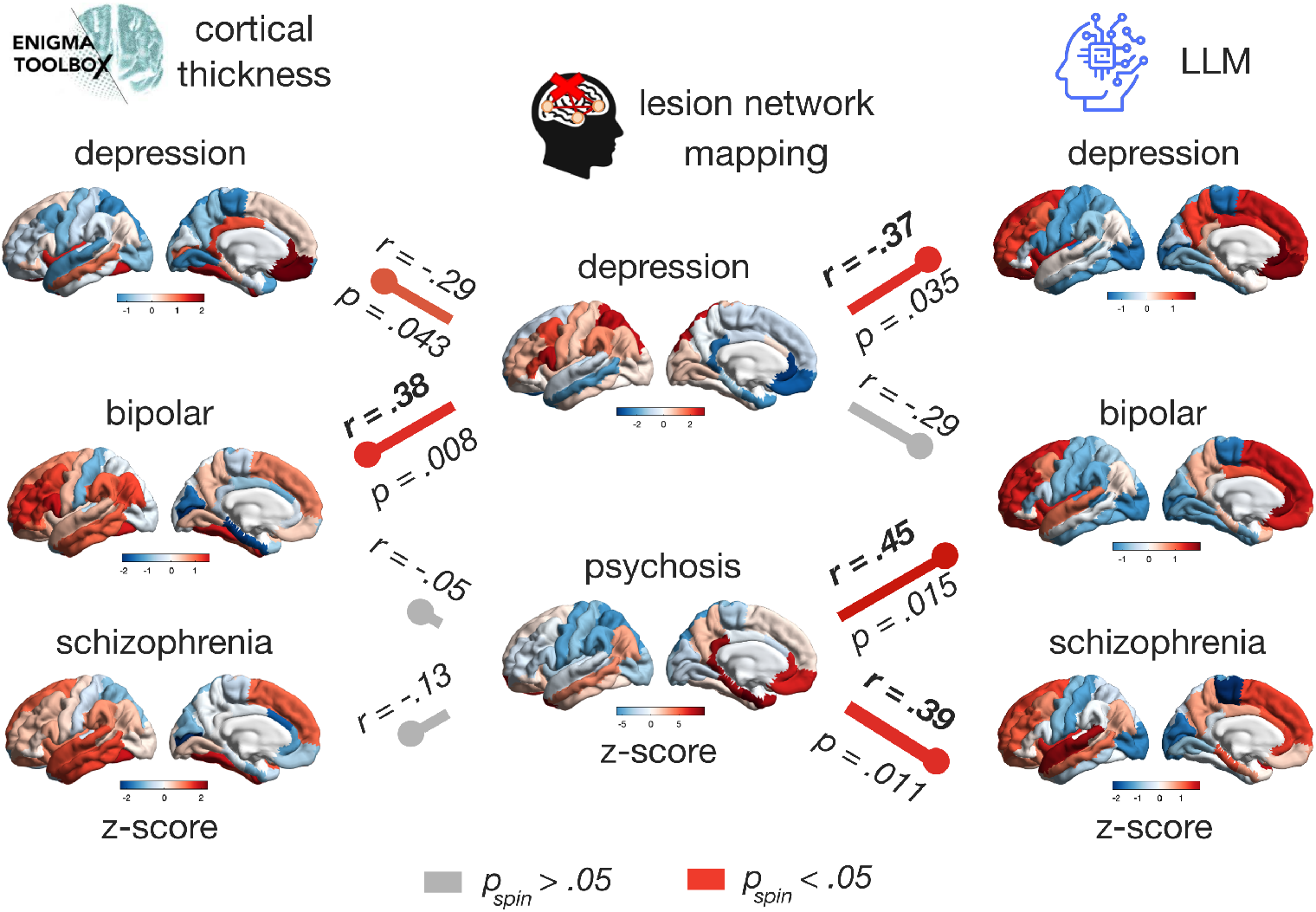
LLMs recapitulate circuits of cognitive dysfunction from lesion network mapping. We use two maps of lesion circuits: (1) a psychosis circuit, derived from regions whose lesion induces psychosis symptoms (58)(*N* = 153 lesion cases), and (2) a depression circuit, derived from the convergence of regions whose lesion induces depressive symptoms, and whose brain stimulation (using DBS or TMS) alleviates depressive symptoms (57)(14 studies; *N* = 461 lesion cases and *N* = 251 stimulation cases). In other words, the two maps provide a causal mapping from regions to symptoms (5). We compare these lesion maps (middle column) with the ENIGMA (left-most column) and LLM-derived maps (right-most column) for depression and schizophrenia, respectively, as well as bipolar disorder, which involves symptoms of both. Except for the bipolar-depression link, the disease maps generated by AI experts consistently exhibit stronger (and significant) associations with the causal maps from lesion network mapping. Colored bars indicate the Spearman correlation between the brain maps displayed at either side, with length proportional to the correlation magnitude; grey bars indicate that the correlation is non-significant (*p >* 0.05). Significance of correlations is assessed against a population of null maps with preserved spatial autocorrelation. See Fig. S17 for the corresponding null distributions, and Fig. S18 for replication using ENIGMA case-control effect sizes instead of their magnitude. All maps are z-scored across regions. For the ENIGMA and LLM maps, a negative z-score indicates below-average anatomical (for ENIGMA) and functional (for LLMs) involvement in the disorder, respectively. Maps are displayed on the lateral and medial surface of the left cortical hemisphere.

### Opening new research directions with automated neuroscientists

What new investigations are possible with this analytic framework, that were not available before? As a final step, we showcase the capacity of LLMs to take the results of two recent multi-modal integration studies that up to this point were only possible in humans (130, 131), and extrapolate them to other model organisms. We use partial least squares analysis (PLS) to integrate LLM-generated maps for 22 representative cognitive operations (corresponding to the most fine-grained subdivision of the cognitive ontology of Beam et al. (69)), and expression patterns of 81 brain-related genes (65, 132) that have available orthologs in human (from microarray (102)), macaque (from stereo-seq (60, 61)) and mouse (from in situ hybridization (63)). In each of human, macaque, and mouse, we consistently find a statistically significant latent variable of gene-cognition association (Fig. S22), replicating the human results originally obtained by Hansen et al. (131) using NeuroSynth, and extending them to two additional species (Fig. 7). In all three species, the main latent variable of gene-cognition association recapitulates the one obtained by (131) in human: it distinguishes LLM-derived emotion- and memory-related cognitive operations from sensory and motor ones. We also formally demonstrate that in each species, LLM-derived cognitive loadings exhibit strong and significant correlations with those obtained for the human brain by meta-analytic maps as per (131) (human: Spearman *r* = 0.75, *p <* 0.001; macaque: Spearman *r* = 0.69, *p <* 0.001; mouse: Spearman *r* = 0.60, *p* = 0.004; Fig. 7). Likewise, in each species we observe a significant association between the gene loadings produced by PLS against LLM-derived maps, and the gene loadings obtained for the human brain by PLS against meta-analytic maps as per (131), even though the original human gene expression was from microarray, whereas the macaque gene expression is from stereoseq (Spearman *r* = 0.57; *p <* 0.001) and the mouse gene expression is from in situ hybridization (Spearman *r* = 0.53, *p <* 0.001) (Fig. 7). Altogether, three separate data-driven analyses in three separate species, each with a different technique for spatial transcriptomics, converge to identify a conserved molecular architecture for cognition in the mammalian brain.

**Figure 7.**
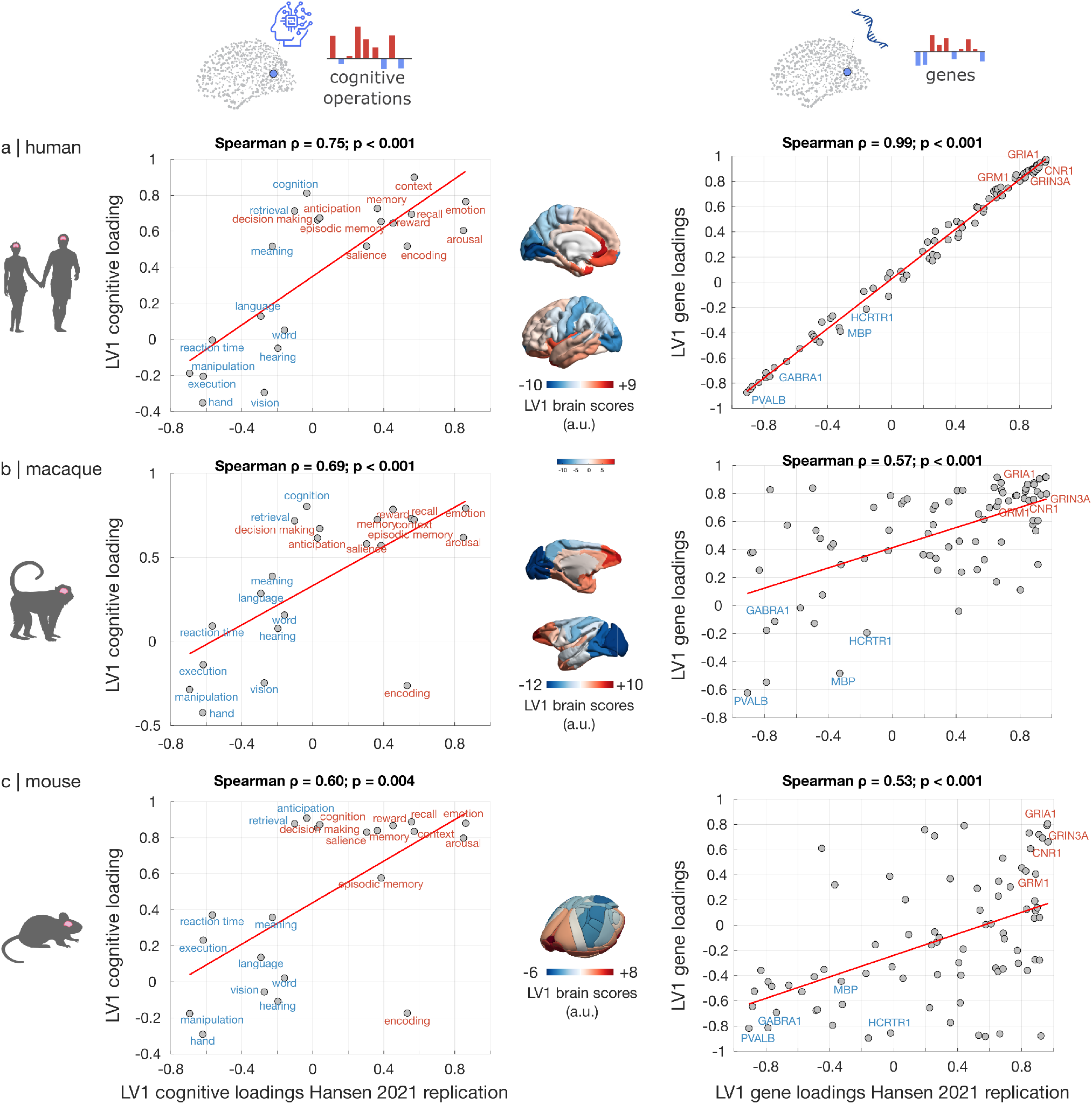
AI-driven discovery of consistent gene-cognition associations across mammalian brains. **a** For human brain, the LLM-derived maps of brain function provide essentially identical insights as NeuroSynth, about the relationship between human cognition and the human brain’s molecular architecture. Despite using only a subset of genes and a subset of terms, and despite using cognitive maps from LLMs rather than from NeuroSynth, we successfully replicate the results of Hansen et al. (131) to a high degree. The main latent variable of gene-cognition association from Partial Least Squares analysis (LV1) displays the same anteriorposterior brain pattern, with both term loadings (left column) and gene loadings (right column) onto this latent variable being highly correlated with the corresponding loadings obtained from linking human gene expression with meta-analytic maps as per (131) (terms: Spearman *r* = 0.75, *p <* 0.001; genes: Spearman’s *r* = 0.99, *p <* 0.001). See Fig. S23 for correspondence between PLS LV1 scores. **b** LV1 cognitive loadings and gene loadings obtained from PLS of LLM-derived cognitive maps against stereoseq gene expression in the macaque brain, are significantly correlated with the corresponding loadings obtained from linking human gene expression with meta-analytic maps as per (131) (terms: Spearman *r* = 0.69, *p <* 0.001; genes: Spearman’s *r* = 0.57, *p <* 0.001). **c** LV1 cognitive loadings and gene loadings obtained from PLS of LLM-derived cognitive maps against in situ hybridization gene expression in the mouse brain, are significantly correlated with the corresponding loadings obtained from linking human gene expression with meta-analytic maps as per (131) (terms: Spearman *r* = 0.60*p* = 0.004; genes: Spearman’s *r* = 0.53, *p <* 0.001). In each species, significance of the PLS latent variables is assessed against a population of null maps with preserved spatial autocorrelation (Methods). See Fig. S22 for the corresponding null distributions. Brain maps display the LV1 scores for each species (a.u.; note that sign is arbitrary). Brain maps are displayed on the lateral and (for human and macaque) medial surface of the left cortical hemisphere.

In the Supplementary Results we also extend across species the recent results of Luppi et al. (130). We combine principles of network control theory (133–135) and *ex vivo* tractography in non-human species with our AI-driven cartography of macaque and mouse cognition, to reveal that the architecture of macaque and mouse brain networks is wired to favour energetically efficient transitions between cognitive states, even beyond the effect of geometric embedding (Fig. S24) – thereby generalising to two new species a recent finding that up to this point had only been possible for the human brain, due to its reliance on NeuroSynth (130). Altogether, use of LLMs to perform automated structure-function mapping opens the path to consistent discoveries across species and across modalities, beyond what is possible with the current human-specific meta-analytic frameworks.

### Validation, robustness and sensitivity

To evaluate the robustness and generalisability of our findings, we assess the capacit of AI expertsy to predict unseen manual meta-analyses. We also benchmark LLMs’ meta-analytic performance against cost, across a broader range of 20 proprietary, open-weight, and local LLMs.

### Prediction of unseen meta-analysis

Because previously published NeuroSynth or NeuroQuery maps may be represented in LLM training corpora, we next tested whether LLM-derived maps generalise to evidence unavailable during model training. We used as target an expert-curated meta-analytic map that summarises the results of 39 studies of functional MRI, structural MRI, and [F^18^]FDG-PET in 1,156 patients with disorders of consciousness (136) first released in January 2026, together with LLMs whose knowledge cutoffs preceded its release. Namely, we use `Mistral AI’s mistral-small-3.1-24b-instruct` (cutoff: October 2023), OpenAI’s `GPT-4o-mini` (cutoff: July 2024), DeepSeek’s `DeepSeek-R1` (cutoff: January 2025) (Fig. S25a). We find that the LLM-generated map of ‘consciousness’ significantly correlates with the unseen expert-curated meta-analytic map (Spearman’s *r* = 0.42), outperforming the predictions of NeuroQuery and also NeuroSynth (Fig. S25b). In other words, LLMs are not just returning the results of previous meta-analyses included in their training corpora, but rather actively performing a synthesis of the broader evidence at their disposal, much like a human expert might.

### Benchmarking LLMs’ capacity for neuroscientific meta-analysis as a function of model, cost, and recency

LLMs from different companies and model families are trained on different corpora of data, and also differ in terms of specific architecture (e.g., number of parameters; instruction-tuning; reasoning models; dense models or mixture-of-experts) and training regime - giving rise to potentially valuable diversity among their neuroscientific expertise. Thus, the NeuroNaut framework is designed to give users freedom in the choice of which and how many LLMs to use. Users may visualise an upto-date list of models available on OpenRouter and their price with the command list-models; both proprietary and open-weight LLMs are included, and some can be used free of charge.

We benchmark the capacity of 20 different LLMs to recapitulate brain maps of cognition from NeuroSynth (Fig. S26a and Table 1). In addition to the 3 frontier LLMs used for the main results, we also include additional proprietary models from OpenAI (`GPT-4o-mini and o3`), Anthropic (`Claude-4.6-Sonnet`) and Google (`Gemini-3-flash` and `Gemini-2.5-flash`). We also consider a range of open-weight models from OpenAI (OSS-120b and the smaller OSS-20b, enabling direct size comparison) and Google (`Gemma-4-31b-instruct`) but also Alibaba (`Qwen3-235b-a22b-2507, Qwen3.5-flash, and Qwen-QwQ-32b`); Meta (`llama-4-maverick and llama-3.3-70B`); DeepSeek (`r1` and `r1-distill`); and Mistral (`Mistral-7B`). We further include Mistral’s `Mistral-Small-24B` as a local model, using 4-bit quantization to enable running the model locally on a personal laptop (MacBook Pro M3). These models span over at least two orders of magnitude in size (number of parameters), and over two orders of magnitude in price (from $0.03 to $5.00 per 1M tokens; Table 1). Partial Spearman correlation reveals a significant correlation between model recency and performance (*β* = 0.52, *p* = 0.011; Fig. S26b), beyond which the relationship between performance and cost is non-significant (*β* = 0.33, *p* = 0.085; Fig. S26c). In other words, LLMs keep improving their capacity to summarise the neuroscientific literature about region-function associations, suggesting that if the current trend continues, the current performance may improve even further in the future.

**TABLE 1.**
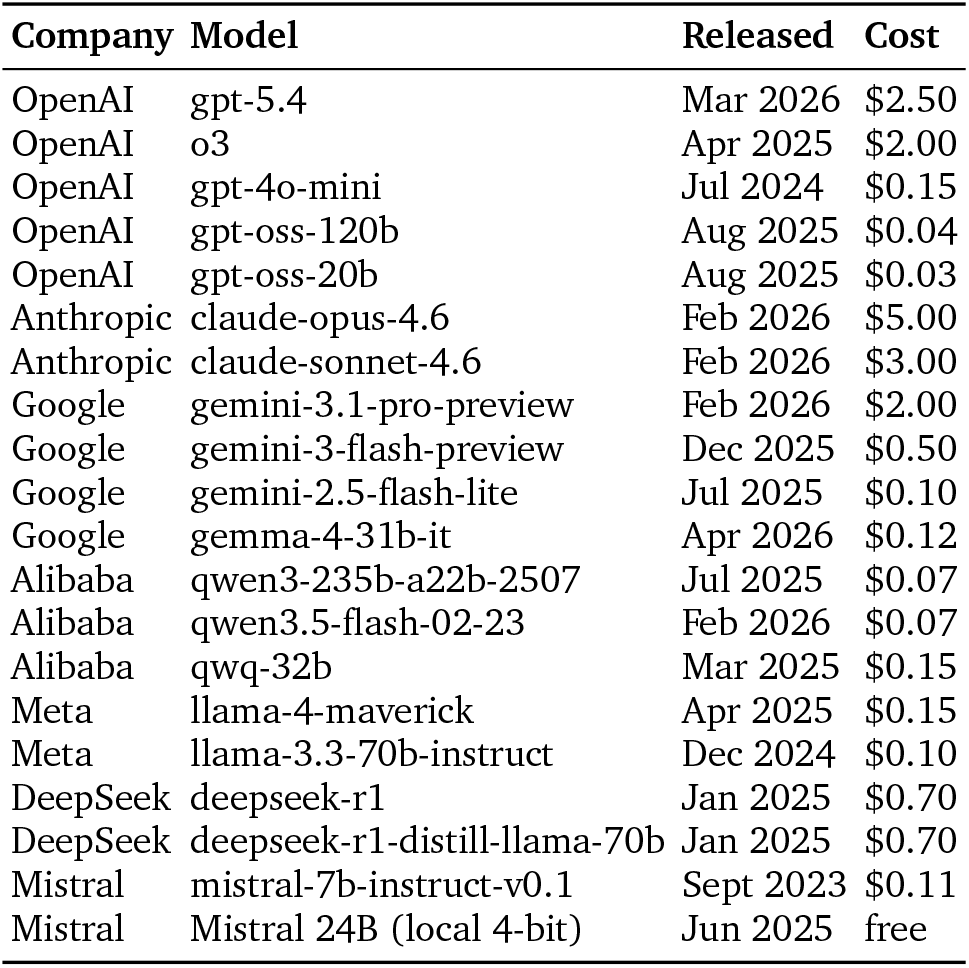
Large language models, their OpenRouter IDs, and their release times, as of June 2026. Cost is expressed as $/1M tokens.

After assessing the performance of individual LLMs, we consider whether performance can be improved by combining multiple AI experts’ opinions, and how performance changes as a function of available budget (price per 1M tokens; Fig. S27). Note that the actual cost of a 68-region brain map is far below the million-token price tag: approximately $0.30 for the combined `Claude-4.6-Opus`, `Gemini 3.1 Pro` and `GPT-5.4` used for our main analyses. We find that a plateau in performance is reached already at a budget of $0.50 with `Gemini-3-flash`, but the best performance is obtained by aggregating the expert judgments of `Claude-4.6`-Opus with `Gemini 3.1 Pro` and `GPT-5.4` (i.e., the combination of top-tier models used in our main analyses; Fig. S27). The neuroscientific expertise exhibited by frontier models of the GPT-5, Claude-4, and Gemini-3-Pro families is consistent with recent demonstration of their top-level performance for biological and medical domain-knowledge (47, 74, 75).

Crucially, highly consistent results are also obtained for under $0.01 per map, using three frontier openweight models: `Qwen3-235b` (37), `OSS-120b` (54), and `Gemma4-31b`. These open models successfully replicate our main results (Fig. S2, Fig. S28, and Fig. S29). Altogether, we find that aggregating top-tier models can further improve their neuroscientific expertise beyond the single best among them; and even though a gap between proprietary and open-weight models remains at the very top end, open-weight LLMs can achieve a 10-to-100-fold reduction in cost, for less than 10% drop in performance, ensuring that the NeuroNaut framework for LLM-based cognitive cartography is widely accessible beyond reliance on any one model or AI company.

### Additional robustness and sensitivity

Through extensive additional analyses we also establish robustness to elicitation method (one region at a time, or pairwise rankings), model family (proprietary or open-weight), choice of parcellation, and test-retest reliability. We demonstrate that region-function association scores closely agree with formal probabilities obtained from pairwise Bradley–Terry comparisons, for a fraction of the cost. We also show that proprietary and open-weight models produce highly concordant maps. We also show high consistency across different LLMs, and high test-retest across multiple API calls of the same LLM. Automated neuroscience experts can successfully identify different names for the same cortical territory from the literature, whether based on convention (e.g., ‘V1’) or anatomical (‘pericalcarine’), functional (‘primary visual cortex’), or cytoarchitectonic criteria (‘striate cortex’, ‘Brodmann Area 17’). Consequently, results replicate across anatomical, cytoarchitectonic and multimodal cortical and subcortical parcellations, with resolution spanning from 44 to 360 regions. We implement a tool for LLM-assisted filtering of NeuroSynth abstracts, NeuroSieve (https://github.com/netneurolab/neurosieve), and we use it to demonstrate that the correspondence between LLM-generated and NeuroSynth-generated maps for ‘manipulation’ improves markedly upon refining the corpus aggregated by NeuroSynth to filter out irrelevant abstracts. We also show that the LLM map better matches a map of lesion-induced loss-of-function from the classic work of Wernicke’s disciple Karl Kleist (137, 138). We show how AI experts successfully map a broader repertoire of 123 cognitive operations, exhibiting significant correlations with NeuroSynth for over 100 of them. Finally, we provide additional demonstrations of how AI experts can be used to generalise human-specific findings to new species, replicating the recent control-theoretic analysis of Luppi et al. (130) for the human brain, and extending its results to macaque and mouse brains. Full results are provided in the Supplementary Results.

## DISCUSSION

The goal of cognitive neuroscience is to understand how brain architecture gives rise to cognition, behaviour, and dysfunction. The literature is vast and variegated, encompassing both healthy humans and patients, but also more experimentally accessible animal models such as non-human primates and rodents; and spanning a wide range of methodologies, from different neuroimaging modalities, to lesions and brain stimulation. We developed and made openly available NeuroNaut: a fully automated workflow to summarise the knowledge of the scientific literature embedded in the pre-training corpora of general-purpose large language models, without need for any additional training.

Automated meta-analysis of AI experts goes beyond traditional coordinate-based meta-analysis, extending its reach from a single imaging modality to the broader literature on direct interventions, lesions, and disease. In parallel, use of AI experts to summarise the literature extends to macaque and mouse analytic approaches that were previously the sole domain of human neuroscience. This capability is showcased by generalising to macaque and mouse the previous human-only results from Smith et al. (101) and Toro et al. (109) that haemodynamic FC between regions tracks the similarity of their functional profiles; the efficiency of the brain’s network architecture for promoting cognitive transitions (130); and the molecular signatures of cognition and affect from Hansen et al. (131).

The evaluations performed here suggest that general-purpose pre-trained LLMs have great value as automated experts for summarising the functions of specific regions, and the neural circuits underlying specific cognitive functions. Brain maps of specific cognitive functions assembled through consensus of AI experts bear significant spatial overlap with conventional meta-analytic maps, while also going beyond them, capturing maps of direct electrical stimulation or lesions that are not available from current meta-analytic engines (5, 56, 57). Notably, LLMs are never explicitly asked about relationships between diseases. Yet we found that the brain circuits of disease generated by AI experts spontaneously cluster in accordance with the clinician-defined relationships of comorbidities and shared symptoms between diseases – which is not apparent for corresponding circuits of disease-related gene expression.

This overwhelming utility of AI experts is consistent with the fact that they can draw on large swathes of the scientific literature and they interpret findings similarly to human experts–though it is worth bearing in mind that the literature itself is not free from bias (13, 139), and neither are human experts.

Like other large-scale literature-aggregation methods, LLM-based summaries inherit variation in the quality, recency and reproducibility of the underlying literature. A key advantage of AI experts is that they are not restricted to papers that explicitly report MNI coordinates, and therefore they can draw on a potentially broader evidence-base: not only neuroimaging studies that do not report coordinates or use region names instead, but also other modalities beyond MRI such as classic neuropsychology, which have been foundational in shaping modern brain mapping. Crucially, this methodology allows us to circumvent possible bias of both manual and automated coordinate-based meta-analyses due to lack of reported coordinates in neuroimaging studies (estimated to be around 42.55% in MRI/PET studies), which according to recent evidence might remain unretrievable in about one-third of cases, even when directly contacting the authors (136). Across the present analyses, we assessed convergent validity using independent causal datasets from lesions and stimulation, as well as expert-curated data unpublished at the models’ knowledge cut-off date, and negative controls (telekinesis, telepathy). We also implemented NeuroSieve, to filter out irrelevant uses of polysemous words from the abstracts that NeuroSynth includes. We showed that increasing the quality of NeuroSynth maps brings them into closer alignment with human experts’ expectation, and crucially it also brings them into closer alignment with the LLM-generated maps. Ultimately, automated summary of the literature is only one component of the scientific triangulation strategy, complementing the distinct and fundamental roles of both formal meta-analysis of individually expert-curated studies, and high-quality, theory-driven primary studies.

The NeuroNaut workflow is easy to use, supporting hundreds of popular LLMs – including both proprietary and open-weights models – from a variety of big and small AI companies, based in several countries. Ultimately, differences in cost, performance, and even environmental impact may also need to be factored into one’s choice of model(s) to use. To aid in this choice, we provide benchmarking of model cost, size, and open-weight status against meta-analytic performance for 20 popular LLMs from major AI companies: OpenAI, Anthropic, Google, Meta, Mistral, Alibaba, and DeepSeek. Crucially, we also implemented the ability to use local LLMs, which are open-weight, free of charge, and under full user control.

Future work may further expand on NeuroNaut in several directions. We chose to rely on pre-trained general-purpose LLMs without need for any fine-tuning, to maximise both scalability and the ease and immediacy of use. Indeed, a recent study (51) showed that across all domains of neuroscience tested, off-the-shelf LLMs consistently outperform expert human neuroscientists by a wide margin at identifying the true results of a neuroscientific abstract based on the description of background and methods, achieving 81% correct (vs human experts’ 66% correct) even without any fine-tuning–providing confidence in our own use of the LLMs as ‘automated experts’. Although fine-tuning may further improve performance over the pre-trained baseline of a model (44, 51) (e.g., from 83% to 86% correct for neuroscientific expertise), recent reports indicate that frontier general-purpose LLMs of the GPT-5, Claude-4, and Gemini-3-Pro families (which we used in the present work) comprehensively outperform specialised AI models fine-tuned for clinical expertise across a broad range of medical benchmarks (47, 75). Alternative prompt engineering strategies may also prove valuable (48, 140–142). To extract quantitative data in a systematic fashion we made use of strictly regimented prompts. Our approach therefore does not currently include the opportunity for users to engage in dialogue with the LLMs, challenge them, or pursue clarifying questions –although we do provide the option to obtain a textual justification.

Future extensions could also incorporate retrieval-augmented generation (RAG) (143, 144) to produce references for the provided output, though at the cost of requiring continuous knowledge-base maintenance and additional workflow complexity. Inevitably, different approaches will have their own specific advantages and shortcomings. We seek to facilitate such extensions by making the NeuroNaut code-base available online (https://github.com/andrealuppi/NeuroNaut), by providing an interface for using our tool with local LLMs, and by enabling use of custom prompts beyond the `-default` option. Altogether, by allowing users the freedom to access hundreds of AI experts, we believe that our workflow is well positioned to capitalise on both the rapid pace of AI models’ development, and the continuously expanding knowledge-base on which they are trained by AI companies.

The `NeuroNaut` workflow differs from a classic statistical meta-analysis in the traditional sense of directly aggregating effect sizes from specific individual studies to obtain an overall effect size (21). Rather, the automated consensus of AI experts introduced here offers a quantitative overview of the literature, akin to what may be obtained from aggregating human experts’ opinions (e.g., (145)). Our results suggest that this approach is useful in the sense that AI experts—even though only prompted about individual functions in individual brain regions— can recover additional organizational features, such as inter-regional covariance patterns and hierarchical differentiation.

Although we found broad agreement between the cognitive maps generated by different AI experts, it is important to keep in mind that experts – both AI and human – can be wrong, and expert consensus should not be equated with ground truth, lest it result in undue silencing of alternative perspectives. Like other forms of scientific literature aggregation, the present approach may be susceptible to the literature’s well documented biases, such as confirmation bias and publication bias (12, 13, 139, 146). Additionally, neuroscience presents its own set of systemic biases, including a disproportionate focus on specific brain regions and cognitive tasks (147–150), a limited repertoire of animal models (151– 153), and an uneven sampling across geographic and demographic populations (154). These imbalances in coverage may lead to extrapolations by LLMs in domains where evidence is sparse. It will therefore be increasingly important to both address the biases inherent to our field and interpret the results of LLM meta-analyses considering these biases. Crucially, cases when LLMs’ answers consistently diverge from each other (Fig. S30), from human expectation, or from modality-specific metaanalyses – which our workflow makes easy to discover – offers valuable clues about latent patterns and missing links in the literature, providing the opportunity for automated hypothesis-generation and highlighting follow-up experiments and analyses (139, 155).

At a time of exciting technological, analytical and data sharing advances (1, 78, 156, 157), and an exponentially expanding literature, there is an innate sense of loss and even sadness as one realizes that it is simply impossible to keep abreast of all new knowledge about the brain. The present work showcases how the use of widely accessible AI technologies can summarise knowledge about the brain in a way that is readily useful for generating quantitative insight and asking fundamentally new questions about brain function.

## METHODS

### NeuroNaut: Automated consensus of AI experts using Large Language Models

We introduce `NeuroNaut`: a computational framework for automated meta-analysis of AI experts, summarising knowledge about brain region functions across species using Large Language Models (LLMs). The `NeuroNaut` framework implements three primary analysis modes: (1) hypothesis-driven region-function association, to estimate the likelihood of a specific cognitive function (or dysfunction) being associated with a particular brain region; (2) pairwise rankings, whereby pairs of regions are provided to the model, which decides which of the two is more relevant for a given function, in a binary head-to-head comparison (3) data-driven function listing, to identify the top functional associations for different brain regions. Our tool supports multiple species and brain atlases to enable comprehensive comparative analysis. The present analysis is primarily conducted across three species: human, macaque (*Macaca mulatta*), and mouse (*Mus musculus*), using the Desikan-Killiany (70), Regional Mapping (71), and Allen Mouse Brain Atlas (72), respectively. `NeuroNaut` is freely and openly available online at https://github.com/andrealuppi/NeuroNaut, with full instructions for installation and use. Below we provide an overview of the functionalities used in the present work.

#### Large Language Models

A total of 20 LLMs are employed in this study, encompassing both commercial and open-source models to ensure robust and diverse analytical perspectives (Table 1). The commercial models used for the main analysis are OpenAI’s `GPT-5.4` (33), and `Google’s Gemini 3.1` Pro (35), and Anthropic’s `Claude-4.6-Opus`, representing some of today’s most advanced models. The open-weight models used for replication of the main results are Alibaba’s `Qwen3 235b` (37), OpenAI’s `OSS120b` (54), and Google’s `Gemma4 31b` (36). Note that this list is not exhaustive: as of June 2026, there are over 320 different LLMs available on OpenRouter, including both proprietary and open-weight models, whether free or paid, from a broad range of AI companies based in several countries. Additionally, our framework allows users to add local LLMs (see below) that can operate even on a personal laptop under full user control.

### API-based models

API-based models are accessed via direct API integration with OpenRouter, meaning that users can access hundreds of LLMs from multiple major AI labs using a single API key, instead of requiring a different API key for each provider. Note that this option requires an OpenRouter account, and most models require purchasing OpenRouter credits. All models implement exponential backoff retry logic to handle rate limiting and temporary API failures. Default model parameters are used for all models, with the ‘seed’ parameter pre-set to 42 and temperature set to the most conservative/reproducible value of 0.0 by default to minimise test-retest differences due to stochastic generation, as per recent practice (48). Exploring the effect of temperature on AI experts’ summaries of the literature is outside the scope of this study, but we note that previous literature has shown that even with a temperature of 0.0 “the specific text could vary, from near identity to considerable differences in phrasing” – albeit with conceptually equivalent meaning (48).

### Local models

Local LLMs are also enabled. Such models can be called by using local instead of the provider name, followed by the model name. A desired local LLM must be separately installed by users on their local machine, and the path then added to the NeuroNaut configuration as per the instructions on the GitHub repository (https://github.com/andrealuppi/NeuroNaut). Note that depending on the capacity of the local machine, quantization may be required (158). As proof of concept, here we use a MacBook M3 laptop to run Mistral AI’s `Mistral-Small-24B`, converted to 4-bit MLX format from mistralai/`Mistral-Small-24B-Instruct-2501` using `mlx-lm` version 0.21.0, as available on HuggingFace (https://huggingface.co/mlx-community/Mistral-Small-24B-Instruct-2501-4bit).

#### Region-function association scores

Analysis is performed on a per-region basis: the LLM is queried about a specific region using its anatomical name as per the corresponding atlas. LLMs’ extensive neuroscientific knowledge-base enables them to recognise different names for the same brain region, much like a human expert would (see Supplementary Results for formal demonstration) –thereby enabling systematic mapping of function to anatomy like coordinate-based meta-analytic engines, while overcoming the need for exact coordinates that restricts them to human MRI.

For robustness and sensitivity analysis, in addition to the gyral-based Desikan-Killiany atlas (70) we also replicate the main results using the Cammoun anatomical atlas with cortex and subcortex (159); the Glasser multimodal atlas with two resolutions: 180 unique regions, or 22 cortical territories (160); and the classic cytoarchitectonic atlas of Brodmann (138), which “remains one of the most widely used in neuroscience and continues to serve as a default reference system for reporting the locations of effects in current neuroimaging studies” (160, 161).

For predetermined cognitive functions, LLMs are prompted to estimate the probability that a specific brain region is involved in a specific function (words shown here in [**braces**] are replaced by specific instances based on the other inputs provided):

> “You are an expert in neuroscience literature analysis. Your task is to estimate the probability that the brain region [**hemisphere**] [**region name**] in [**species**] brain is involved in [**function**].
>
> These probabilities should be based on a simulated review of neuroscience literature, reflecting how frequently these functions are associated with this brain region across **peer-reviewed** studies, textbooks, and reputable sources.
>
> ### Guidelines
>
> 1. Consider only [**species**]-specific neuroscience literature. Do **not** use data from other species.
> 2. The probability should be a single decimal number between 0 and 1.
> 3. This number should approximate the relative frequency with which this function is linked to the given brain region in literature.
> 4. **DO NOT** provide explanations, citations, or any extra text — **return only the probability value**.
>
> ### Expected Output Format
>
> 0.XX”

Responses are parsed to extract decimal values of estimated region-function association between 0 and 1, with text parsing algorithms designed to handle various response formats and convert non-numeric responses to standardised probability scores.

#### Pairwise comparisons and Bradley-Terry model

We also make it possible to elicit AI experts’ knowledge using pairwise comparisons rather than by requesting direct numerical probabilities. For a chosen function X, AI experts are asked to decide which of two regions A or B is more relevant. This process is repeated for all possible pairs of regions, across both hemispheres. Users may access this option through the rank-pairs command, whose standardised prompt instructs the LLM as follows:

> “You are an expert in neuroscience literature analysis. Your task is to determine which of two brain regions is more relevant to the function **[**function**]** in the **[**species**]** brain.
>
> Region 1: **[**region 1**]** (Hemisphere: [**hemisphere 1**])
>
> Region 2: **[**region 2**]** (Hemisphere: [hemisphere 2])
>
> Base your assessment on a simulated review of neuroscience literature, reflecting how frequently each region is associated with this function across **peer-reviewed** studies, textbooks, and reputable sources.
>
> ### Guidelines
>
> 1. Consider only **[**species**]**-specific neuroscience literature. Do **not** use data from other species.
> 2. **DO NOT** provide explanations, citations, or any extra text — **return only the number**.
> 3. Return **1** if Region 1 is more relevant, or **2** if Region 2 is more relevant.
>
> ### Expected Output Format
>
> 1 or 2”

The result is a binary tournament matrix *W* of pairwise comparisons, with *W*_*ij*_ = 1 if region *i* is judged more relevant than region *j*, and 0 otherwise (note that no ties are allowed, and the diagonal is ignored). *W* can then be provided as input to a range of preference-modelling algorithms (162), such as simple win-loss ratios, ranking methods based on graph theory such as PageRank, or probabilistic latent-strength models. Here we use the classic Bradley-Terry model (76), which is closely related to the Elo system of chess rankings. The BT model assumes that each item *i* (here, a brain region) is associated with an unobserved (latent) positive strength parameter *π*_*i*_. When item *i* is compared with item *j*, the probability that *i* is preferred to *j* is defined as:

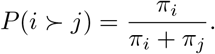

With *θ*_*i*_ = log *π*_*i*_, the model can be re-expressed as:

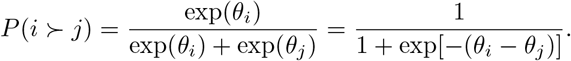

The model is fitted by maximum-likelihood estimation over all observed pairwise outcomes. A small regularisation parameter is added to avoid values going to infinity. The fitted vector (*θ*_1_, …, *θ*_*N*_) provides a single preference score for each region, with larger values indicating regions that are more likely to be preferred across pairwise comparisons.

#### Justifications for region-function association scores

We implement a `–justify` command whereby, alongside assigning a region-function association score to the relevance of region X for function Y, the automated expert will also produce a 1-2 sentence explanation for the score that it provided. This functionality is implemented by modifying the required outputs in the prompt provided to the LLM, as follows:

> “### Expected Output Format
>
> 0.XX | justification
>
> Where justification is a 1-2 sentence explanation (max 200 characters) of your probability estimate.
>
> Use “|” (space-pipe-space) as the separator.
>
> ”

Note that by default, the framework limits the LLM to 256 tokens to keep numerical outputs concise (although this parameter can be overruled by the user if desired). When `–justify` is invoked, the `max-tokens` parameter is automatically increased to 512 tokens to give the model room to return its reasoning.

#### Embedding of top functions

For each brain region, LLMs are queried to identify the top *N* primary functions based on neuroscience literature (here, *N* = 5). The standardised prompt template instructs models to:

> “List out the top 5 functions that region [**region**] in the [**hemisphere side**] hemisphere of the [**species**] brain is involved in. These functions should be based on a simulated review of neuroscience literature, reflecting how frequently these functions are associated with this brain region across peerreviewed studies, textbooks, and reputable sources.”

This option is available through the `top-functions` command. Raw LLM responses undergo systematic cleaning using regular expressions to extract function lists, remove extraneous text, and standardise formatting. Cleaned responses of the top functions are then concatenated into a single text string, which is subsequently converted into semantic embeddings using OpenAI’s text-embedding-3-large model (3,073 dimensions). Local embeddings are also supported as an open-access, free-of-charge alternative. These embeddings serve as high-dimensional representations of each region’s functional profile according to each LLM.

After this, inter-regional similarity is computed using cosine similarity on the semantic embeddings derived from the above queries. For each species and LLM, pairwise cosine similarity matrices are generated to quantify functional similarity between all brain regions. The cosine similarity measure is chosen for its effectiveness in high-dimensional semantic spaces and its interpretability as a measure of functional overlap between regions. Similarity matrices preserve the original atlas ordering of brain regions to maintain neuroanatomical consistency and enable systematic comparison across models. Missing or failed embeddings are handled through exclusion from similarity calculations, with appropriate documentation of data completeness.

### Automated meta-analytic brain maps from NeuroSynth and NeuroQuery

Continuous measures of the association between voxels in MNI-152 standard space and cognitive categories were obtained from NeuroSynth, an automated term-based meta-analytic tool that summarises results from more than 14 000 published human fMRI studies by searching for high-frequency key words (such as “pain” and “attention” terms) that are systematically mentioned in the papers alongside fMRI voxel coordinates (https://github.com/neurosynth/neurosynth, using the volumetric association test maps (12)). This measure of association strength (z-score from a two-way ANOVA testing for the presence of a non-zero association between term use and voxel activation) is the tendency that a given term is reported in the functional neuroimaging study if there is activation observed at a given voxel. In other words, the association test maps tracks whether activation in a voxel occurs more consistently for studies that mention the chosen term than for studies that do not. Note that NeuroSynth does not distinguish between areas that are activated or deactivated in relation to the term of interest, nor the degree of activation, only that certain brain areas are frequently reported in conjunction with certain words. The probabilistic measure reported by NeuroSynth can be interpreted as a quantitative representation of how regional fluctuations in activity are related to psychological processes.

For our main analyses, we used NeuroSynth maps pertaining to eight fundamental domains of cognition, corresponding to the recent data-driven ontology published by Beam et al. (69): emotion, cognitive control, listening, memory, language, vision, reward, and manipulation (69). We also obtained the NeuroSynth map for human ‘working memory’ and parcellated it according to the Regional Mapping atlas of Kotter and Wanke (71), adapted from the macaque to the human brain by Bezgin et al. (87). For the multivariate analysis linking cognition and genes, we used the most fine-grained version of the same data-driven ontology developed by Beam et al. (69), comprising 22 terms. Robustness analysis used a further, broader set of NeuroSynth maps pertaining to 123 cognitive operations selected from the intersection of NeuroSynth and the Cognitive Atlas, a public ontology of cognitive science (148), which includes a comprehensive list of neurocognitive terms. Terms range from umbrella terms (“attention”, “emotion”) to specific cognitive processes (“visual attention”, “episodic memory”), behaviours (“eating”, “sleep”), and emotional states (“fear”, “anxiety”). This set of 123 NeuroSynth maps (Table 2) was chosen because it has been used extensively in previous publications (130, 131, 163, 164). NeuroSynth provides continuous voxelwise maps in MNI-152 standard space, which were parcellated into 68 cortical regions from the Desikan-Killiany anatomical atlas (70) using the `netneurotools` toolbox (https://github.com/netneurolab/netneurotools) (165), by averaging the values of all voxels belonging to the same parcel. Note that no threshold was applied, so as to retain the continuous nature of the map (166). Robustness analyses additionally used the Glasser multimodal atlas with either 180 unique regions or 22 cortical territories (160); the Brainnetome multimodal atlas with 210 cortical regions; the Cammoun anatomical atlas, which corresponds to the Desikan-Killiany cortical parcellation plus 14 subcortical regions (159); and the classic cytoarchitectonic atlas of Brodmann (138, 161).

**TABLE 2.** NeuroSynth terms. List of 123 terms that overlapped between the NeuroSynth database (12) and the Cognitive Atlas (148).

|  |  |  |  |  |
| --- | --- | --- | --- | --- |
| action | adaptation | addiction | anticipation | anxiety |
| arousal | association | attention | autobiographical memory | balance |
| belief | categorization | cognitive control | communication | competition |
| concept | consciousness | consolidation | context | coordination |
| decision | decision making | detection | discrimination | distraction |
| eating | efficiency | effort | emotion | emotion regulation |
| empathy | encoding | episodic memory | expectancy | expertise |
| extinction | face recognition | facial expression | familiarity | fear |
| fixation | focus | gaze | goal | hyperactivity |
| imagery | impulsivity | induction | inference | inhibition |
| insight | integration | intelligence | intention | interference |
| judgment | knowledge | language | language comprehension | learning |
| listening | localization | loss | maintenance | manipulation |
| meaning | memory | memory retrieval | mental imagery | monitoring |
| mood | morphology | motor control | movement | multisensory |
| naming | navigation | object recognition | pain | perception |
| planning | priming | psychosis | reading | reasoning |
| recall | recognition | rehearsal | reinforcement learning | response inhibition |
| response selection | retention | retrieval | reward anticipation | rhythm |
| risk | rule | salience | search | selective attention |
| semantic memory | sentence comprehension | skill | sleep | social cognition |
| spatial attention | speech perception | speech production | strategy | strength |
| stress | sustained attention | task difficulty | thought | uncertainty |
| updating | utility | valence | verbal fluency | visual attention |
| visual perception | word recognition | working memory |  |  |

For our replication and robustness analyses, we also obtained brain maps from NeuroQuery (13). The voxelwise maps in MNI-152 standard space were downloaded from the NeuroQuery website (https://neuroquery.org/) and parcellated using `netneurotools` as described above. NeuroQuery contains terms from the neuroscientific vocabularies that occur in more than 5 out of 10 000 articles, and uses (pre-LLM) natural language processing and semantic smoothing to combine semantically related terms used in the literature, which may be treated as distinct (leading to loss of statistical power) by earlier automated approaches such as NeuroSynth (13). Additionally, whereas NeuroSynth is designed for in-sample statistical inference, NeuroQuery is intended to provide out-of sample prediction: given a query, it uses its semantic model to list related studies and their reported activations (13). Briefly, across studies from the training corpus, the activation of each voxel is regressed on term occurrence frequencies in the corresponding publications. As reported by Dockès et al. (13), after selecting the most predictive terms from its extensive vocabulary, NeuroQuery fits a linear regression to link brain activity with the selected keywords. Therefore, NeuroQuery provides a valuable complement to NeuroSynth as a means of obtaining quantitative summaries of the human cognitive mapping literature.

#### Refined selection of NeuroSynth abstracts

To refine NeuroSynth maps by only retaining relevant uses of the terms of interest, we passed the original abstracts of each study in the NeuroSynth corpus (>14 000 articles) to a large language model not used elsewhere in the study (`Mistral Small 4`) and asked it to re-label, for each term–study pair, the semantic role of the term within the abstract. Each query presented the model with a single term, the study title, and its abstract, under a system prompt that designated the LLM as a cognitive neuroscience expert and required it to assign the term to one of four mutually exclusive roles: ‘primary condition’, ‘contrast condition’, ‘background only’, or ‘absent’ (see the full prompt below for the exact definition) with either high or low confidence. To mitigate the risk of confabulations, the prompt instructed the model to use only information stated in the abstract, to return a lowconfidence rating for ambiguous cases, and justify each label with a short verbatim phrase from the abstract. The system prompt was as follows:

> “You are an expert in cognitive neuroscience. All studies in this database are confirmed neuroimaging studies that report brain activation coordinates. Your sole task: determine what role the provided term plays in this specific study.
>
> - primary condition: the term names the psychological process, cognitive function, behavioural task, or mental state that participants were engaged in or experiencing WHILE being scanned, and to which the reported activation coordinates are attributed.
> - contrast condition: the term describes a baseline or control condition being subtracted from, not the condition of interest.
> - background only: the term appears in abstract but only to frame or motivate the study, not as a scanned condition.
> - absent: the term is not meaningfully present.
>
> Rules: - Use only information present in the abstract. Do *not* infer unstated methods or conditions. - If genuinely ambiguous, choose the most likely role and set confidence to low. - For reasoning, cite the shortest abstract phrase that determined your classification.”

This prompt was iterated separately over every article in the NeuroSynth corpus (*>* 14 000). Inputs were the term of interest, the article’s title, and its abstract:

Term: term

Title: title

Abstract: abstract

Outputs were constrained to a JSON schema reporting (i) the assigned role; (ii) a binary confidence rating (high or low); and (iii) the supporting phrase. To obtain the refined term–study association set, we retained only studies from the NeuroSynth corpus where the term of interest was rated as being the primary condition with high confidence, discarding all contrast, background, absent, and low-confidence classifications. Documented code to implement this method, termed NeuroSieve, is made freely available at https://github.com/netneurolab/neurosieve.

### Cognitive circuits from intracranial electrical stimulation

We analyze a dataset of direct intracranial electrical stimulation (iES) in surgical patients (*N* = 1, 537 cortical sites in 67 participants) (56). Briefly, participants selfreported their subjective experience upon direct electrical stimulation at different cortical locations, which cover all human intrinsic brain networks. Integration of the first-person reports yields probabilistic maps of how direct stimulation elicits broad categories of cognitive processes. As reported in the original publication (56), “eight categories were employed: (1) somatomotor effects, including somatosensations and any observable motor effects, such as muscle twitches or limb movements; (2) visual effects, from simple phosphenes to more complex perturbations of visual perception such as distortion of faces; (3) olfactory effects; (4) vestibular effects, such as feelings of rotation, flotation and acceleration; (5) emotional effects of either positive or negative valence; (6) language effects, including arrest or alteration (for example, slurring) of speech; (7) memory recall; and (8) physiological and interoceptive effects, such as perceived changes in body temperature or heart rate”. Among these, four overlap with the cognitive domains from (69) and were used for the present work: ‘emotion’, ‘language’, ‘memory’, and ‘vision’. Classified results were then pooled to yield an estimate of the prevalence of each effect type for each of the brain’s intrinsic connectivity networks, and the corresponding value was assigned to each region according to its intrinsic network membership.

### Cognitive circuits from lesion studies

Classical neuropsychology identifies the causal roles of brain regions by identifying spatial overlap among lesions that cause the same symptoms. Lesion network mapping (5, 6, 125) extends this principle from overlap in 3D anatomical space, to overlap in ‘network space’: it identifies regions that overlap in terms of their functional connectivity with the sites of lesions that cause the same symptoms. This approach can also be extended to sites of stimulation that ameliorate the same symptoms (57). Here we include publicly available maps from two such studies: (1) a psychosis circuit, derived from the functional connectivity of regions whose lesion induces psychosis symptoms (58)(*N* = 153 lesion cases), available in MNI-152 standard space from NeuroVault at https://neurovault.org/collections/20510; and (2) a depression circuit, derived from the functional convergence of regions whose lesion induces depressive symptoms, and whose brain stimulation (using DBS or TMS) alleviates depressive symptoms (57)(14 studies; *N* = 461 lesion cases and *N* = 251 stimulation cases), likewise available in MNI-152 standard space from NeuroVault at https://neurovault.org/images/787859. Both maps were parcellated according to the 68 cortical regions of the Desikan-Killiany atlas.

A map of loss-of-function pertaining to motor skill was obtained from the classic work of Karl Kleist, who assigned functions to different Brodmann areas based on the deficits observed following traumatic brain injuries and focal brain lesions (137). We used the version of the Kleist atlas recently digitised and shared in MNI standard space by Pijnenburg et al. (138). A binary map was obtained by assigning value of ‘1’ to regions whose lesion produced deficits of motor skill (e.g., ‘dexterity’, ‘individual movements’, ‘motor skill’, ‘individual consecutive action’, as per the Supplementary Materials of Pijnenburg et al. (138)), and ‘0’ otherwise. The resulting binary surface-based MNI-space map was then resampled to the Desikan-Killiany atlas, such that the value of each Desikan-Killiany region reflects the fraction of its vertices that were assigned ‘1’ in the corresponding location of the Kleist map.

### Unpublished meta-analysis of disorders of consciousness

The meta-analytic map of disorders of consciousness is the one used in (136). The map corresponds to a coordinate-based meta-analysis of sMRI, fMRI and [F^18^]-fluordoxyglucose (FDG)-PET studies of patients with disorders of consciousness, based on 39 studies, 40 experiments and 1,156 participants. We used maps of decreased cerebral integrity (as indexed by decreased sMRI or resting-state fMRI and [F^18^]FDG-PET metrics) in disorders of consciousness, as compared to healthy volunteers. The preprint was first available online in January 2026. Full details are provided in (136) and in the associated PROSPERO protocol (CRD42022327151). We used the unthresholded map in MNI-152 standard space for the analysis, parcellated into the 68 cortical regions of the Desikan-Killiany atlas.

### Regional morphometric disease association from ENIGMA consortium

Spatial maps of case-versus-control cortical thickness were obtained by including all the neurological, neurodevelopmental, and psychiatric diagnostic categories available from the ENIGMA (Enhancing Neuroimaging Genetics through Meta-Analysis) consortium (17, 18) and the Enigma toolbox (https://github.com/MICA-MNI/ENIGMA) (14) and recent related publications (https://github.com/netneurolab/hansen_crossdisorder_vulnerability) (167). The ENIGMA consortium is a data-sharing initiative that relies on standardized image acquisition and processing pipelines, such that cortical thickness maps are comparable (18). Altogether, over 17 000 patients were scanned, against almost 22 000 controls. The values for each map are zscored effect sizes (Cohen’s *d*) of cortical thickness in patient populations versus healthy controls. For the main analysis, we use the magnitude of these changes as an anatomical measure of regional involvement in each disorder. We also replicate results using the original maps of case-control effect size. Imaging and processing protocols can be found at http://enigma.ini.usc.edu/protocols/. Here we include maps for autism spectrum disorder (168), attention-deficit/hyperactivity disorder (169), bipolar disorder (170), depression (171), and schizophrenia (172). For every disorder, we constructed a vector of cortical thickness changes, where each element represents a diagnostic category’s magnitude of change in cortical thickness at a specific region. Note that unlike the manual meta-analyses, and unlike the automated meta-analyses of NeuroSynth and Neuro-Query, the ENIGMA consortium does not share maps in volumetric MNI-152 standard space. Rather, brain maps are provided in terms of the 68 regions of the Desikan-Killiany atlas - being one reason for the choice of this atlas for our main analyses.

### Molecular signatures of regional disease risk

#### Genome-wide association studies

The ENIGMA toolbox (https://github.com/MICA-MNI/ENIGMA) (14) provides lists of disease-associated genes, obtained from genome-wide association studies: ADHD (173), autistic spectrum disorder (174), bipolar disorder (175), major depressive disorder (114) and schizophrenia (176).

#### DisGeNET database

Zeighami et al. (112) obtained lists of genes pertaining to 40 common human brain diseases, identified through the DisGeNET database (www.disgenet.org) (177, 178), a platform aggregated from multiple sources including curated repositories, GWAS catalogs, animal models, and the scientific literature. After removing substance-related disorders, those with overlapping names (e.g., ‘post-partum depression’ vs ‘depression’) and those with very broad names (e.g., ‘learning disorders’), we retained 33 disorders, categorised into 7 groups: Neurocognitive disorders and Dementias (Alzheimer Disease; Amnesia; Dementia; Delirium; Frontotemporal Lobar Degeneration); Neurodegenerative and movement disorders (Amyotrophic Lateral Sclerosis; Ataxia; Dystonia; Huntington Disease; Hereditary Spastic Paraplegia; Parkinson Disease; Tremor); Neurodevelopmental disorders (Autistic Disorder; Neurofibromatosis 1; Williams Syndrome); Brain tumors (Astrocytoma; Glioblastoma; Glioma; Gliosarcoma; Oligodendroglioma); Neurological disorders (Temporal Lobe Epilepsy; Hyperalgesia; Migraine Disorders; Multiple Sclerosis; Neuralgia); Sleep and Wake disorders (Narcolepsy; Sleep Disorders); and Mood and Behaviour disorders (Schizophrenia; Bipolar Disorder; Depressive Disorder; Seasonal Affective Disorder; Panic Disorder; Obsessive Compulsive Disorder).

As reported in (112), the gene set sizes range widely from frontotemporal lobar degeneration (11 genes) to schizophrenia (733 genes). For both GWAS-derived and DisGeNET-derived lists of risk genes, one map of regional molecular risk per disease was obtained by averaging the normalised maps of the corresponding genes (see below for details of human gene expression processing and normalisation).

### Multimodal networks of inter-regional interactions

#### Haemodynamic connectivity from functional MRI

Haemodynamic connectivity, commonly simply referred to as “functional connectivity”, captures how similarly pairs of cortical regions exhibit fMRI BOLD activity at rest (95). The fMRI BOLD time-series picks up on magnetic differences between oxygenated and deoxygenated haemoglobin to measure the haemodynamic response: the oversupply of oxygen to active brain regions (179).

The dataset of functional and structural neuroimaging data used in this work came from the Human Connectome Project (HCP, http://www.humanconnectome.org/), Release Q3 (180). Per HCP protocol, all subjects gave written informed consent to the HCP consortium. These data contained fMRI and diffusion weighted imaging (DWI) acquisitions from 100 unrelated subjects of the HCP 900 data release (180). All HCP scanning protocols were approved by the local Institutional Review Board at Washington University in St. Louis. Detailed information about the acquisition and imaging is provided in the dedicated HCP publications. Briefly, data were acquired using the following parameters. Structural MRI: 3D MPRAGE T1-weighted, TR = 2, 400 ms, TE = 2.14 ms, TI = 1, 000 ms, flip angle = 8 deg, FOV = 224 × 224, voxel size = 0.7 mm isotropic. Two sessions of 15-min resting-state fMRI: gradient-echo EPI, TR = 720 ms, TE = 33.1 ms, flip angle = 52 deg, FOV = 208 × 180, voxel size = 2 mm isotropic. Here, we used functional data from only the first scanning session, in LR direction. HCP-minimally preprocessed data (181) were used for all acquisitions. The minimal preprocessing pipeline includes bias field correction, functional realignment, motion correction, and spatial normalisation to Montreal Neurological Institute (MNI-152) standard space with 2mm isotropic resampling resolution (181). We also removed the first 10 volumes, to allow magnetisation to reach steady state. Additional denoising steps were performed using the SPM12-based toolbox CONN (http://www.nitrc.org/projects/conn), version 17f (182) To reduce noise due to cardiac and motion artifacts, we applied the anatomical CompCor method of denoising the functional data. The anatomical CompCor method (also implemented within the CONN toolbox) involves regressing out of the functional data the following confounding effects: the first five principal components attributable to each individual’s white matter signal, and the first five components attributable to individual cerebrospinal fluid (CSF) signal; six subject-specific realignment parameters (three translations and three rotations) as well as their firstorder temporal derivatives (183). Linear detrending was also applied, and the subject-specific denoised BOLD signal timeseries were band-pass filtered to eliminate both low-frequency drift effects and high-frequency noise, thus retaining frequencies between 0.008 and 0.09 Hz. The parcellated time-series were used to construct functional connectivity matrices as a Pearson correlation coefficient between pairs of regional time-series. A group-average functional connectivity matrix was constructed as the mean functional connectivity across all individuals.

#### Metabolic connectivity from dynamic FDG-PET

Metabolic connectivity indexes how similarly two cortical regions metabolize glucose over time and therefore how similarly two cortical regions consume energy (55). Here we followed the same procedures as in (55). Briefly, volumetric 4D PET images of [F^18^]-fluordoxyglucose (FDG, a glucose analogue) tracer uptake over time were obtained from Jamadar et al. (100). Specifically, 26 healthy participants (77% female, 18–23 years old) were recruited from the general population and underwent a 95 minute simultaneous MR-PET scan in a Siemens (Erlangen) Biograph 3-Tesla molecular MR scanner. Participants were positioned supine in the scanner bore with their head in a 16-channel radiofrequency head coil and were instructed to lie as still as possible with eyes open and think of nothing in particular. FDG (average dose 233 MBq) was infused over the course of the scan at a rate of 36 mL/h using a BodyGuard 323 MR-compatible infusion pump (Caesarea Medical Electronics, Caesarea, Israel). Infusion onset was locked to the onset of the PET scan. This data has been validated and analyzed previously in (184, 185).

PET images were reconstructed and preprocessed according to (185). Specifically, the 5700-second PET time-series for each subject was binned into 356 3D sinogram frames each of 16-second intervals. The attenuation for all required data was corrected via the pseudo-CT method (186). Ordinary Poisson-Ordered Subset Expectation Maximization algorithm (3 iterations, 21 subsets) with point spread function correction was used to reconstruct 3D volumes from the sinogram frames. The reconstructed DICOM slices were converted to NIFTI format with size 344×344×127 (voxel size: 2.09×2.09×2.03 mm^3^) for each volume. A 5 mm FWHM Gaussian postfilter was applied to each 3D volume. All 3D volumes were temporally concatenated to form a 4D (344×344×127×356) NIFTI volume. A guided motion correction method using simultaneously acquired MRI was applied to correct the motion during the PET scan. 225 16-second volumes were retained commencing for further analyses.

Next, the 225 PET volumes were motion corrected (FSL MCFLIRT (187)) and the mean PET image was brain extracted and used to mask the 4D data. The fPET data were further processed using a spatiotemporal gradient filter to remove the accumulating effect of the radiotracer and other low-frequency components of the signal (188). Finally, each time point of the PET volumetric time-series were registered to MNI152 template space using Advanced Normalization Tools in Python (ANTSpy, https://github.com/ANTsX/ANTsPy), parcellated to 68 cortical regions according to the Desikan-Killiany atlas, and time-series at pairs of cortical regions were correlated (Pearson’s *r*) to construct a metabolic connectivity matrix for each subject. A group-averaged metabolic connectome was obtained by averaging connectivity across subjects.

#### Electrophysiological connectivity from magnetoencephalogrphy

Electrophysiological connectivity was measured using magnetoencephalography (MEG) recordings, which tracks the magnetic field produced by neural currents (55). Resting state MEG data was acquired for *n* = 33 unrelated healthy young adults (age range 22–35 years) from the Human Connectome Project (S900 release (180)). The data includes resting state scans of approximately 6 minutes long and noise recording for all participants. MEG anatomical data and 3T structural MRI of all participants were also obtained for MEG preprocessing.

The present MEG data was first processed and used by Hansen et al. (55), Shafiei et al. (96). Resting state MEG data was preprocessed using the opensource software, Brainstorm (https://neuroimage.usc.edu/brainstorm/ (189)), following the online tutorial for the HCP dataset (https://neuroimage.usc.edu/brainstorm/Tutorials/HCP-MEG). MEG recordings were registered to individual structural MRI images before applying the following preprocessing steps. First, notch filters were applied at 60, 120, 180, 240, and 300 Hz, followed by a high-pass filter at 0.3 Hz to remove slow-wave and DC-offset artifacts. Next, bad channels from artifacts (including heartbeats, eye blinks, saccades, muscle movements, and noisy segments) were removed using Signal-Space Projections (SSP).

Pre-processed sensor-level data was used to construct a source estimation on HCP’s fsLR4k cortex surface for each participant. Head models were computed using overlapping spheres and data and noise covariance matrices were estimated from resting state MEG and noise recordings. Linearly constrained minimum variance (LCMV) beamformers was used to obtain the source activity for each participant. Data covariance regularization was performed and the estimated source variance was normalized by the noise covariance matrix to reduce the effect of variable source depth. All eigenvalues smaller than the median eigenvalue of the data covariance matrix were replaced by the median. This helps avoid instability of data covariance inversion caused by the smallest eigenvalues and regularizes the data covariance matrix. Source orientations were constrained to be normal to the cortical surface at each of the 8 000 vertex locations on the cortical surface, then parcellated according to the Desikan-Killiany atlas (70).

After preprocessing and parcellating the data, amplitude envelope correlations were performed between time-series at each pair of brain regions, for six canonical frequency bands separately (delta (2–4 Hz), theta (5–7 Hz), alpha (8–12 Hz), beta (15–29 Hz), low gamma (30– 59 Hz), and high gamma (60–90 Hz)). Amplitude envelope correlation is applied instead of directly correlating the time-series because of the high sampling rate (2034.5 Hz) of the MEG recordings. An orthogonalization process was applied to correct for the spatial leakage effect by removing all shared zero-lag signals (190). The composite electrophysiological connectivity matrix is the first principal component of all six connectivity matrices (55).

#### Gene co-expression from microarray transcriptomics

Correlated gene expression represents the transcriptional similarity between pairs of cortical regions (55, 103). Regional microarry expression data were obtained from 6 post-mortem brains (1 female, ages 24.0–57.0, 42.50 ± 13.38) provided by the Allen Human Brain Atlas (AHBA, https://human.brain-map.org (102)). We followed the same preprocessing as recently described (61). Briefly, the Allen Human Brain Atlas (AHBA) is a publicly available transcriptional atlas containing gene expression data measured with DNA microarrays and sampled from hundreds of histologically validated neuroanatomical structures across six (five male and one female) normal postmortem human brains. We extracted and mapped gene expression data to the 68 cortical ROIs of the Desikan-Killiany atlas using the abagen toolbox (version 0.1.1; https://github.com/rmarkello/abagen (191)). Data was pooled between homologous cortical regions to ensure adequate coverage of both left (data from six donors) and right hemisphere (data from two donors). Distances between samples were evaluated on the cortical surface with a 2mm distance threshold. Only probes where expression measures were above a background threshold in more than 50% of samples were selected. A representative probe for a gene was selected based on highest intensity. 15,633 genes survived these preprocessing and quality assurance steps. Finally, the region × region correlated gene expression matrix was constructed by correlating (Pearson’s *r*) the normalised gene expression profile at every pair of brain regions.

#### Receptor co-expression from Positron Emission Tomography

Receptor similarity indexes the degree to which the receptor density profiles at two cortical regions are correlated (55, 88). Here we followed the same procedures as in (55). Briefly, receptor densities were estimated using PET tracer studies for a total of 18 receptors and transporters, across 9 neurotransmitter systems, recently made available by Hansen and colleagues at https://github.com/netneurolab/hansen_receptors (88). These include dopamine (D_1_ (192), D_2_ (193–196), DAT (197), noradrenaline (NAT (198–201), serotonin (5-HT_1A_ (202), 5-HT_1_B (202–207), 5-HT_2A_ (208), 5-HT_4_ (208), 5-HT_6_ (209, 210) 5-HTT (208)), acetylcholine (*α*_4_*β*_2_ (211, 212), M_1_ (213), VAChT (214, 215), glutamate (mGluR_5_ (216, 217) NMDA (218, 219), GABA (GABA_A_ (220))), histamine (H_3_ (221)), cannabinoid (CB_1_ (222–225)) and opioid (MOR (226)). Volumetric PET images were registered to the MNI-ICBM 152 nonlinear 2009 (version c, asymmetric) template, averaged across participants within each study, then parcellated and receptors/transporters with more than one mean image of the same tracer (5-HT_1_B, D_2_, VAChT) were combined using a weighted average (88). A regionby-region receptor similarity matrix was constructed by correlating (Pearson’s *r*) receptor profiles at every pair of cortical regions.

#### Laminar similarity from BigBrain histology

Laminar similarity is estimated from histological data and aims to uncover how similar pairs of cortical regions are in terms of cellular distributions across the cortical laminae. Specifically, we use data from the Big-Brain, a high-resolution (20 *µ*m) histological reconstruction of a post-mortem brain from a 65 year old male (105, 227). Cell-staining intensity profiles were sampled across 50 equivolumetric surfaces from the pial surface to the white mater surface to estimate laminar variation in neuronal density and soma size. Intensity profiles at various cortical depths can be used to approximately identify boundaries of cortical layers that separate supragranular (cortical layers I–III) granular (cortical layer IV), and infragranular (cortical layers V-VI) layers.

The data were obtained on *fsaverage* surface (164k vertices) from the `BigBrainWarp` toolbox (228) and were parcellated into 68 cortical regions according to the Desikan-Killiany atlas (70). The region × region laminar similarity matrix was calculated as the partial correlation (Pearson’s *r*) of cell intensities between pairs of cortical regions, after correcting for the mean intensity across cortical regions. Laminar similarity was first introduced in Paquola et al. (105) and has also been referred to as “microstructure profile covariance”.

#### Similarity of subjective response to intracranial electrical stimulation

We obtained a matrix of inter-regional similarity of subjective experiences elicited by direct intracranial electrical stimulation, using the data from (56) as described above. Based on the intrinsic connectivity network that it belongs to, each cortical region has a ‘fingerprint’ given by its probability of eliciting each of the 8 classes of subjective experiences from (56) upon electrical stimulation. Correlation of these probability fingerprints produces a matrix of similarity of subjective elicitation between each pair of cortical regions.

#### Structural connectivity

We used diffusion MRI (dMRI) data from the same 100 unrelated participants (54 females and 46 males, mean age = 29.1 ± 3.7 years) of the HCP 900 participants data release (229), as described above. The diffusion weighted imaging (DWI) acquisition protocol is covered in detail elsewhere (181). The diffusion MRI scan was conducted on a Siemens 3T Skyra scanner using a 2D spin-echo single-shot multiband EPI sequence with a multi-band factor of 3 and monopolar gradient pulse. The spatial resolution was 1.25 mm isotropic. TR=5500 ms, TE=89.50ms. The b-values were 1000, 2000, and 3000 s/mm^2^. The total number of diffusion sampling directions was 90, 90, and 90 for each of the shells in addition to 6 b0 images. We used the version of the data made available in DSI Studio-compatible format at http://brain.labsolver.org/diffusion-mri-templates/hcp-842-hcp-1021 (230).

We adopted previously reported procedures to reconstruct the human connectome from DWI data. The minimally-preprocessed DWI HCP data (181) were corrected for eddy current and susceptibility artifact. DWI data were then reconstructed using q-space diffeomorphic reconstruction (QSDR (231)), as implemented in DSI Studio (www.dsi-studio.labsolver.org). QSDR is a model-free method that calculates the orientational distribution of the density of diffusing water in a standard space, to conserve the diffusible spins and preserve the continuity of fiber geometry for fiber tracking. QSDR first reconstructs diffusion-weighted images in native space and computes the quantitative anisotropy (QA) in each voxel. These QA values are used to warp the brain to a template QA volume in Montreal Neurological Institute (MNI) space using a nonlinear registration algorithm implemented in the statistical parametric mapping (SPM) software. A diffusion sampling length ratio of 2.5 was used, and the output resolution was 1 mm. A modified FACT algorithm (232) was then used to perform deterministic fiber tracking on the reconstructed data, with the following parameters (233): angular cutoff of 55◦, step size of 1.0 mm, minimum length of 10 mm, maximum length of 400 mm, spin density function smoothing of 0.0, and a QA threshold determined by DWI signal in the cerebrospinal fluid. Each of the streamlines generated was automatically screened for its termination location. A white matter mask was created by applying DSI Studio’s default anisotropy threshold (0.6 Otsu’s threshold) to the spin distribution function’s anisotropy values. The mask was used to eliminate streamlines with premature termination in the white matter region. Deterministic fiber tracking was performed until 1, 000, 000 streamlines were reconstructed for each individual.

For each individual, their structural connectome was reconstructed by drawing an edge between each pair of regions *i* and *j* from the Desikan-Killiany cortical atlas (234) if there were white matter tracts connecting the corresponding brain regions end-to-end; edge weights were quantified as the number of streamlines connecting each pair of regions, normalised by ROI distance and size.

A group-consensus matrix *A* across participants was then obtained using the distance-dependent procedure of Betzel and colleagues, to mitigate concerns about inconsistencies in reconstruction of individual participants’ structural connectomes (235). This approach seeks to preserve both the edge density and the prevalence and length distribution of inter- and intra-hemispheric edge length distribution of individual participants’ connectomes, and it is designed to produce a representative connectome (235, 236). This procedure produces a binary consensus network indicating which edges to preserve. The weight of each non-zero edge is then computed as the mean of the corresponding non-zero edges across participants.

### Macaque multimodal data

#### Macaque functional MRI

The non-human primate MRI data was made available as part of the Primate neuroimaging Data-Exchange (PRIME-DE) monkey MRI data sharing initiative, a recently introduced open resource for non-human primate imaging (78).

The data preprocessing and denoising followed the same procedures as in a previous publication (84). We used fMRI data from rhesus macaques (*Macaca mulatta*) scanned at Newcastle University. This samples includes 14 exemplars (12 male, 2 female); Age distribution: 3.9-13.14 years; Weight distribution: 7.2-18 kg (full sample description available online: http://fcon_1000.projects.nitrc.org/indi/PRIME/files/newcastle.csv and http://fcon_1000.projects.nitrc.org/indi/PRIME/newcastle.html).

Ethics approval: All of the animal procedures performed were approved by the UK Home Office and comply with the Animal Scientific Procedures Act (1986) on the care and use of animals in research and with the European Directive on the protection of animals used in research (2010/63/EU). We support the Animal Research Reporting of In Vivo Experiments (ARRIVE) principles on reporting animal research. All persons involved in this project were Home Office certified and the work was strictly regulated by the U.K. Home Office. Local Animal Welfare Review Body (AWERB) approval was obtained. The 3Rs principles compliance and assessment was conducted by National Centre for 3Rs (NC3Rs). Animal in Sciences Committee (UK) approval was obtained as part of the Home Office Project License approval. Animal care and housing: All animals were housed and cared for in a group-housed colony, and animals performed behavioural training on various tasks for auditory and visual neuroscience. No training took place prior to MRI scanning. Macaque MRI acquisition

Animals were scanned in a vertical Bruker 4.7T primate dedicated scanner, with single channel or 4-8 channel parallel imaging coils used. No contrast agent was used. Optimization of the magnetic field prior to data acquisition was performed by means of 2nd order shim, Bruker and custom scanning sequence optimization. Animals were scanned upright, with MRI compatible headpost or non-invasive head immobilisation, and working on tasks or at rest (here, only resting-state scans were included). Eye tracking, video and audio monitoring were employed during scanning. Resting-state scanning was performed for 21.6 minutes, with a TR of 2600ms, 17ms TE, Effective Echo Spacing of 0.63ms, voxels size 1.22 × 1.22 × 1.24. Phase Encoding Direction: Encoded in columns. Structural scans comprised a T1 structural, MDEFT sequence with the following parameters: TE: 6ms; TR: 750 ms; Inversion delay: 700ms; Number of slices: 22; In-plane field of view: 12.8 × 9.6 cm^2^ on a grid of 256 × 192 voxels; Voxel resolution: 0.5 × 0.5 × 2mm; Number of segments: 8.

The macaque MRI data were preprocessed using the recently developed pipeline for non-human primate MRI analysis, *Pypreclin*, which addresses several specificities of monkey research. The pipeline is described in detail in the associated publication (237). Briefly, it includes the following steps: (i) Slice-timing correction.

(ii) Correction for the motion-induced, time-dependent B0 inhomogeneities. (iii) Reorientation from acquisition position to template; here, we used the recently developed National Institute of Mental Health Macaque Template (NMT): a high-resolution template of the average macaque brain generated from in vivo MRI of 31 rhesus macaques (Macaca mulatta). (iv) Realignment to the middle volume using FSL MCFLIRT function. (v) Normalisation and masking using Joe’s Image Program (JIP-align) routine (http://www.nmr.mgh.harvard.edu/~jbm/jip/, Joe Mandeville, Massachusetts General Hospital, Harvard University, MA, USA), which is specifically designed for preclinical studies: the normalization step aligns (affine) and warps (non-linear alignment using distortion field) the anatomical data into a generic template space. (vi) B1 field correction for low-frequency intensity non-uniformities present in the data. (vii) Coregistration of functional and anatomical images, using JIP-align to register the mean functional image (moving image) to the anatomical image (fixed image) by applying a rigid transformation. The anatomical brain mask was obtained by warping the template brain mask using the deformation field previously computed during the normalization step. Then, the functional images were aligned with the template space by composing the normalization and coregistration spatial transformations.

##### Denoising

The aCompCor denoising method implemented in the CONN toolbox (182) was used to denoise the macaque functional MRI data, to ensure consistency with the human data analysis pipeline. White matter and CSF masks were obtained from the corresponding probabilistic tissue maps of the high-resolution NMT template (eroded by 1 voxel); their first five principal components were regressed out of the functional data, as well as linear trends and 6 motion parameters (3 translations and 3 rotations) and their first derivatives. To make human and macaque data comparable, the macaque data were also bandpass filtered in the same 0.008-0.09 Hz range used for human data. Out of the 14 total animals present in the Newcastle sample, 10 had available awake restingstate fMRI data; of these 10, all except the first animal had two scanning sessions available. Thus, the total was 19 distinct sessions across 10 individual macaques. Data were parcellated into the 82-ROI cortical parcellation of Kötter and Wanke (71). The parcellated time-series were used to construct functional connectivity matrices as a Pearson correlation coefficient between pairs of regional time-series. A group-average functional connectivity matrix was constructed as the mean functional connectivity across all individuals.

#### Macaque structural connectivity

Anatomical connectivity for the macaque brain was obtained from the fully weighted, whole-cortex macaque connectome recently developed by Shen and colleagues (238). This connectome was generated by combining information from two different axonal tract-tracing studies from the CoCoMac database (http://cocomac.g-node.org/main/index.php) (239) with diffusion-based tractography obtained from nine adult macaques (*Macaca mulatta* and *Macaca fascicularis*) (238). The resulting connectome provides a matrix of weighted, directed anatomical connectivity between each of the cortical ROIs of the Regional Mapping atlas of Kötter and Wanke (238).

#### Macaque cortical gene expression from stereo-seq

We used cortex-wide macaque gene expression data recently made available by (60), who combined single-nucleus RNA sequencing (“snRNA-seq”) with high-resolution, large-field-of view spatial transcriptomics from spatiotemporal enhanced resolution omics-sequencing (“stereo-seq”) (240). Specifically, the authors made available (https://macaque.digital-brain.cn/spatial-omics) post-mortem gene expression data covering 143 regions of the left cortical hemisphere of one 6yo male cynomolgus macaque (*Macaca fascicularis*). We refer the reader to (60) for details.

Briefly, Chen and colleagues obtained 119 coronal sections at 500-*µ*m spacing, covering the entire cortex of the left hemisphere, which were used for stereo-seq transcriptomics (60). Adjacent 50-*µ*m thick sections were also acquired for regional microdissection and singlenucleus RNA sequencing (snRNA-seq) analysis, as well as 10-*µ*m sections adjacent to each stereo-seq section, which were used for the anatomical parcellation of brain regions via immunostaining (60). As reported in (60), for each coronal section, the cortical region and layer parcellation were manually delineated on stereo-seq data background, based on cytoarchitectual pattern (e.g. cell density, cell size) revealed by total mRNA expression, nucleic acid staining, and NeuN staing of adjacent sections. Gene expression data were made available at https://macaque.digital-brain.cn/spatial-omics for 143 cortical regions of the left hemisphere, including prefrontal, frontal, cingulate, somatosensory, insular, auditory, temporal, parietal, occipital and piriform areas. On the data-sharing portal, separate normalised gene expression data are made available for each region and for each of its cortical layers. To make the gene expression comparable across our datasets, we used the version of the macaque gene expression data that was mirrored between hemispheres and mapped onto the “regional mapping” macaque atlas of Kötter and Wanke (71), as made available by (61). To obtain inter-regional covariance of gene expression, each gene expression pattern was zscored and then genes were correlated across regions, thereby obtaining a regions-by-regions matrix.

#### Macaque receptor density from in vitro receptor autoradiography

*In vitro* autoradiography data for 14 neurotransmitter receptors were obtained from (59): *AMPA, kainate, NMDA, GABA*_*A*_, *GABA*_*B*_, *GABA*_*A/BZ*_, *M*_*1*_, *M*_*2*_, *M*_*3*_, *α*_*1*_, *α*_*2*_, *5HT*_*1A*_, *5HT*_*2A*_, *D*_*1*_. The authors applied quantitative in vitro receptor autoradiography to label 14 neurotransmitter receptors in three male *Macaca fascicularis* brains (7.3±0.6 years old; body weight 6±0.8 kg) obtained from Covance Preclinical Services, where they were housed and used as control animals for pharmaceutical studies performed in compliance with legal requirements. Animal experimental procedures and husbandry had the approval of the respective Institutional Animal Care and Use Committee and were carried out in accordance with the European Council Directive of 2010 (59). We refer the reader to (59) for details.

Briefly, the data of density of receptors per neuron were made available for 109 cortical areas of the macaque brain, which were identified based on their cytoarchitecture and receptor-architecture characteristics (59). To enable comparison across datasets, we used the version of the macaque receptor density data that was mapped onto the Regional Mapping macaque cortical parcellation of Kötter and Wanke (71) as made available by (61). The matrix of similarity of regional receptor expression was obtained by correlating the z-scored receptor density patterns.

### Mouse multimodal data

#### Mouse functional MRI

The mouse fMRI data used here have been published before (64). In vivo experiments were conducted in accordance with the Italian law (DL 26/214, EU 63/2010, Ministero della Sanita, Roma) and with the National Institute of Health recommendations for the care and use of laboratory animals (64). The animal research protocols for this study were reviewed and approved by the Italian Ministry of Health and the animal care committee of Istituto Italiano di Tecnologia (IIT). All surgeries were performed under anesthesia.

Adult (*<* 6 months old) male C57BL/6J mice were used throughout the study. Mice were group housed in a 12:12 hours light-dark cycle in individually ventilated cages with access to food and water ad libitum and with temperature maintained at 21 ± 1 degrees centigrade and humidity at 60 ± 10%. All the imaged mice were bred in the same vivarium and scanned with the same MRI scanner and imaging protocol employed for the awake scans (see below).

A first group of mice (n = 10, awake dataset) underwent head-post surgery, scanner habituation and fMRI image acquisitions as described below. See (64) for the full surgical, habituation, and scanner protocol. The scans so obtained constitute the awake rsfMRI mouse dataset we used throughout our study. To prevent motion, each mouse was secured using an implanted headpost in the custom-made MRI-compatible animal cradle and the body of the mouse was gently restrained (for details of the headpost implantation and habituation protocol, see the original publication (64)). All scans were acquired at the IIT laboratory in Rovereto (Italy) on a 7.0 Tesla MRI scanner (Bruker Biospin, Ettlingen) with a BGA-9 gradient set, a 72 mm birdcage transmit coil, and a four-channel (awake, halothane) or three-channel (medetomidine-isoflurane) solenoid receive coil. Awake scans were acquired using a single-shot echo planar imaging (EPI) sequence with the following parameters: TR/TE=1000/15 ms, flip angle=60 degrees, matrix=100 x 100, FOV=2.3 × 2.3 cm, 18 coronal slices (voxel-size 230 × 230 × 600 mm), slice thickness=600 mm and 1920 time points, for a total time of 32 minutes (64). Preprocessing of fMRI images was carried out as described in previous work (64). Briefly, the first 2 minutes of the time series were removed to account for thermal gradient equilibration. Functional MRI time-series were then time despiked (3dDespike, AFNI), motion corrected (MCFLIRT, FSL), skull stripped (FAST, FSL) and spatially registered (ANTs registration suite) to an inhouse mouse brain template with a spatial resolution of 0.23 × 0.23 × 0.6 mm^3^. Denoising involved the regression of 25 nuisance parameters. These were: average cerebral spinal fluid signal plus 24 motion parameters determined from the 3 translation and rotation parameters estimated during motion correction, their temporal derivatives and corresponding squared regressors. No global signal regression was employed. In-scanner head motion was quantified via calculations of framewise displacement (FD). To rule out a contribution of residual head-motion, we further introduced frame-wise fMRI scrubbing (FD *>* 0.075 mm). The resulting time series were band-pass filtered (0.01-0.1 Hz band) and then spatially smoothed with a Gaussian kernel of 0.5 mm full width at half maximum. Finally, the time-series were trimmed to ensure that the same number of timepoints were included for all animals, resulting in 1414 volumes per animal. Finally data were parcellated into 72 cortical symmetric regions from the Allen Mouse Brain Atlas (CCFv3) (72).

#### Mouse structural connectivity

For the mouse structural connectome, we used a parcellated version of the high-resolution mouse connectome of Coletta et al. (62). Below, we summarise how Coletta and colleagues obtained the high-resolution mouse structural connectome.

The present mouse structural connectome is based on “high-resolution models of the mouse brain connectome (100 *µ*^3^) previously released by Knox and colleagues (241). The Knox connectome is based on 428 viral microinjection experiments in C57BL/6J male mice obtained from the Allen Mouse Brain Connectivity Atlas (http://connectivity.brain-map.org/). The connectome data were derived from imaging enhanced green fluorescent protein (eGFP)–labeled axonal projections that were then registered to the Allen Mouse Brain Atlas and aggregated according to a voxel-wise interpolation model” (62).

Before constructing the SC matrix, Coletta et al. ensured symmetry along the right-left axis for all the major macrostructures of the mouse brain. To this purpose, they “flipped each macrostructure (isocortex, hippocampal formation, subcortical plate, pallidum, striatum, pons, medulla, midbrain, thalamus, hypothalamus, cerebellum, and olfactory bulb) along the sagittal midline (once for the right hemisphere and once for the left hemisphere) and took the intersection with the respective nonflipped macrostructure. This procedure resulted in the removal of a set of nonsymmetric voxel (total fraction, 8.6%), the vast majority of which reside in fringe white/gray matter or cerebrospinal fluid/gray matter interfaces. The removal of these nonsymmetric voxels did not substantially affect the network structure of the resampled connectome, as assessed with a spatial correlation analysis between the symmetrized and nonsymmetrized right ipsilateral (i.e., squared) connectome”. Coletta et al. then filtered out fiber tracts and ventricular spaces, and estimated SC using a resampled version of the recently published voxel scale model of the mouse structural connectome (62), to make the original matrix computationally tractable. Resampling of the Knox et al. connectome was carried out by aggregating neighboring voxels according to a Voronoi diagram based on Euclidean distance between neighboring voxels, preserving the intrinsic architectural foundation of the connectome while minimizing spatial blurring and boundary effects between ontogenically distinct neuroanatomical divisions of the mouse brain, or white/gray matter, and parenchymal/ventricular interfaces (see Coletta et al. for details of the Voronoi aggregation scheme (62)). By averaging the connectivity profile of neighboring voxels based on their relative spatial arrangement, this strategy has also the advantage of mitigating limitations related to the enforced smoothness of source space used by the original kernel interpolation used by (62). A wholebrain connectome was then built under the assumption of brain symmetry. The obtained Voronoi diagram made it possible to map the results back into the original 100 *µ*^3^ three-dimensional coordinate system of the Allen Institute mouse brain connectome [CCFv3] (72). For each pair of the 72 Allen Atlas cortical regions that we employed, their structural connectivity was obtained by averaging the connectivity of the respective constituent voxels.

#### Mouse brain gene expression from in situ hybridization

Mouse gene expression profiles were obtained using in situ hybridization data from the Allen Mouse Brain Atlas (63). Briefly, the Allen Mouse Brain Atlas consists of data acquired from a pipeline that includes semi-automated riboprobe generation, tissue preparation and sectioning, in-situ hybridization (ISH), imaging, and data postprocessing. These data were acquired both sagittally and (for a smaller set of genes) coronally, and were further processed, aligned by the Allen Institute to their Common Coordinate Framework version 3 (CCFv3) reference atlas (72) and summarized voxelwise through a measure termed gene expression energy (defined as the sum of expressing pixel intensity divided by the sum of all pixels in a division), resulting in 3D gene expression images at a 200 *µ*m isotropic resolution. This gene expression energy increases in regions of high expression, and is bounded by zero in regions of no expression. For our study, we used gene expression energy data from the coronal dataset (4345 gene expression images corresponding to 4082 unique genes), because of its wholebrain coverage and data quality. Voxelwise gene expression data were further summarized as normalized mean expression within regions of interest as defined by the CCFv3 reference atlas. For each ROI and each gene, voxelwise expression energy data were averaged over voxels containing valid expression signal. Finally, independently for each region, gene expression data were normalized by regressing out mean gene expression across the brain.

#### Mouse Neuropixels maps

We used the dataset of brain maps reconstructed using in vivo electrophysiological recordings of 621, 733 neurons recorded during a decision-making task with 699 Neuropixel probes in 139 mice, recently released by the International Brain Laboratory consortium (9). The data were annotated with task components related to sensation, choice, action and reward, as well as internal cognitive states. Specifically, we use brain maps that report for each cortical region, the success of decoding, in units of *R*^2^. We refer the reader to the original publication (9) and extensive online documentation (https://www.internationalbrainlab.com/data) for full details.

#### Mouse dopamine D1 receptor map

A map of the distribution of dopamine D1 receptor-positive neurons in the mouse forebrain was obtained from Bjerke et al. (89), who processed microscopic images of immunohistochemically stained serial sections from male and female Drd1a-EGFP transgenic mice at five stages of development (postnatal day 17 (P17), P25, P35, P49, and adult), known as the DOPAMAP collection and originally used in the study of (242). We used adult values of regional density registered to the Allen Mouse brain Common Coordinate Framework, freely available on the EBRAINS knowledge-sharing platform (https://doi.org/10.25493/KB7F-4VW).

### Anatomical hierarchy from in vivo T1w:T2w ratio

For each species, we used maps of intracortical myelination (from T1w:T2w ratio) as an in vivo marker of the anatomical cortical hierarchy (61, 65, 110, 243). For the human, we used the map available in the neuromaps toolbox (1). For the macaque, we used the T1w:T2w ratio map originally from (244) and resampled to the Regional Mapping macaque cortical parcellation of Kötter and Wanke (71) by (61). For the mouse, we used the T1w:T2w ratio data from (65).

### Statistical analyses

The use of non-parametric tests alleviated the need to assume normality of data distributions (which was not formally tested). All tests were two-sided, with an *α* value of 0.05. The effect sizes were estimated using Cohen’s measure of the standardized mean difference, *d*. To ensure robustness to possible outliers, correlations were quantified using Spearman’s rank-based non-parametric correlation coefficient.

#### Spatial autocorrelation-preserving surrogates for spatial correlation

To guard against inflation of false positives in spatial correlations due to the inherent spatial autocorrelation of brain data, significance of spatial correlations is assessed against a population of 5, 000 null maps with preserved spatial autocorrelation (245). Spatial autocorrelation-preserving surrogates are generated using the spin test (163) for human cortical maps, which preserves the distribution of values while breaking their relationship with underlying anatomy; Moran spectral randomisation is used for human cortical-subcortical maps and also for macaque and mouse maps. Moran spectral randomisation generates null maps based on on the inverse Euclidean distances between parcel centroids, as implemented in the BrainSpace toolbox (https://brainspace.readthedocs.io/en/latest/) (246). Moran spectral randomisation quantifies the spatial autocorrelation in the data in terms of Moran’s *I* coefficient, by computing spatial eigenvectors known as Moran eigenvector maps. The Moran eigenvectors are then used to generate null maps by imposing the spatial structure of the empirical data on randomised surrogate data (164, 246).

#### Multivariate gene-cognition association with Partial Least Squares

Partial Least Squares (PLS) analysis was used to relate regional gene expression to functional associations. PLS analysis is an unsupervised multivariate statistical technique that decomposes relationships between two datasets *X*_*n×g*_ and *Y*_*n×f*_ into orthogonal sets of latent variables with maximum covariance, which are linear combinations of the original data (247, 248). In the present case, *X*_*n×g*_ is regional gene expression across *n* regions (68 for the human; 82 for the macaque; and 72 for the mouse) and *g* genes (the same set of 81 for each species). *Y*_*n×f*_ is the matrix of regional functional associations, across 22 cognitive operations.

PLS finds components from the predictor variables (regional gene expression) that have maximum covariance with the response variables (regional functional associations). The PLS components (i.e., linear combinations of the weighted variables) are ranked by the covariance between predictor and response variables so that the first few PLS components provide a low-dimensional representation of the covariance between the higher-dimensional data matrices. Concretely, this is achieved by performing singular value decomposition (SVD) on the matrix *Y* ^*′*^*X*, such that:

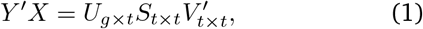

where *U*_*g×t*_ and *V*_*t×t*_ are orthonormal matrices consisting of left and right singular vectors, and *S*_*t×t*_ is a diagonal matrix of singular values. The *i*th columns of *U* and *V* constitute a latent variable, and the *i*th singular value in *S* represents the covariance between singular vectors. The *i*th singular value is proportional to the amount of covariance between gene expression and cognitive relevance captured by the *i*th latent variable, where the effect size can be estimated as the ratio of the squared singular value to the sum of all squared singular values.

The statistical significance of the covariance explained by each PLS model is tested by permuting the response variables 10 000 times while considering the spatial dependency of the data by using spatial autocorrelation-preserving permutation tests (164, 249). The p-value is computed as the proportion of null singular values that are greater in magnitude than the empirical singular values. Thus, these p-values represent the probability that the observed spatial correspondence between genes and cognitive operations could occur by randomly correlating maps with comparable spatial autocorrelation.

#### Network null models

We used two different network null models to disambiguate the role of connectome topology and geometric embedding in shaping control energy (249). The first null model is the well-known Maslov-Sneppen degree-preserving rewired network, whereby edges are swapped so as to randomise the topology while preserving the exact binary degree of each node (degree sequence), and the overall distribution of edge weights (250). As a second, more stringent null model, we adopted a null model that in addition to preserving exactly the same degree sequence and exactly the same edge weight distribution as the original network, also approximately preserves the original network’s edge length distribution (based on Euclidean distance between regions), and the weight-length relationship (251).

For each null model, we generated a population of 500 null networks starting from the empirical connectome, and computed the control energy between each pair of LLM-generated brain maps, as done for the empirical connectome. We compared the overall control energy between all possible states obtained from the empirical connectome and from the distribution of null instances.

For the analysis of modularity of disorder association networks, the modularity score is computed using the Newman-Girvan algorithm (252, 253) based on the assignment into 7 disease categories from the ICD manual. To assess the significance of the difference in modular structure between the gene expression and LLM-derived matrices, we build a null distribution by re-computing the difference in modularity according to the same assignments, after randomly rearranging the matrix (5,000 nulls).

#### Dominance analysis

To consider all inter-regional interactions together and evaluate their respective contributions to LLM-derived functional similarity, we performed a dominance analysis with all multimodal empirical networks as predictors, and the network of functional similarity from LLMs as target. Dominance analysis seeks to determine the relative contribution (“dominance” of each independent variable to the overall fit (adjusted *R*^2^)) of the multiple linear regression model (https://github.com/dominance-analysis/dominance-analysis) (106). This is done by fitting the same regression model on every combination of predictors (2^*p*^ −1 submodels for a model with *p* predictors). Total dominance is defined as the average of the relative increase in *R*^2^ when adding a single predictor of interest to a submodel, across all 2^*p*^ − 1 submodels. The sum of the dominance of all input variables is equal to the total adjusted *R*^2^ of the complete model, making the percentage of relative importance an intuitive method that partitions the total effect size across predictors. Therefore, unlike other methods of assessing predictor importance, such as methods based on regression coefficients or univariate correlations, dominance analysis accounts for predictor–predictor interactions and is interpretable. We use a null model with label permutation (5,000 iterations) to establish the probability that in each of the three species, the top-ranked predictor will be a functional network just by chance. Functional networks are defined as follows. For human: fMRI, MEG, FDG-PET, and electrical co-elicitation; for macaque: fMRI; for mouse: fMRI. At each iteration, the null model permutes predictors’ labels independently within each species; we then record whether the topranked predictor in each species is functional, and count a success when the top-ranked predictor is functional in all three species. A one-sided p-value is then computed as the proportion of permutations yielding a success.

## Supporting information

Supplementary Results and Figures

## Data and code availability

We have made freely and openly available the code for the NeuroNaut framework at https://github.com/andrealuppi/NeuroNaut. NeuroSynth metaanalytic maps are freely available from the NeuroSynth database at https://github.com/neurosynth/neurosynth. NeuroQuery meta-analytic maps are freely available from https://neuroquery.org/. ENIGMA cortical thickness data are provided as part of the ENIGMA Toolbox (v1.1.3), available at https://github.com/MICA-MNI/ENIGMA. Maps of subjective experience probabilities elicited by intracranial electrical stimulation are available from the supplementary materials of (56). Lesion network mapping circuits for depression and psychosis are available from NeuroVault at https://neurovault.org/images/787859 and https://neurovault.org/collections/20510. Mouse Neuropixels maps are available from the International Brain Laboratory at https://www.internationalbrainlab.com/data. The neuromaps toolbox for fetching human brain maps (version 0.0.1) is freely available at https://netneurolab.github.io/neuromaps/. Human Connectome Project Young Adult fMRI and diffusion MRI data are available from https://www.humanconnectome.org/study/hcp-young-adult. Diffusion MRI data for the Human Connectome Project in DSI Studio-compatible format are available at http://brain.labsolver.org/diffusion-mri-templates/hcp-842-hcp-1021. The Human PET receptor and transporter maps are available at https://github.com/netneurolab/hansen_receptors. The macaque fMRI data are available from the PRIME-DE database (http://fcon_1000.projects.nitrc.org/indi/indiPRIME.html). The macaque connectome is available on Zenodo at https://doi.org/10.5281/zenodo.1471588. The CoCo-Mac database on which it is based, is also available online at http://cocomac.g-node.org/main/index.php?. The mouse fMRI data and mouse structural connectome are available from author A.G. Human gene expression data from the Allen Human Brain Atlas are available at https://human.brain-map.org/. Macaque cortical gene expression data from (60) are available at https://macaque.digital-brain.cn/spatial-omics. The dataset is provided by Brain Science Data Center, Chinese Academy of Sciences (https://braindatacenter.cn/). Mouse gene expression data from in situ hybridization are available from the Allen Mouse Brain Atlas (63) at https://mouse.brain-map.org/. The original macaque receptor density data from autoradiography are available from https://balsa.wustl.edu/study/P2Nql and https://search.kg.ebrains.eu/instances/de62abc1-7252-4774-9965-5040f5e8fb6b (59). Regional density of D1R-expressing neurons in the adult mouse brain from (89) is freely available on the EBRAINS knowledge-sharing platform (https://doi.org/10.25493/KB7F-4VW). The original map of macaque intracortical myelination from T1w:T2w ratio from (59) is available at https://balsa.wustl.edu/study/P2Nql. Human networks of metabolic connectivity, electrophysiological connectivity, and laminar similarity are are available on GitHub at https://github.com/netneurolab/hansen_many_networks/tree/v1.0.0. The meta-analysis map of disorders of consciousness is available for scientific collaboration from author A.S., and will be made openly available upon publication of the original study (136). Digitised versions of the Kleist and Brodmann atlases in MNI-152 standard space are available from (138). The abagen toolbox for processing of the AHBA human transcriptomic dataset is available at https://abagen.readthedocs.io/. The BrainSpace toolbox for Moran spectral randomisation and spin-tests is available at https://brainspace.readthedocs.io/en/latest/. The netneurotools toolbox is freely available at https://github.com/netneurolab/netneurotools. The new NeuroSieve tool for refining NeuroSynth maps is made freely available at https://github.com/netneurolab/neurosieve. MATLAB code used to generate geometry-preserving null networks is freely available at https://www.brainnetworkslab.com/coderesources. Third-party Python software (version 3.8 was used) for dominance analysis is freely available at https://github.com/dominance-analysis/dominance-analysis. The CONN toolbox (v17f) for fMRI preprocessing is freely available at http://www.nitrc.org/projects/conn.

## Acknowledgments

The authors are grateful to Dr Michael D. Fox and Dr Shan Siddiqi for making available the lesion circuit maps of depression and psychosis; and to members of the Network Neuroscience Lab for valuable feedback. AIL acknowledges the support of St John’s College, Cambridge; and a Wellcome Early Career Award (grant number 226924/Z/23/Z). We also acknowledge support of the Brain Canada Foundation, through the Canada Brain Research Fund with the support of Health Canada [to D.B.]; the National Institutes of Health (grants no. NIH R01 AG068563A and NIH R01 R01DA053301-01A1) [to D.B.]; the Canadian Institute of Health Research (grants no. CIHR 438531 and CIHR 470425) [to D.B.]; the Healthy Brains Healthy Lives initiative (Canada First Research Excellence fund) [to D.B.]; Google (Research Award, Teaching Award) [to D.B.]; the CIFAR Artificial Intelligence Chairs programme (Canada Institute for Advanced Research) [to D.B.]; the European Research Council (ERC) under the European Union’s Horizon 2020 research and innovation program [DISCONN; no. 802371 to A.G.; and no. 101125054 - BRAINAMICS to A.G.]; BM acknowledges support from the Natural Sciences and Engineering Research Council of Canada (RGPIN-2017-04265), Canadian Institutes of Health Research (PJT-180439), and Canada Research Chairs Program (CRC-2022-00169). ZQL and FM acknowledge support from the Fonds de Recherche du Québec – Nature et Technologies (FRQNT). Human connectome data were provided by the Human Connectome Project, WU–Minn Consortium (1U54MH091657; Principal Investigators David Van Essen and Kamil Ugurbil) funded by the 16 National Institutes of Health (NIH) institutes and centers that support the NIH Blueprint for Neuroscience Research, and by the McDonnell Center for Systems Neuroscience at Washington University. For the purpose of open access, the authors have applied a Creative Commons Attribution (CC BY) licence to any Author Accepted Manuscript version arising from this submission. Any opinions, findings, and conclusions or recommendations expressed in this material are those of the authors and do not reflect the views of the funders.

## Supplementary Materials

