## Supplementary Results and Figures for "Cognitive cartography of mammalian brains using meta-analysis of AI experts"

*(i) Formal probability estimation from the Bradley-Terry model*

Although we ask AI experts to return estimated probabilities, the resulting region-function association scores are not probabilities in the formal sense. Rather, they are similar in nature to the subjective probabilities that would be elicited from human experts by a similar query. Just like human experts, AI experts may be trained on different bodies of evidence and use the same scale differently. Indeed, some AI experts tend to provide generally higher or lower values than others (Fig. S6, Fig. S31 and Fig. S32), yet the rankings that they provide for the same function are highly consistent with each other (Fig. S2) – just like different human neuroscientists may be more or less liberal in the scores that they attribute, yet broadly agree about the ranking of regions. This is the reason for our approach of z-scoring each LLM map prior to averaging them, thereby highlighting only the relative differences. Note that for the present purposes of generating brain maps, what matters is indeed the relative values attributed to different regions for the same cognitive function by the same AI expert, rather than the absolute value of each region. This is because we never consider brain regions in isolation, but always in the context of all other regions comprising the relevant brain map. This therefore only requires each LLM to be *internally* calibrated, by which we mean that for the same function, more important regions get assigned higher values. Correlations of individual LLMs' cognitive maps with NeuroSynth (Fig. S4) and with each other (Fig. S2) confirm that this is the case.

In other words, the approach used in the main analysis effectively uses the region-function association scores as an indicator of the relative rank that each AI expert assigns to each region. In the literature on learning from human preferences, discrepant numeric scores for the same item may be observed due to differences between raters' internal standards, interpretation of the numeric scale, or calibration against prior examples. A way to overcome such potential discrepancies is to use comparative judgments instead of absolute rating scores, and then aggregate them into formal probabilities (76, 162). We therefore adopt this pairwise-judgment approach as an alternative way of eliciting brain maps from AI experts, with the resulting scores being formally interpretable as probabilities. This approach involves two steps. First, rather than asking AI expert about individual regions and using the resulting scores to rank them, we explicitly ask each AI expert to choose between a *pair* of regions *i* and *j* in a head-to-head comparison. No explicit scores are solicited: just a binary choice between the two options (Fig. S33a). This process is repeated systematically for every pair of regions. Second, the resulting regional patterns of wins and losses in this 'tournament' are used as inputs to the Bradley-Terry model, which uses them to associate each item (here, a region) with a latent parameter (Fig. S33a), such that differences between latent parameters provide the formal probabilities that make the observed pairwise comparison data as likely as possible (76). In other words, rather than explicitly asking for numeric scores and using them to rank regions, which implicitly requires making global judgments, the Bradley-Terry model starts from pair-wise rankings (which only requires local judgments) and then infers the numerical scores that best explain them.

Two findings emerge. First, the brain maps of latent probabilities from the BT model are much more focalised, placing most of the emphasis on a small number of regions that overwhelmingly dominate the respective comparisons (Fig. S34). Second, the resulting brain maps are highly and significantly correlated both with the NeuroSynth maps pertaining to the same functions, and even more so with the corresponding brain maps of region-function association scores obtained from the same AI experts (with Spearman's *r* ranging between 0.79 and 0.94; Fig. S33b). In other words, we find that the approach of eliciting direct region-function association scores is internally calibrated, and highly consistent with the latent probabilities obtained from pairwise comparisons (i.e., they provide very similar rankings). Crucially, eliciting region-function association scores has the advantage of only requiring one LLM query for each of the *n* regions in one's chosen atlas, whereas the pairwise approach requires $n \times (n-1)$ queries and is therefore substantially more expensive. Hence while both approaches are made available in our computational framework, we favour the approach of direct region-function association scores, which provides highly similar results for a fraction of the cost.

*(ii) Justifications for LLMs' region-function association scores*

We also provide measures to increase transparency and provide insight into the automated experts' rationale for their region-function association scores. In addition to asking for a numerical region-function association score between 0 and 1 intended to approximate the probability of regional association with a function, we implement a `-justify` command whereby the automated expert will also produce a 1-2 sentence textual explanation for its rating. Examples are provided as Supplementary Materials.

As example, we show the justifications provided by `Claude-Opus-4.6`, `GPT-5.4`, and `Gemini 3.1 Pro` for 'memory' across regions that we expect to be either highly relevant (parahippocampal) or to only exhibit limited involvement (pericalcarine/V1).

Parahippocampal:

- `Claude-opus-4.6`: "The left parahippocampal cortex is extensively implicated in memory, particularly episodic and verbal memory encoding/retrieval, with robust evidence from fMRI, lesion, and intracranial studies."

- GPT-5.4: "Left parahippocampal cortex is very frequently linked to episodic, contextual, and spatial memory in human imaging and lesion studies."
- Gemini-3.1-pro: "The left parahippocampal gyrus is a core part of the medial temporal lobe memory system, almost universally cited in human literature for its critical role in episodic and contextual memory."

Pericalcarine:

- Claude-opus-4.6: "Pericalcarine cortex is primary visual cortex (V1), primarily linked to visual processing. Memory associations exist mainly via visual imagery/memory encoding but are relatively infrequent."
- GPT-5.4: "Pericalcarine cortex is primarily visual; memory links are uncommon and usually indirect via visual imagery or recall, not core memory processes."
- Gemini-3.1-pro: "The pericalcarine cortex (V1) primarily processes low-level vision. While occasionally noted in visual working memory studies, it is rarely the primary focus of human memory literature."

We also see that justifications provided for the involvement of the precentral gyrus (primary motor cortex) in 'manipulation' are closely aligned with our own neuroscientific expertise – both in terms of interpreting manipulation as referring to object handling, and in terms of the importance of motor regions for this function (we elaborate further on this point in Supplementary Results (iii):

- Claude-opus-4.6: "The left precentral gyrus (primary motor cortex) is strongly implicated in manipulation, particularly right-hand object manipulation and skilled hand movements, frequently reported in fMRI studies."
- GPT-5.4: "Left precentral gyrus contains primary motor hand areas, frequently activated in human studies of grasping and object manipulation, though parietal/premotor regions also contribute."
- Gemini-3.1-pro: "The left precentral gyrus houses the primary motor cortex, which is heavily cited in human neuroimaging and lesion studies for executing voluntary motor actions like object manipulation."

Consistent and sensible justifications are also provided when AI experts are asked about the non-scientific notion of 'telepathy', used here as a negative control:

- Claude-opus-4.6: "Telepathy is not a scientifically validated phenomenon, and the left precentral gyrus (primary motor cortex) is associated with motor control, not purported extrasensory functions."
- GPT-5.4: "Telepathy has no established peer-reviewed neural basis; the left precentral gyrus is primarily linked to voluntary motor control."
- Gemini-3.1-pro: "Telepathy is not a scientifically recognized phenomenon; peer-reviewed neuroscience literature associates the left precentral gyrus with motor control, not telepathic abilities."

These examples bring to light several noteworthy observations. First, on many occasions the AI experts' answers explicitly identify one or more alternative names for the same region (e.g., pericalcarine/primary visual cortex/V1, precentral/primary motor). Second, they also correctly identify (without being explicitly prompted) that the left precentral gyrus controls the movements of the *right* hand. Third, the explanations provided by different AI experts are highly consistent with each other. By asking LLMs to rate the strength of support expressed in each other's textual justifications, we also find strong and significant correlations with the region-function association scores originally provided alongside the justifications (Fig. S35).

To formally test the agreement between different LLMs' justifications, we embed the justifications through the OpenAI API, and quantify the alignment (correlation) of the resulting high-dimensional embedding vectors (3072 dimensions) (Fig. S36a). In other words: how aligned are the explanations in latent space? We show that the justifications provided across regions and functions by Anthropic's Claude-4.6-Opus, OpenAI's GPT-5.4 and Google's Gemini 3.1 Pro, as well as the open-weight models OSS-120b, Gemma-4-31b-instruct, and Qwen3-235b-a22b-2507, are all significantly and positively correlated in embedding space (Fig. S36b). As a further robustness check, we confirm that justification embeddings are also significantly more similar when they pertain to the same function than different functions (Fig. S36c).

*(iii) Refining NeuroSynth inputs improves match with LLM-derived map*

Neurosynth associates terms with activation coordinates via automated text-mining of study abstracts (12). Its standard implementation relies on traditional frequency-based natural language processing methods that ignore the semantic role of each term: a well-known limitation of the approach. For example, the term 'manipulation' may appear in an abstract as a cognitive operation of interest (meaning 'object handling', whether physically or in the metaphorical sense of 'symbol transformation') but it may also feature in the description of the study in the extraneous sense of 'experimental manipulation' (e.g., "[...] impact of neuropharmacological manipulation on brain activity dynamics" (254)). NeuroSynth does not distinguish between these meanings, and coordinates would be included from abstracts featuring both uses of the word. This issue can degrade the quality of the resulting meta-analytic maps, by including coordinates from irrelevant studies.

However, this limitation of NeuroSynth also offers an opportunity. If refining the NeuroSynth map by filtering out irrelevant abstracts brings the resulting brain map into closer alignment with the LLM-generated map, then we have evidence that AI experts are performing their own summary of the literature, and not just returning previously published NeuroSynth maps, because in that case the correspondence should actively decline after filtering instead. Additionally, we could further increase our credence that the AI experts' summaries of the literature are appropriate, being a better match for a more accurate than a less accurate alternative map. This is the hypothesis that we pursue next.

To refine the NeuroSynth maps, we pass the original abstracts of each study to a large language model not used elsewhere in the study (`Mistral Small 4`) and ask it to judge whether the term of interest was the study's primary condition, a contrast condition, used only in the background, or entirely absent, alongside a confidence score (see Methods). Studies are retained if the model judges with high confidence that the term of interest is the primary condition. This procedure can be applied to a broader set of articles than the ones originally selected for inclusion by NeuroSynth, thereby simultaneously reducing false positives while also reducing false negatives. The NeuroSynth model is re-run on the coordinates extracted from the subset of studies that meet this criterion.

We find that filtering the NeuroSynth abstracts for 'manipulation' increases the prominence of somatomotor regions (Fig. S37a), and also drastically increases the spatial correspondence with the map generated by AI experts, from Spearman's $r = 0.38$ to $r = 0.62$ ($p < 0.001$; Fig. S37b). In other words, improving the quality of the NeuroSynth map, also brings it into closer alignment with the LLM-generated map. As an additional validation, we compare all three maps (NeuroSynth, NeuroSynth-refined, and LLM-generated) against a brain map of lesion-induced loss-of-function pertaining to motor skill, recently digitised by Pijnenburg et al. (138) from the classic work of Karl Kleist, who assigned functions to different Brodmann areas based on the deficits observed following traumatic brain injuries and focal brain lesions (137). Neither the original nor the refined NeuroSynth maps exhibit significant spatial correspondence with the Kleist loss-of-function map, although filtering does substantially improve the correlation (from Spearman's $r = 0.16$ to $r = 0.40$; Fig. S38) – as we would expect it the original map were contaminated by irrelevant studies, which are removed by filtering. In contrast, the map of 'manipulation' generated by AI experts exhibits strong and statistically significant correlation with the map of lesion-induced loss of motor skill (Spearman's $r = 0.64, p = 0.002$). Hence, multiple sources of evidence point to the brain map generated by AI experts as a more faithful representation of the neural circuits involved in 'manipulation'.

*(iv) Automated neuroscience experts map a broader repertoire of 123 cognitive operations*

Is LLMs' neuroscientific expertise restricted to broad domains of cognition, or can they also offer more fine-grained insights? To address this question we consider a much broader set of 123 cognitive and behavioural terms from the Cognitive Atlas (148), selected in an unbiased manner based on extensive use in previous publications (130, 131, 163, 164). Terms range from umbrella terms ("memory", "emotion") to specific cognitive processes ("visual attention", "episodic memory"), behaviours ("eating", "sleep"), and emotional states ("fear", "anxiety") (Table 2).

For each of the 123 terms, we compare the brain maps generated by NeuroSynth and by AI experts. Controlling for spatial autocorrelation, we find that 101 terms exhibit significant positive correlations; 1 exhibits a negative significant correlation; and 21 are non-significant (Fig. S39a). When considering pairs of maps that do not agree between AI experts and NeuroSynth, several observations emerge. First, the non-significant terms include 'consciousness', for which independent validation indicates that the map generated by AI experts is actually the more appropriate of the two (Fig.S25. Second, the only significant negative correlation pertains to 'morphology'. The brain map generated by automated neuroscience experts displays prominent peaks in language-related regions - whereas the NeuroSynth map peaks in medial prefrontal regions, with some of the lowest scores in language-related regions (Fig. S39b). Since in cognitive science 'morphology' refers to the internal structure of words, the language-related map generated by LLMs appears to be a superior candidate. Third, many of the non-significant terms such as 'association', 'insight', 'goal', 'context', are very broad and likely to feature in the text of neuroscience papers, not as topics of study, but as part of the scientific discourse (e.g., "our study's *focus* is to obtain *insight* into the *association* between X and Y in the *context* of Z") – which might then be misinterpreted by NeuroSynth analogously to the case of 'manipulation' (Fig. S37). Indeed, the 21 non-significant terms appear to be generally challenging to summarise: 8 of them are also non-significant between NeuroSynth and NeuroQuery ('focus', 'induction', 'insight', 'sustained attention', 'effort', 'strength', 'context', 'visual perception'), with a 9th term ('balance') being significantly *negatively* correlated between the two engines for term-based meta-analysis (Fig. S39c). To evaluate whether semantically ambiguous use of these terms may have resulted in noisy NeuroSynth maps and hindered convergence with AI experts and NeuroQuery, we turn to the same strategy adopted for 'manipulation': re-computing NeuroSynth maps after refining the abstract selection. The improvement is drastic. 18 of the 21 terms that were non-significant with the default NeuroSynth maps, become significantly correlated with the maps generated by AI experts upon refinement (Fig. S40). Refinement also significantly increases the magnitude of correlations between NeuroSynth maps, and LLM-generated maps, more than doubling it from a mean Spearman's *r* of $0.19$ ($SD = 0.12$) without refinement, to a mean of $r = 0.48$ ($SD = 0.22$) upon refinement ($t(20) = -5.66, p < 0.001$, Hedge's $g = -1.57$ from two-sided paired-samples t-test; Fig. S41).

Figure S42 provides illustrative examples. The brain maps for 'visual perception' appear similar across the four approaches (Fig. S42a), and broadly match our neuroscientific expectation. The main exception is medial PFC for the original NeuroSynth map, which likely explains its lack of correlation with LLMs and NeuroQuery, and which is removed upon refinement, whereupon the correlation with the LLM map becomes significant. In contrast, considering 'balance' reveals three entirely different brain maps for LLMs, NeuroSynth and NeuroQuery (Fig. S42b). Yet upon refinement, the NeuroSynth map becomes much more similar (significantly so, in fact; Fig. S40) to the map generated by AI experts, with high importance for somatomotor regions, which we would expect to see involved in maintaining one's balance (Fig. S42b). AI experts, NeuroSynth, and NeuroQuery also produce very different maps for 'sustained attention' and 'focus' (Fig. S42c-d). Crucially, focus and sustained attention refer to closely related cognitive operations, yet the brain maps for NeuroQuery and NeuroSynth are entirely different, both for the same meta-analytic engine and for the same term (Fig. S42c-e). In stark contrast, the brain maps for 'sustained attention' and 'focus' generated by AI experts are highly and significantly correlated (Spearman's $r = 0.93, p < 0.001$; Fig. S42e). The situation is very different for the refined NeuroSynth maps, which (i) are now visibly more similar (and significantly correlated) with the corresponding maps generated by AI experts (Fig. S42c-d); and (ii) are now also significantly correlated with each other (Spearman's $r = 0.78, p < 0.001$; Fig. S42e), thereby matching both our neuroscientific expectations, and the behaviour of AI experts' maps.

In sum, we find that AI experts produce brain maps that converge with the corresponding NeuroSynth meta-analysis for over 100 out of 123 terms. We also provide convergent evidence that (i) nearly half (9/21) of the non-significant terms are also inconsistent between NeuroSynth and NeuroQuery, suggesting that they may be challenging terms for automated meta-analyses in general; (ii) among the discordant terms, we find several instances where the discrepancy favours the maps generated by AI experts, either based on human expertise (e.g., 'morphology'), or greater consistency for synonymous terms (e.g., 'focus' and 'sustained attention'), or on the ground of superior match to a third-party reference (e.g., 'consciousness'); and (iii) crucially, refining the set of abstracts used to compute the NeuroSynth maps, thereby excluding studies where polysemous words are not used in the relevant sense (such as 'experimental manipulation' or 'the focus of our study'), results in significance for 18 of the 21 terms. In other words, improving the quality of NeuroSynth search brings the resulting brain maps to converge with LLM-generated maps. Altogether, we find that AI experts' neuroscientific expertise is not limited to broad categories of human cognition, but rather it can also encompass a much more fine-grained ontology.

### *(v) LLM-generated brain maps reflect spontaneous human brain activity from rs-fMRI better than NeuroSynth and NeuroQuery maps*

Even at rest, human brain activity is neither static nor random, instead fluctuating to support ongoing cognition on a moment-to-moment basis. Thus, each volume of BOLD signal obtained during resting-state functional MRI can be viewed as a brain map. In turn, at each point in time we can ask how well this brain map matches, as a whole, the meta-analytic brain maps representing different cognitive operations: whether produced by NeuroSynth, or NeuroQuery, or AI experts (Fig. S43a). Remarkably, we find that the mean correlation between meta-analytic maps and instantaneous patterns of human rs-fMRI is significantly higher when using LLM-generated maps, than when using NeuroSynth (Fig. S43b) or NeuroQuery maps (Fig. S44a) – not only for the 8 canonical cognitive operations from (69), but also when considering a broader repertoire of up to 123 cognitive operations (131) (Fig. S44b). These results are especially noteworthy because NeuroSynth and NeuroQuery start from a position of advantage, being both derived from functional MRI. This analysis therefore suggests that although the brain maps of cognition generated by AI experts display high spatial correspondence with the meta-analytic maps from NeuroSynth and NeuroQuery (Fig. 1), they are not merely a degraded copy: on the contrary, LLM-generated maps transcend the maps generated by current meta-analytic engines in providing a more accurate depiction of human cognitive operations, corroborating the causal evidence from lesions (Fig. 6) and intracranial stimulation (Fig. 4).

*(vi) Systematic comparison between LLM-generated and NeuroSynth-generated brain maps as an engine for hypothesis generation*

Although we find strong and significant correlations between LLM-generated and NeuroSynth-generated cognitive maps, the differences between them can also be informative. Specifically, we ask which regions exhibit a NeuroSynth-LLM difference more extreme than what would be expected by chance, quantified using a null distribution of 'spun' maps with preserved spatial autocorrelation followed by FDR correction across regions (Fig. S45). As expected based on the previous analysis, we find that NeuroSynth appears to underestimate the importance of primary motor cortex for 'manipulation' (Fig. S45). Notably, however, NeuroSynth also appears to consistently attribute greater importance to primary visual cortex than LLMs do: not only for manipulation, but also for cognitive control, emotion, memory and reward - as well as attributing greater importance than LLMs to auditory cortex for 'vision' (Fig. S45).

The consistent attribution of a higher rank to unimodal (especially visual) regions by NeuroSynth versus the automated neuroscience experts could be an indication that such regions truly play a more widespread role for cognition than previously recognised - in line with recent reports emphasising the distributed nature of brain patterns supporting sensation and cognition (255). An alternative explanation could be due to the widespread use of visual stimuli and motor responses for most neuroimaging tasks, with the resulting activation foci being picked up by the NeuroSynth algorithm, whereas LLMs are able to identify them as irrelevant for the task at hand. Under this interpretation, AI experts provide a more accurate summary of the literature - as supported by the superior spatial correspondence of LLM-generated maps with maps obtained from alternative modalities such as lesions and intracranial stimulation (Fig. 4, Fig. 6, and Fig. S38). The emergence of these two contraposed accounts of unimodal regions' role in cognitive functions beyond their traditional ones, which could be resolved through targeted experimentation, represents an example of using automated neuroscience experts to generate testable hypotheses from the neuroscientific literature.

*(vii) Additional generalisation of human-specific findings to macaque and mouse brains*

Whereas Hansen et al. (131) focused on the relationship between cognition and molecular architecture, another study (130) investigated how transitions between different brain states (represented by the NeuroSynth maps of various cognitive operations, and replicated using maps of parameter estimates from in-scanner tasks) depend on the network architecture of the human structural connectome. Using principles of network control theory, they showed that the human connectome minimises the average energy required to transition between pairs of brain states, being more energetically efficient than any topologically random network, even when the number of connections (node degree) of each region is the same. Use of a more stringent null model that preserves not only the number of connections but also their length (thereby preserving the overall geometric embedding of the brain) led to an improvement over the degree-preserving networks, but still without reaching the energetic efficiency of the empirical human connectome – thereby demonstrating that network topology and geometry both play a role in the energetic efficiency of the human brain's wiring diagram.

As with the analysis replicating the study of Hansen et al. (131), here we proceed in two steps. First, we replicate the human results after replacing the NeuroSynth maps with brain maps from automated expert consensus (Fig. S24a,b). Transitions between the same 17 representative cognitive operations described above become significantly energetically more difficult if the connectome topology is disrupted, and even more difficult if both topology and geometry are randomised. Hence, NeuroSynth and automated expert consensus once again provides the same insights about human brain function (Fig. S24c).

Second, we repeat the same analysis in macaque and mouse, using LLM-derived cognitive maps and species-specific connectomes reconstructed from ex vivo tract-tracing (rather than in vivo diffusion MRI tractography). We recover the exact same pattern of results in both species: the empirical connectome is the most energetically efficient for supporting transitions between cognitive brain maps (Fig. S24d,e). Disrupting network topology incurs an energetic premium, and an even greater energetic cost is observed when both topology and geometry are randomised.

*(viii) Assessing LLMs' expertise of the anatomical nomenclature*

An advantage of using LLMs as automated neuroscience experts is that their knowledge of the literature is not confined to standard-space coordinates: just like human experts, but unlike NeuroSynth, BrainMap, or NeuroQuery. Here we provide formal confirmation that LLMs can correctly identify and match different names for the same cortical territories that are used in the literature.

We use an alternative prompting strategy whereby the LLM is asked to

> 'estimate the probability that the synonym/term **[**target**]** refers to the exact same anatomical location as the atlas region **[**region**]** [**hemisphere part**] of **[**species**]** brain'.

For example, the target may be 'V1' and this prompt is repeated for every region in the Desikan-Killiany atlas. We repeat this for multiple target regions, each with several synonyms encompassing anatomical, cytoarchitectonic, and functional labels (Table S1). For each target concept, we evaluated how well the LLM-derived regional relevance vector matched the corresponding anatomical ground truth. The ground-truth vector was defined over Desikan–Killiany atlas parcels, with entries set to 1 for parcels expected to correspond to the target region and 0 for all other parcels. For example, when the target concept was V1, the *pericalcarine* parcel was labelled as the positive ground-truth region, while all other parcels were labelled as negatives. We then computed the area under the receiver operating characteristic curve (ROC-AUC) between the LLM-provided scores and this binary ground-truth vector. This measures whether the LLM assigns higher scores to anatomically appropriate parcels than to unrelated parcels. In other words, we ask whether automated experts rank the correct anatomical parcel(s) above incorrect ones.

We find that across regions and LLMs, the mean AUROC is 0.96, with very rare exceptions for specific combinations of LLMs and region names, and going beyond what can be attributed to spatial autocorrelation alone (Fig. S46). In other words, automated experts possess sufficient neuroscientific knowledge to correctly identify different names for the same cortical regions across the literature.

As an alternative approach, we obtain justifications for LLMs' region-function association scores, for the anatomical Desikan-Killiany atlas and the cytoarchitectonic atlas of Brodmann (138, 161). Unlike the Desikan-Killiany atlas, the Brodmann atlas is defined in terms of cytoarchitecture rather than sulcal-gyral anatomy (161). Its region names are highly similar to each other (e.g., 'Brodmann Area 7', 'Brodmann area 17', 'Brodmann area 18') providing no indication about location or function without neuroscientific expertise. After embedding the justifications using the OpenAI API, we have a quantitative way to ask whether the embedding vectors provided for each function are more similar (higher correlation, or cosine similarity) for matched regions of the two atlases (e.g., 'Brodmann area 17' and 'pericalcarine'), as opposed to mis-matched regions (Fig. S47a). We find that this is indeed the case, across all functions (Fig. S47b). Thus, we see that not only can LLMs *explicitly* recognise multiple different names for the same region, in their capacity as neuroscience experts; they also *implicitly* reason about the same region in similar ways, even when described according to different atlas conventions.

*(ix) Robustness to atlas definition, scale, and inclusion of subcortex*

Although we have seen that AI experts can recognise different names for the same region in the literature without needing explicit standard-space coordinates, requiring pre-defined parcellations also comes with its own challenges compared to voxel-wise predictions. Reliance on discrete region names imposes limits on the spatial resolution that can be achieved, whereas many current meta-analytic engines typically provide voxel- or vertex-wise maps that can then be parcellated if desired. Note however that this is not universal across existing meta-analytic approaches: the ENIGMA consortium relies on the same Desikan-Killiany atlas used here (14) to release its meta-analytic maps of cortical morphometry for different disorders. Indeed, this was a major motivation behind our own choice of the Desikan-Killiany atlas for the present work.

The question may arise: are the present results critically dependent on the choice of atlas? We adopt several strategies to further mitigate this concern. First, we replicate results after including subcortical structures: thalamus, caudate, putamen, globus pallidus, accumbens, hippocampus, and amygdala (both left and right), corresponding to the Cammoun anatomical atlas (159) (Fig. S48). Notably, the LLM-derived clustering remains more faithful to ICD-11 classification than the clustering based on molecular fingerprints of risk-gene expression, even when including subcortical structures, which are implicated in many neurological, psychiatric, and neurodegenerative disorders (Fig. S49). Second, we replicate the meta-analysis results with the classic cytoarchitectonic atlas of Brodmann (138), which "remains one of the most widely used in neuroscience and continues to serve as a default reference system for reporting the locations of effects in current neuroimaging studies" (160, 161). We find that when queried in terms of Brodmann areas rather than sulcal-gyral anatomy, automated neuroscience experts still produce distinctly recognisable brain maps, bearing strong and significant spatial correspondence (all $rho > 0.80$) with the LLM maps obtained with the Desikan-Killiany atlas (Fig. S50). Likewise, results are also replicated with two more recent and much more fine-grained parcellations of the human cerebral cortex, which are defined from in vivo multi-modal neuroimaging rather than being based on anatomy or cytoarchitecture alone: the Brainnetome atlas, with 210 cortical regions defined from structural and functional MRI connectivity (256); and the Glasser atlas, with 360 regions defined from cortical thickness, myeloarchitecture, task activation and fMRI connectivity (160). For both atlases, the topography of the brain maps of literature summary generated by AI experts is significantly correlated with the corresponding NeuroSynth maps, beyond the effect of spatial autocorrelation (Fig. S52 and Fig. S54). We observe that the correlations with NeuroSynth meta-analytic maps are lower in magnitude for the Glasser atlas compared with those obtained from the Desikan-Killiany atlas (though still statistically significant), and region-function association scores obtained for the Glasser atlas are significantly lower than for the Desikan-Killiany atlas (Fig. S56). This result is expected, because the Glasser atlas is over 5-fold more fine-grained: 360 vs 68 regions, such that automated experts may find it less straightforward to clearly tell apart their

TABLE S1. Cortical regions, synonyms, and Desikan–Killiany (DK) ground-truth regions.

| Region | Synonyms | DK ground truth |
|---|---|---|
| Area V1 | Brodmann area 17; Primary visual cortex; Striate cortex; Area V1 | pericalcarine |
| Area V2 | Brodmann area 18; Secondary visual cortex; Prestriate cortex; Area V2 | cuneus; lingual; lateraloccipital |
| Area M1 | Brodmann area 4; Primary motor cortex; Area M1 | precentral |
| Area S1 | Brodmann area 1; Primary somatosensory cortex; Area S1 | postcentral |
| Area A1 | Brodmann areas 41 and 42; Primary auditory cortex; Heschl gyrus; Area A1 | transversetemporal |
| dlPFC | Brodmann areas 9 and 46; Dorsolateral prefrontal cortex | rostralmiddlefrontal |
| Broca area | Brodmann areas 44 and 45; Broca area | parsopercularis; parstriangularis |
| Wernicke area | Brodmann area 22; Wernicke area | superiortemporal; supramarginal; bankssts |

unique functional identities (as indeed we suspect that many human neuroscientists might). More fine-grained maps of literature summary are bound to be noisier: if a cortical territory can be subdivided into 5 regions, any evidence supporting the involvement of one of those regions for function X will also support the involvement of the superordinate territory, but the reverse does not hold. It is therefore expected – indeed, appropriate – that the region-function association scores provided by LLMs are lower for the more fine-grained atlas. In turn, correlating a noisier LLM map and a noisier NeuroSynth map is expected to produce a lower correlation, which is what we observe. Supporting this line of reasoning, we replicate the analysis using a coarser-grained version of the Glasser parcellation, which Glasser et al. (160) obtained by grouping their 180 unique regions into 22 cortical territories based on anatomical and functional considerations (e.g., 'Dorsal stream visual cortex'; Fig. S57a). Using this order-of-magnitude coarser version of the same multimodal parcellation reveals noticeably higher correlation values with the corresponding NeuroSynth maps (Fig. S57b,c).

It should be noted that the Brodmann areas are distinguished by number (e.g., 'Brodmann area 1', 'Brodmann area 17', etc; (161)) and nothing else – yet just as with human experts, this is enough information for AI experts to successfully tell apart their distinct functional roles. Additionally, 97 of the 180 areas in the Glasser atlas were not previously described in the literature (160). Nonetheless, querying the AI experts about the top functions of each Brodmann, Brainnetome, or Glasser area (rather than asking about one function at a time) replicates the main results in producing a distinctly unimodal-transmodal PC1, significantly aligned with the anatomical hierarchy from T1w:T2w ratio (Fig. S51, Fig. S53 and Fig. S55). Indeed, we emphasise that the focus of the present framework is not on specific individual regions in isolation, but rather on brain maps as a whole, where regions are contextualised with respect to each other.

Altogether, while the need for a parcellation with interpretable region names rather than voxelwise spatial resolution remains a limitation, we provide convergent evidence that the present framework and its results are not critically dependent on the use of the Desikan-Killiany atlas. On the contrary, results replicate across over an order of magnitude difference in spatial resolution, encompassing sulcal-gyral, classic cytoarchitectonic, and modern in vivo multimodal definitions of brain regions' identity. Hence, LLMs possess extensive neuroanatomical knowledge that transcends the use of one specific brain atlas.

*(x) Consistency and variability across different LLMs*

To what extent do different AI experts, trained in different ways on potentially different corpora of text, agree in their summaries of the brain mapping literature? For our main analyses we aggregated the opinions of top-tier models from leading AI companies (OpenAI's `GPT-5.4`, Anthropic's `Claude-4.6-Opus` and Google's `Gemini 3.1 Pro`) to ensure the best performance. Here, we complement our extensive benchmarking of different LLMs (Fig. S26) with a more in-depth assessment of the extent to which these proprietary LLMs agree with each other, and with three high-profile open-weight LLMs: OpenAI's `OSS-120b` and Google's `Gemma-4-31b-instruct` (representing the corresponding companies' flagship open-weight offerings), and Alibaba's `Qwen3-235b-a22b-2507`. We find that Qwen and Gemini have a tendency to produce generally higher region-function association scores, and OSS generally lower; finer-grained inspection further indicates that these trends vary across functions (Fig. S31).

We can also ask how LLMs' assessments of the same function vary across regions. In other words: are there regions where different automated experts, trained on vast but potentially non-overlapping corpora, disagree about involvement with a specific function? Indeed, we find that LLMs' disagreements (standard deviation across their region-function association score, for the same region and function) are not uniform but rather display clear and distinct regional patterns for each function (Fig. S30). Specifically, variability across LLMs is lower both for regions assigned consistently high region-function association scores (i.e. where LLMs are all confident that an association is present, such as pericalcarine area for vision), and for regions assigned consistently low values (such as motor cortex for memory) (Fig. S8 and Fig. S30). Variability among AI experts is instead higher for regions whose scores fall in the middle of the range (Fig. S8), as we might expect in the case of less conclusive evidence. Such cases where different LLMs diverge in their assessment of a region's involvement with a particular cognitive function may indicate deficiencies in the literature, providing the grounds for future targeted investigation.

Despite these differences, we find excellent agreement between the cognitive brain maps generated by different LLMs. For each pair of the 6 LLMs considered here, the mean correlation between the brain maps that they provide exceeds 0.70 (except for a few pairs in the range 0.50-0.60 involving the Qwen model), with `GPT-5.4` notably exceeding 0.9 similarity with most other models (Fig. S2). The same pattern is observed for macaque and mouse, with even higher agreement (Fig. S10). Thus, different AI experts provide highly aligned summaries of region-function association, indicating that the present results and the broader framework are not critically dependent on the specific LLMs or providers.

*(xi) Test-retest reliability across multiple API calls of the same LLM*

We also assess the replicability of results across repeated iterations of the workflow. To minimise test-retest differences, we set a fixed seed and the OpenRouter API is used to make outputs more focused and deterministic by setting the temperature to the most most conservative/reproducible temperature value (0.0). Nonetheless, it may not be possible to completely eliminate the inherent stochasticity of LLM outputs (257), and therefore we make it easy to quantify its impact through a `-retest` flag, which enables users to automatically request multiple iterations of the prompt.

For each of the 8 cognitive domains, we run the same prompt 10 times for each of the 3 LLMs from our main analyses and their chosen open-weight counterparts. We show that standard deviation across iterations is very low for all models and functions, ranging between 0.001 and 0.05 (Fig. S58a). Consequently, we find that the most conservative and most liberal maps generated by each LLM for each function (i.e., the maps of minimum and maximum region-function association) are all strongly and significantly correlated: almost all exceed 0.70 in magnitude, with `GPT-5.4`, `Claude-4.6-Opus`, and `Gemma-4-31b` all in the range of 0.94 to 0.99 (Fig. S58b). These analyses indicate that results are highly consistent not only across LLMs, but also within each individual LLM. Altogether, here we combined the outputs of multiple individually high-performing models as a well documented strategy to boost LLMs' factual accuracy (258–260). Reassuringly, our assessment provides confidence in the neuroscientific expertise of individual LLMs, being stable over time and consistent across different AI experts.

**Supplementary Figures**

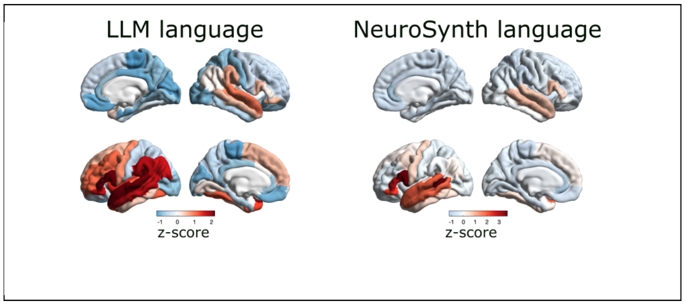


Figure S1. **Bilateral maps of language from LLMs and NeuroSynth |** **(Left)** LLM map of region-function association scores for 'language'. **(Right)** NeuroSynth map for 'language'. Both display the well-known left-lateralisation characteristic of language. Brain maps are displayed on the lateral and medial surface of the right (top) and left (bottom) cortical hemispheres.

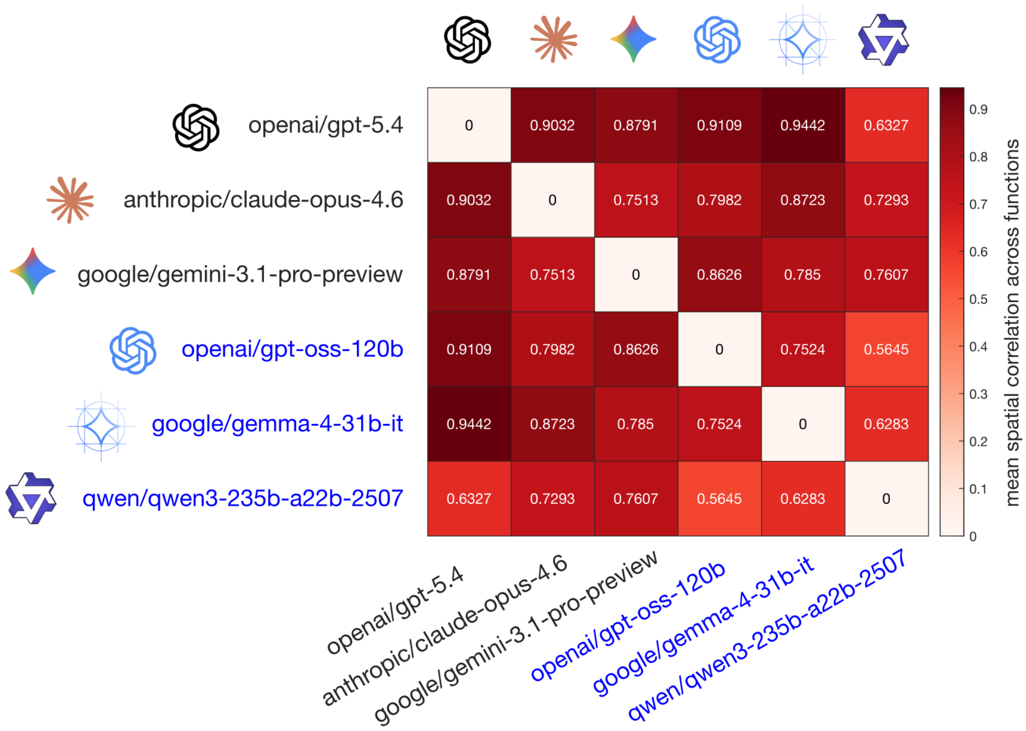


Figure S2. **Similarity of human brain maps generated by different LLMs |** Spatial correlation between brain maps generated by six prominent LLMs from the Claude, GPT, OSS, Gemini, Gemma, and Qwen model families, averaging across 8 representative domains of cognition.

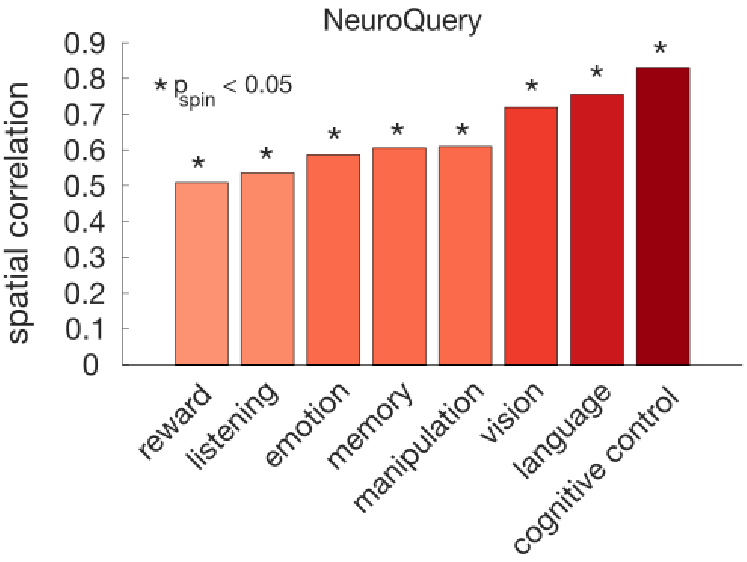


Figure S3. **LLM-generated brain maps of cognition correlate with NeuroQuery maps |** Correlation between meta-analytic maps and LLM-generated maps is replicated when using NeuroQuery instead of NeuroSynth to summarise the human neuroimaging literature. The brain maps generated by NeuroQuery are *predictions of the probability* that a study on the cognitive operation of interest will report observations at each location (13). Significance of correlations is assessed against a population of null maps with preserved spatial autocorrelation.

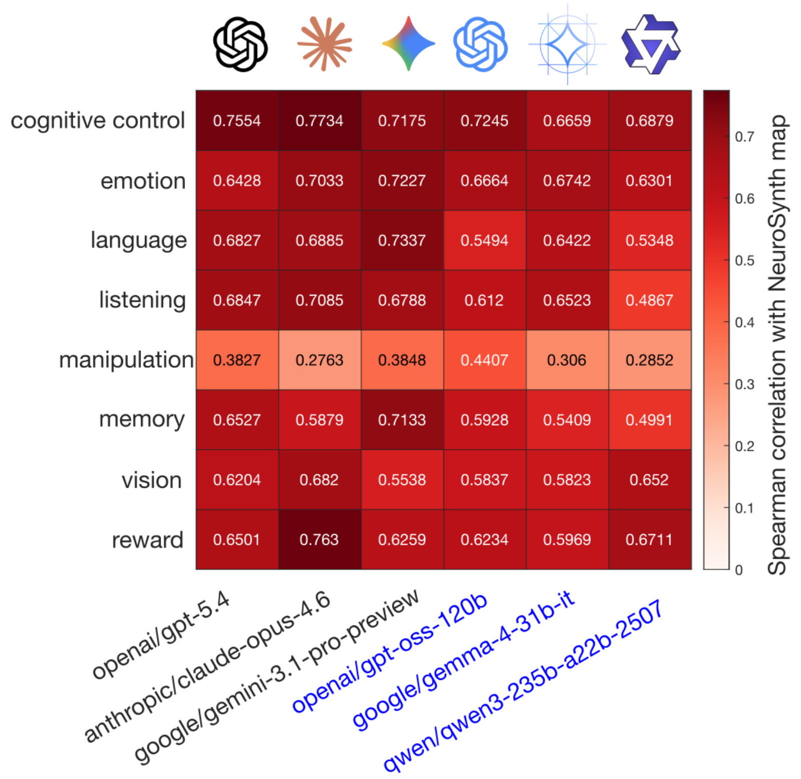


Figure S4. **Individual LLMs generate human brain maps that match NeuroSynth |** Spatial correlation between brain maps for 8 representative domains of cognition generated by NeuroSynth, with corresponding brain maps generated by the three frontier LLMs used in the main analysis (GPT, Claude, Gemini) and by three additional open-weight LLMs used for replication (OSS, Gemma, and Qwen).

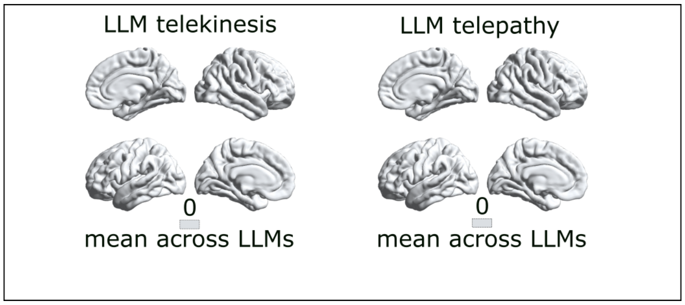


Figure S5. **Negative control with non-existent functions | (Left)** Region-function association scores obtained upon prompting AI experts for the function 'telekinesis'. **(Right)** Region-function association scores obtained upon prompting AI experts for the function 'telepathy'. LLMs return all-zero maps for both functions, consistently with their lack of support in the scientific literature. Brain maps are displayed on the lateral and medial surface of the right (top) and left (bottom) cortical hemisphere.

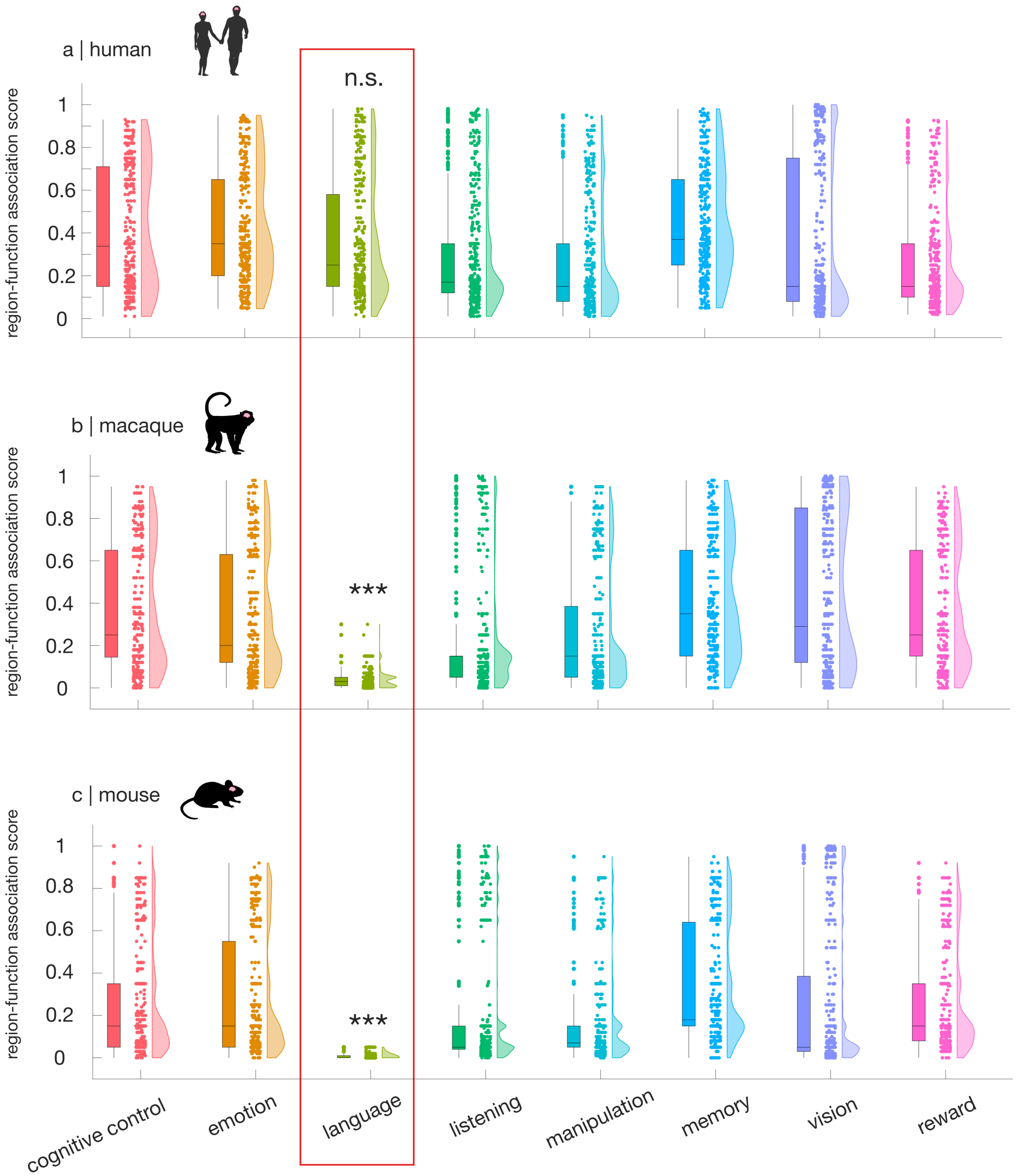


Figure S6. **LLM-estimated region-function association scores for “language” in macaque and mouse is significantly lower than any other cognitive domain, unlike in human | (a)** Human: language mean = 0.35, SD = 0.27; other terms mean = 0.34, SD = 0.28; t(3262) = 0.47, p = 0.633, Hedge’s $g$ = 0.025 from independent-samples t-test between ‘language’ and all other terms combined. **(b)** Macaque: language mean = 0.035, SD = 0.037; other terms mean = 0.34, SD = 0.30; t(3934) = -22.11, $p < 0.001$, Hedge’s $g$ = -1.06 from independent-samples t-test between ‘language’ and all other terms combined. **(c)** Mouse: language mean = 0.0057, SD = 0.0129; other terms mean = 0.25, SD = 0.28; t(3454) = -17.85, $p < 0.001$, Hedge’s $g$ = -0.92 from independent-samples t-test between ‘language’ and all other terms combined. For each species, data are pooled across all regions and all LLMs. Each data-point represents the region-function association score attributed by one LLM to one region. Box-plots: center line, median; box limits, upper and lower quartiles; whiskers, $1.5\times$ interquartile range. ***, $p < 0.001$.

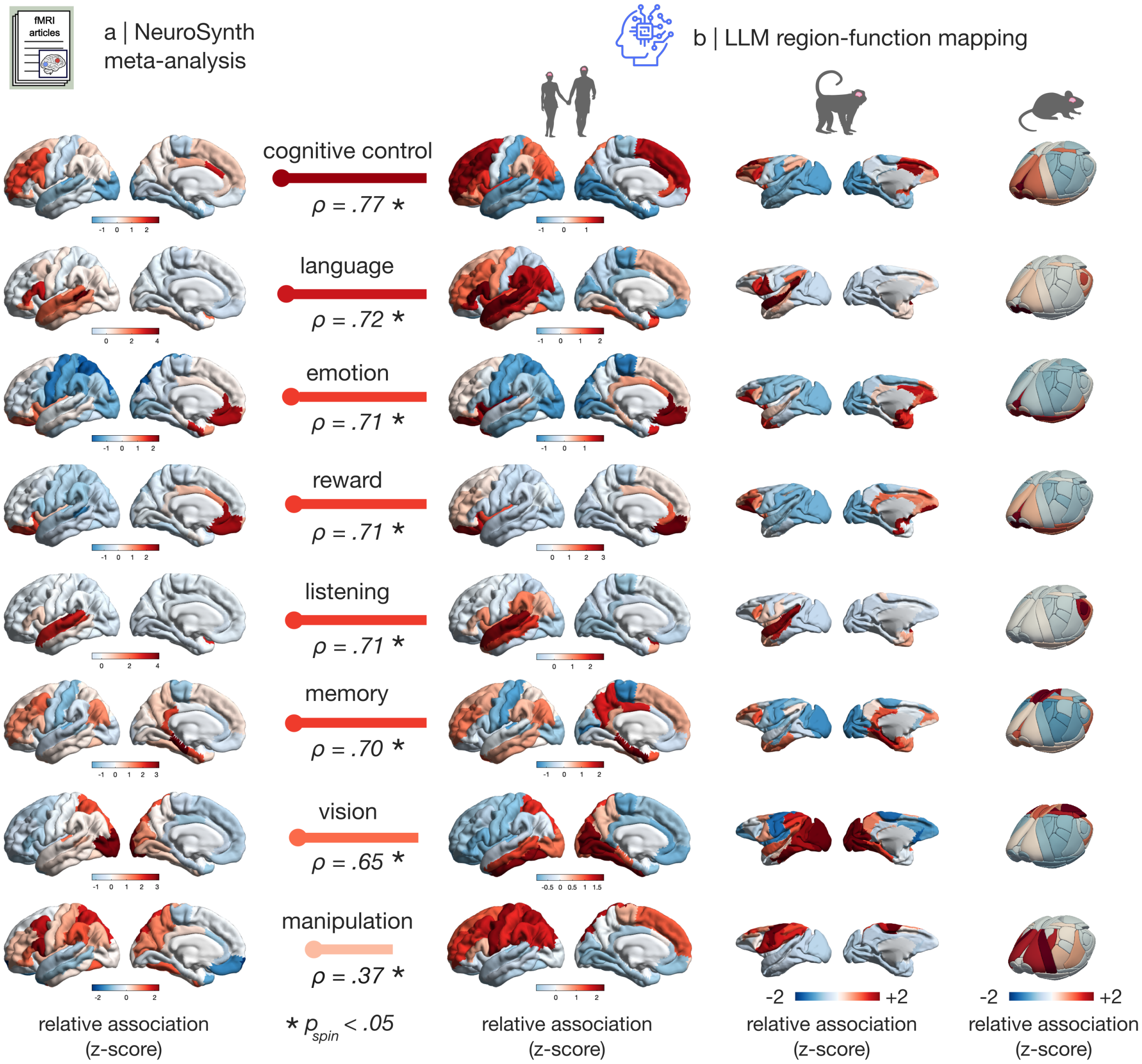


Figure S7. **LLMs generate brain maps of cognition, z-scored prior to averaging across LLMs | (a)** NeuroSynth meta-analytic maps term-based meta-analysis of the fMRI literature) for 8 representative domains of cognition identified through data-driven computational ontology (69)). Brain maps show, for each region, NeuroSynth's measure of association strength between terms and voxels, quantified as the z-score from a two-way ANOVA testing for the presence of a non-zero association between term use and voxel activation (12). Note that NeuroSynth does not distinguish between areas that are activated or deactivated in relation to the term of interest, and a negative number does not imply deactivation or inhibition. To enable comparison, the voxelwise NeuroSynth maps are parcellated by averaging the values of the voxels belonging to the same region of the Desikan-Killiany atlas (see Methods). **(b)** AI experts generate plausible brain maps of cognition for human, macaque and mouse. For each term and species, a summary map is obtained by first z-scoring across regions the region-function association scores assigned by different LLMs (here: `GPT 5.4`; `Gemini 3.1 Pro`; and `Claude 4.6 Opus`), and then averaging the z-scored maps of each term across LLMs. See Fig. 1 for the same figure, but with LLM-generated brain maps averaged without prior z-scoring; all 8 LLM-generated maps have Spearman's $r > 0.99$ between the z-scored and non z-scored versions. Human: 68 cortical regions from the Desikan-Killiany atlas (70). Macaque: 82 cortical regions from the Regional Mapping atlas (71). Mouse: 72 cortical regions from the Allen common coordinate framework atlas (72). Horizontal bars indicate the Spearman correlation between the brain maps displayed at either side, with length proportional to the correlation magnitude. Scatter-plots for each correlation are shown in Fig. S8. Significance of correlations is assessed against a null population of surrogate maps with preserved spatial autocorrelation. See Fig. S9 for the corresponding null distributions. For both (a) and (b), brain maps are displayed on the lateral and (for human and macaque) medial surface of the species-specific left cortical hemisphere.

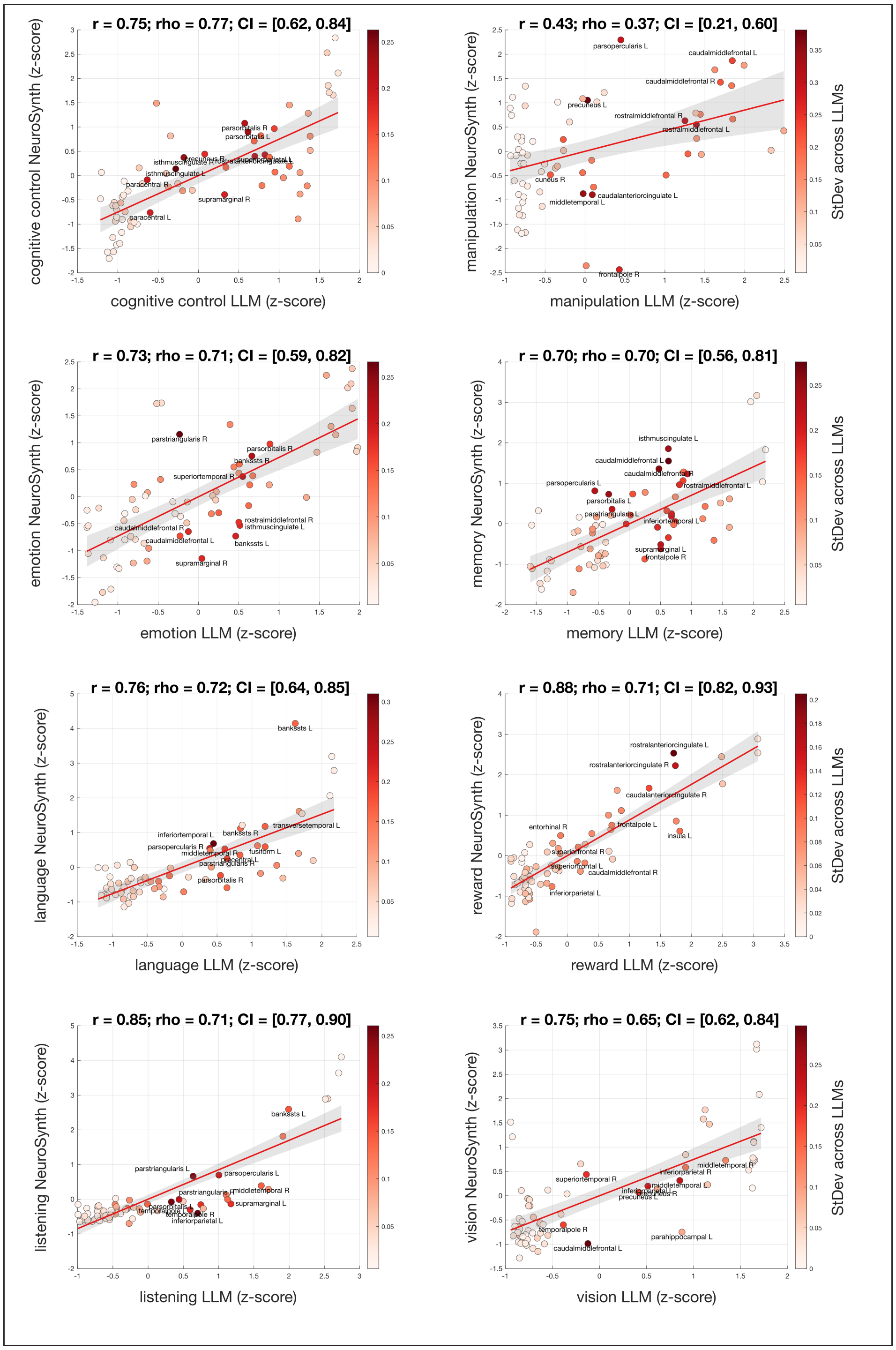


Figure S8. **Spatial correlations between LLM-generated and NeuroSynth-generated maps of cognition |** Pearson and Spearman correlations are shown for each comparison. See Fig. S9 for the corresponding p-values controlling for spatial autocorrelation. Each data-point is one brain region of the Desikan-Killiany atlas, with colour indicating the standard deviation across AI experts for that region and function. Note how higher standard deviations tend to occur towards the middle rather than at the extremes.

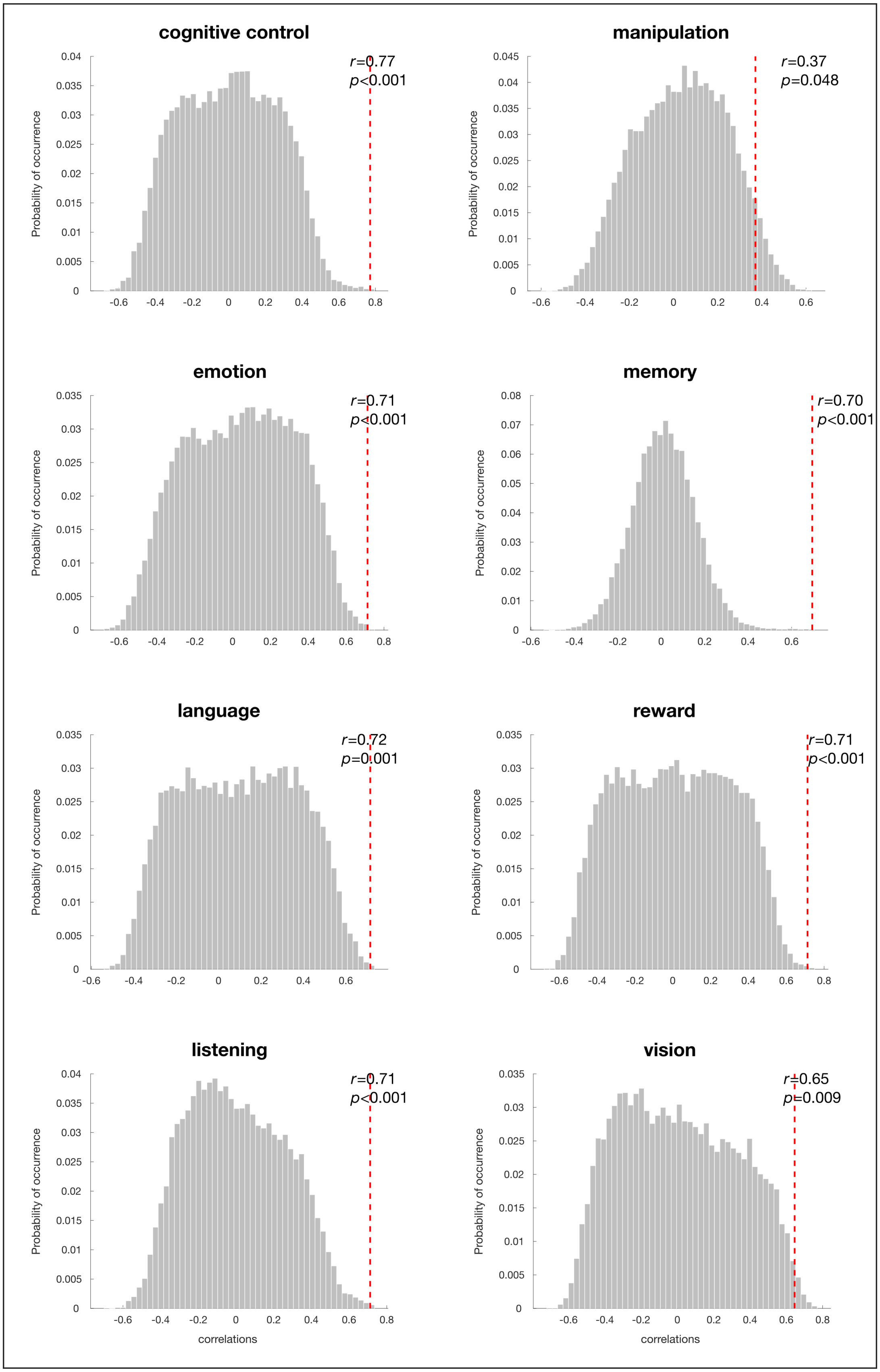


Figure S9. **Brain maps of cognition generated by AI experts correlate with corresponding NeuroSynth maps beyond the effect of spatial autocorrelation |** Red vertical bar: empirical correlation between NeuroSynth map and LLM map. Histograms show the distribution of null values obtained from a population of surrogate maps with preserved spatial autocorrelation.

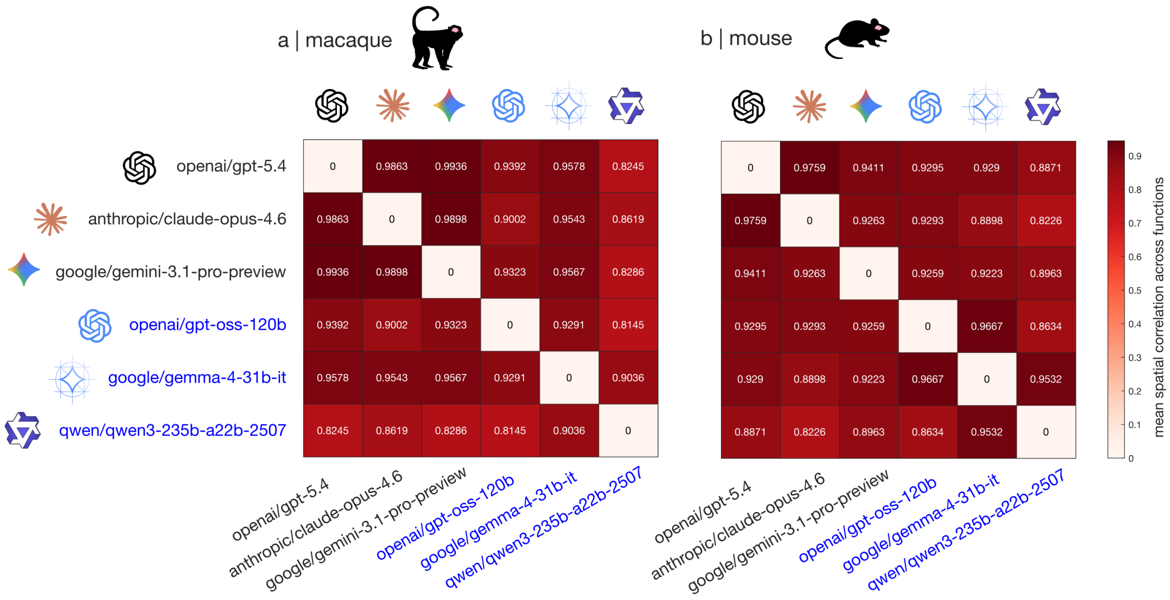


Figure S10. **Similarity of macaque and mouse brain maps generated by different LLMs |** Spatial correlation between brain maps generated by six prominent LLMs from the Claude, GPT, OSS, Gemini, Gemma, and Qwen model families, averaging across 8 representative domains of cognition. **(a)** Macaque. **(b)** Mouse.

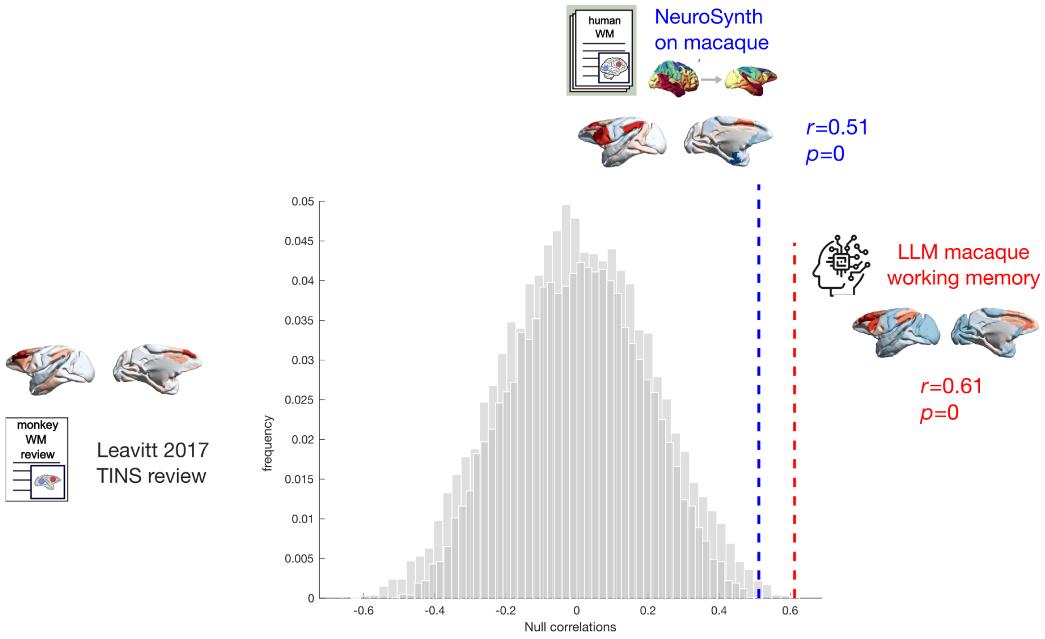


Figure S11. **LLM-derived map of macaque working memory outperforms NeuroSynth-derived map of human working memory projected to macaque brain |** Red vertical bar: empirical correlation between LLM map, and literature summary map of macaque working memory from Leavitt et al. (73). Blue vertical bar: empirical correlation between human NeuroSynth map of working memory projected onto macaque brain, and literature summary map of macaque working memory from Leavitt et al. (73). Histograms show the distribution of null values obtained from a population of surrogate maps with preserved spatial autocorrelation. Brain maps are displayed on the lateral and medial surface of the left macaque cortical hemisphere.

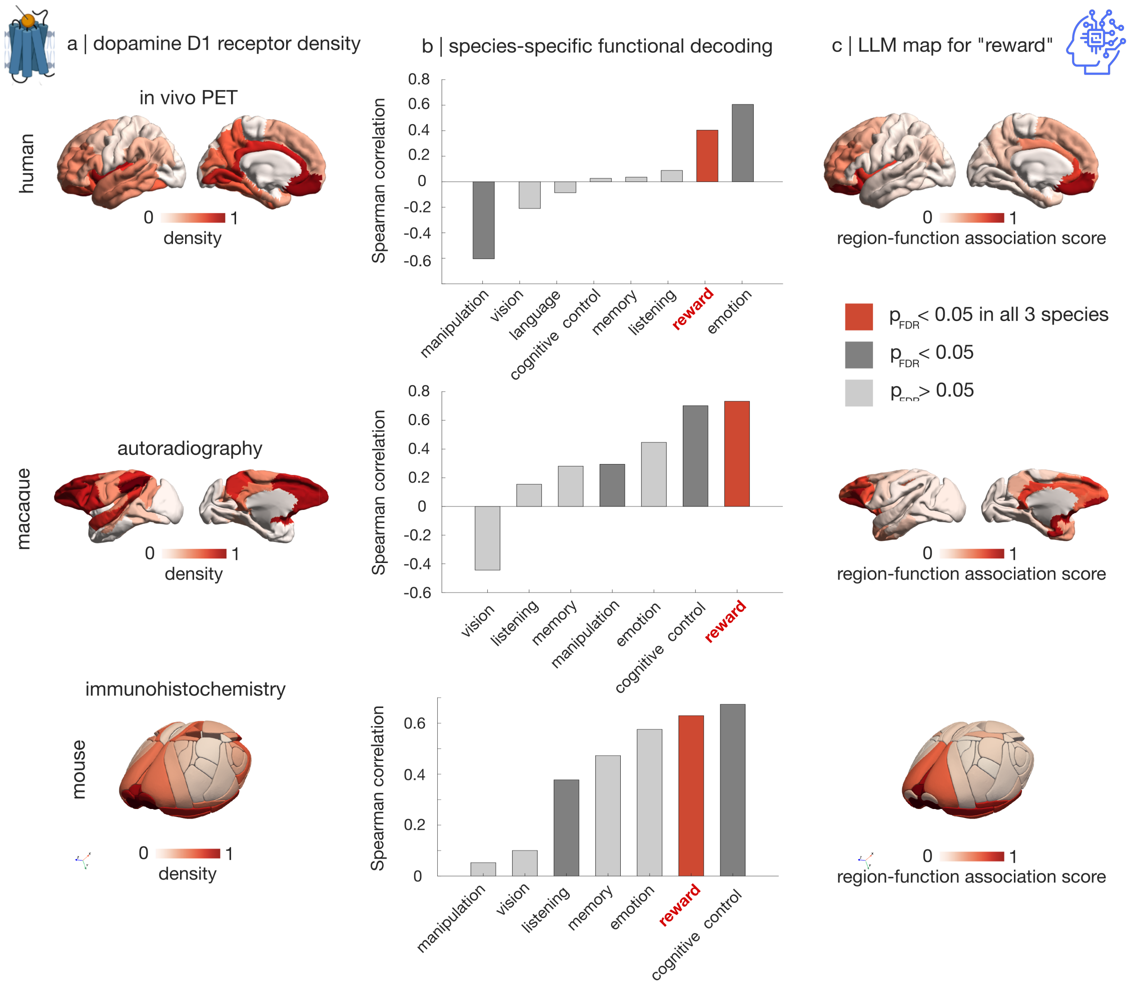


Figure S12. **Decoding species-specific brain maps of D1 receptor distribution with LLM-generated brain maps | (a)** Human D1 receptor map is obtained from in vivo Positron Emission Tomography (88). Macaque D1 receptor map is obtained from in vitro autoradiography (59). Mouse D1 receptor map is obtained from immunohistochemistry (89). **(b)** For each species, the D1 receptor map is spatially correlated against maps of canonical cognitive domains. Significance of correlations is assessed against a population of species-specific null maps with preserved spatial autocorrelation, followed by FDR correction. Within each species, terms that are both SA- and FDR-significant are shown in darker grey. Only one map is consistently significantly associated with D1 receptor distribution across all three species (shown in orange), corresponding to 'reward'. **(c)** Species-specific LLM-generated maps of region-function association for 'reward'. For both (a) and (c), brain maps are displayed on the lateral and (for human and macaque) medial surface of the left cortical hemisphere.

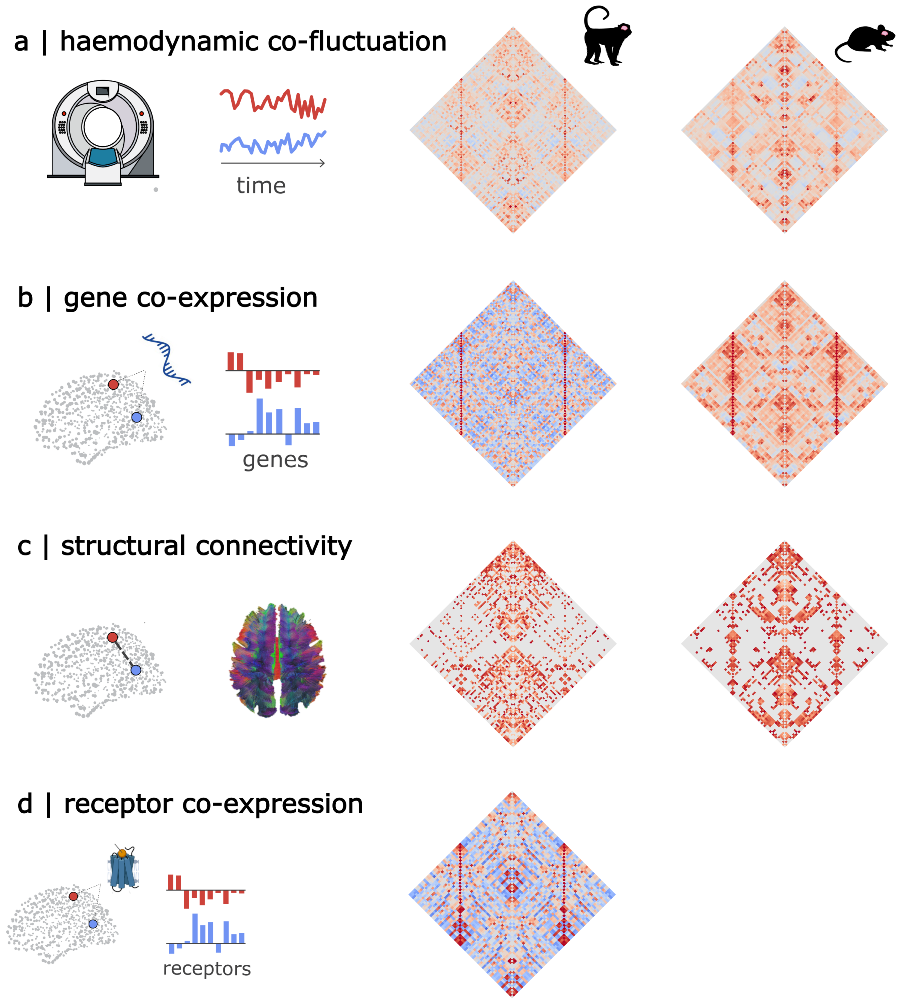


Figure S13. **Networks of structural and functional interactions for macaque and mouse | (a)** Co-fluctuation (correlation over time) of spontaneous activity from fMRI haemodynamics. **(b)** Correlated patterns of gene expression from stereo-seq (macaque) and in situ hybridization (mouse). **(c)** Structural connectivity from combined diffusion tractography and tract-tracing (macaque) and tract-tracing (mouse). **(d)** Correlated patterns of receptor density from in vitro autoradigraphy (macaque only). Note that macaque receptor density is not available for all regions, and regions without receptor expression were therefore excluded from the analysis.

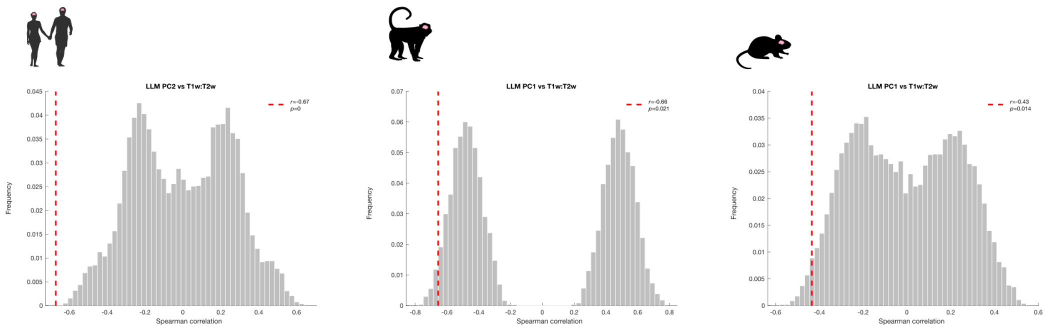


Figure S14. **Significant correlations between principal components of LLM-derived functional similarity, and T1w:T2w ratio across species | (Left)** Human PC2. **(Middle)** Macaque PC1. **(Right)** Mouse PC1. Histograms show the distribution of null values obtained from a population of surrogate maps with preserved spatial autocorrelation.

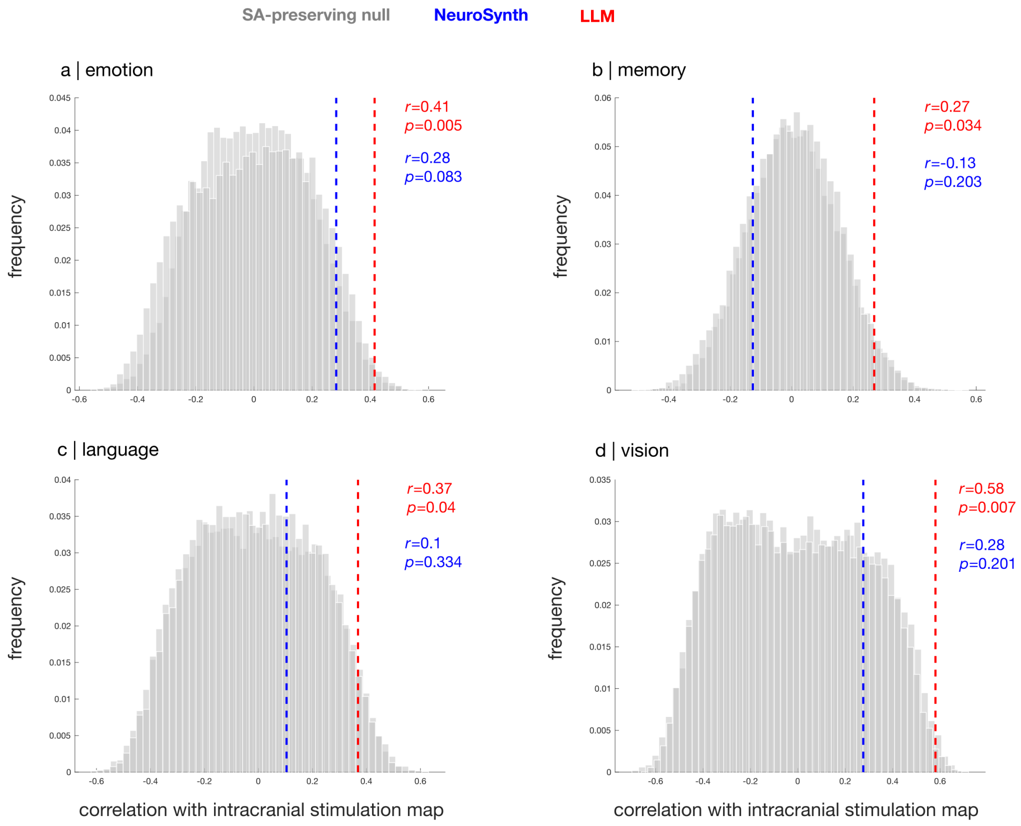


Figure S15. **LLM-derived maps outperform NeuroSynth for recapitulating circuits obtained from direct intracranial stimulation | (a)** Emotion. **(b)** Memory. **(c)** Language. **(d)** Vision. Red vertical bar: empirical correlation between iES map and LLM map. Blue vertical bar: empirical correlation between iES map and NeuroSynth map. Histograms show the distribution of null values obtained from a population of surrogate maps with preserved spatial autocorrelation.

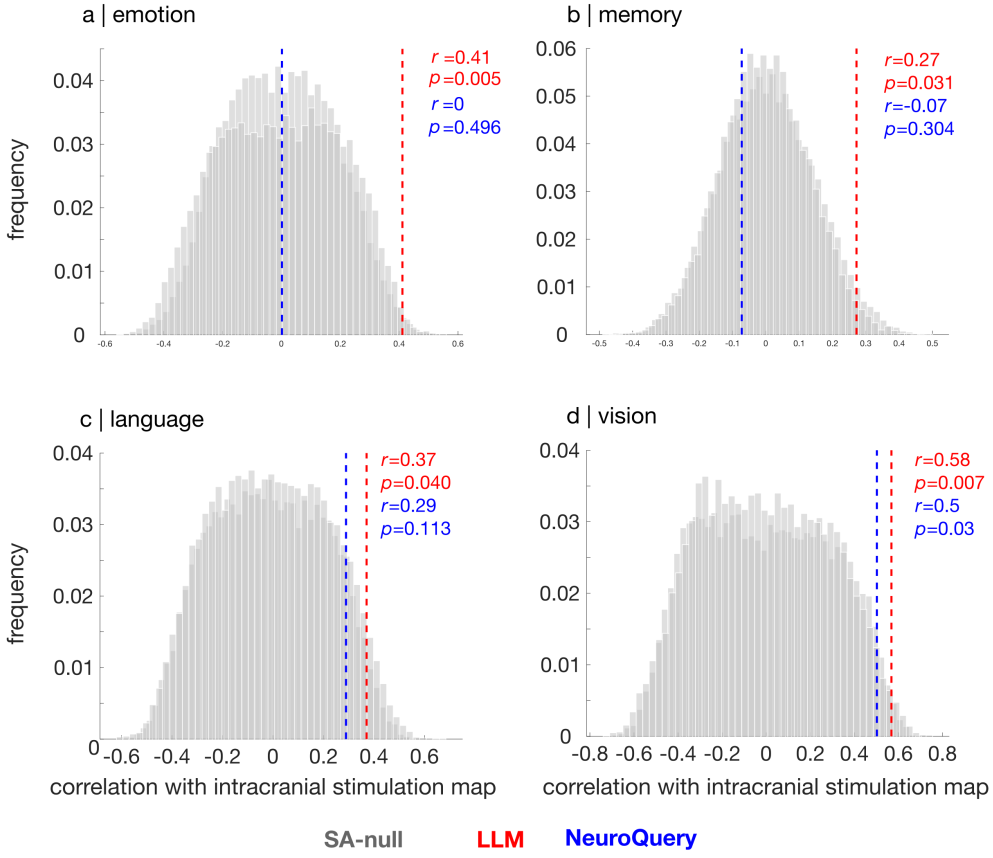


Figure S16. **LLM-derived maps outperform NeuroQuery for recapitulating circuits obtained from direct intracranial stimulation | (a)** Emotion. **(b)** Memory. **(c)** Language. **(d)** Vision. Red vertical bar: empirical correlation between iES map and LLM map. Blue vertical bar: empirical correlation between iES map and NeuroQuery map. Histograms show the distribution of null values obtained from a population of surrogate maps with preserved spatial autocorrelation.

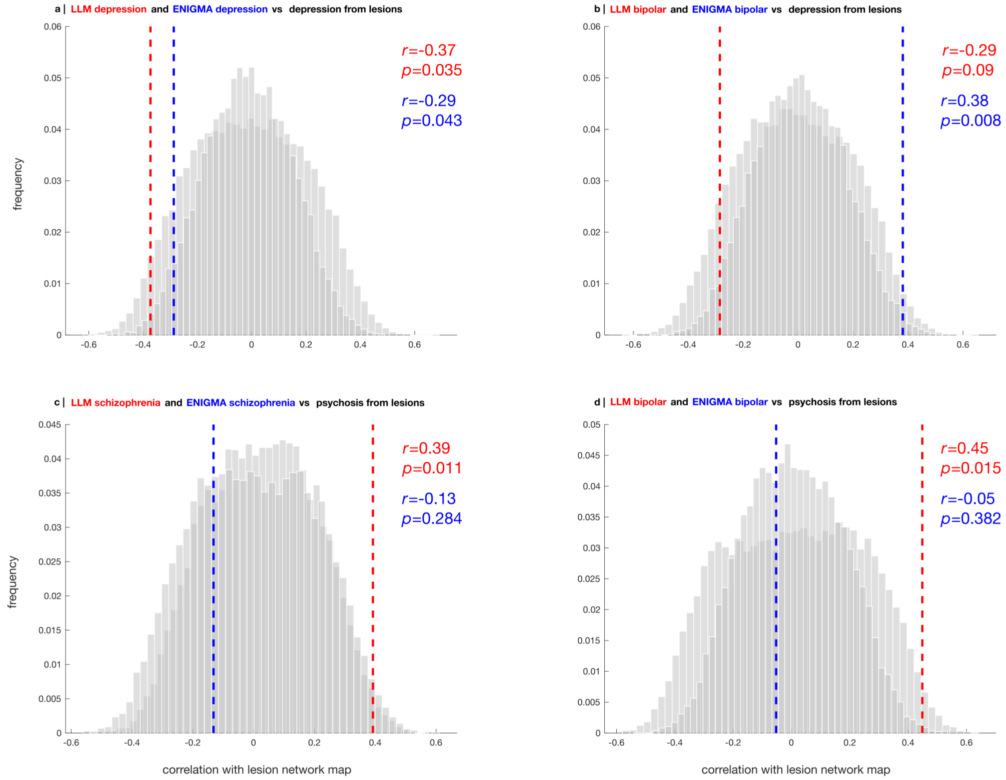


Figure S17. **LLM-derived maps outperform ENIGMA maps of absolute cortical thickness change for recapitulating circuits obtained from lesion network mapping | (a)** Depression maps vs depression lesion circuit. **(b)** Bipolar maps vs depression lesion circuit. **(c)** Schizophrenia maps vs psychosis lesion circuit. **(d)** Bipolar maps vs psychosis lesion circuit. Red vertical bar: empirical correlation between lesion circuit map and LLM map. Blue vertical bar: empirical correlation between lesion circuit map and ENIGMA map (magnitude of case-control effect size). Histograms show the distribution of null values obtained from a population of surrogate maps with preserved spatial autocorrelation.

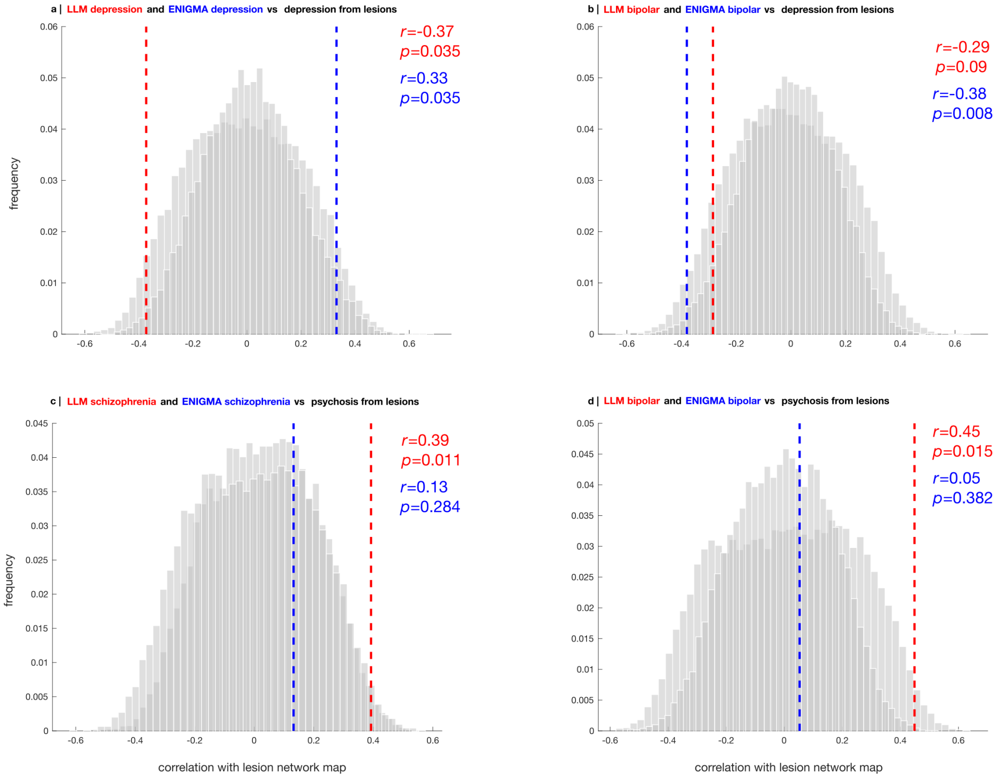


Figure S18. **LLM-derived maps outperform ENIGMA maps of cortical thickness change (Cohen's *d*) for recapitulating circuits obtained from lesion network mapping | (a)** Depression maps vs depression lesion circuit. **(b)** Bipolar maps vs depression lesion circuit. **(c)** Schizophrenia maps vs psychosis lesion circuit. **(d)** Bipolar maps vs psychosis lesion circuit. Red vertical bar: empirical correlation between lesion circuit map and LLM map. Blue vertical bar: empirical correlation between lesion circuit map and ENIGMA map (case-control effect size). Histograms show the distribution of null values obtained from a population of surrogate maps with preserved spatial autocorrelation.

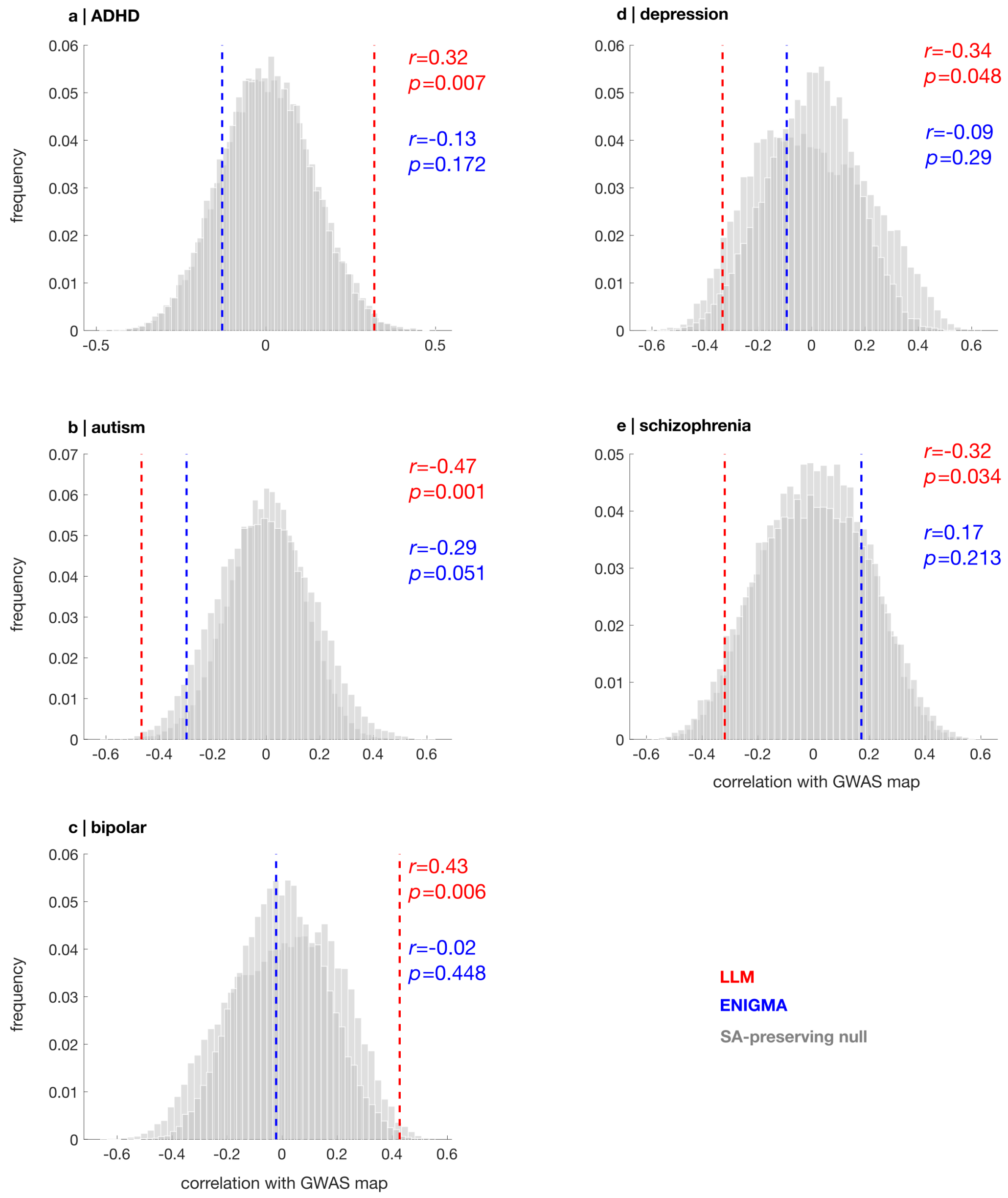


Figure S19. **LLM-derived maps outperform ENIGMA maps of absolute cortical thickness change for recapitulating risk-gene maps from GWAS | (a)** ADHD **(b)** Autism. **(c)** Bipolar. **(d)** Depression. **(e)** Schizophrenia. Red vertical bar: empirical correlation between risk-gene expression map and LLM map. Blue vertical bar: empirical correlation between risk-gene expression map and ENIGMA map (magnitude of case-control effect size). Histograms show the distribution of null values obtained from a population of surrogate maps with preserved spatial autocorrelation.

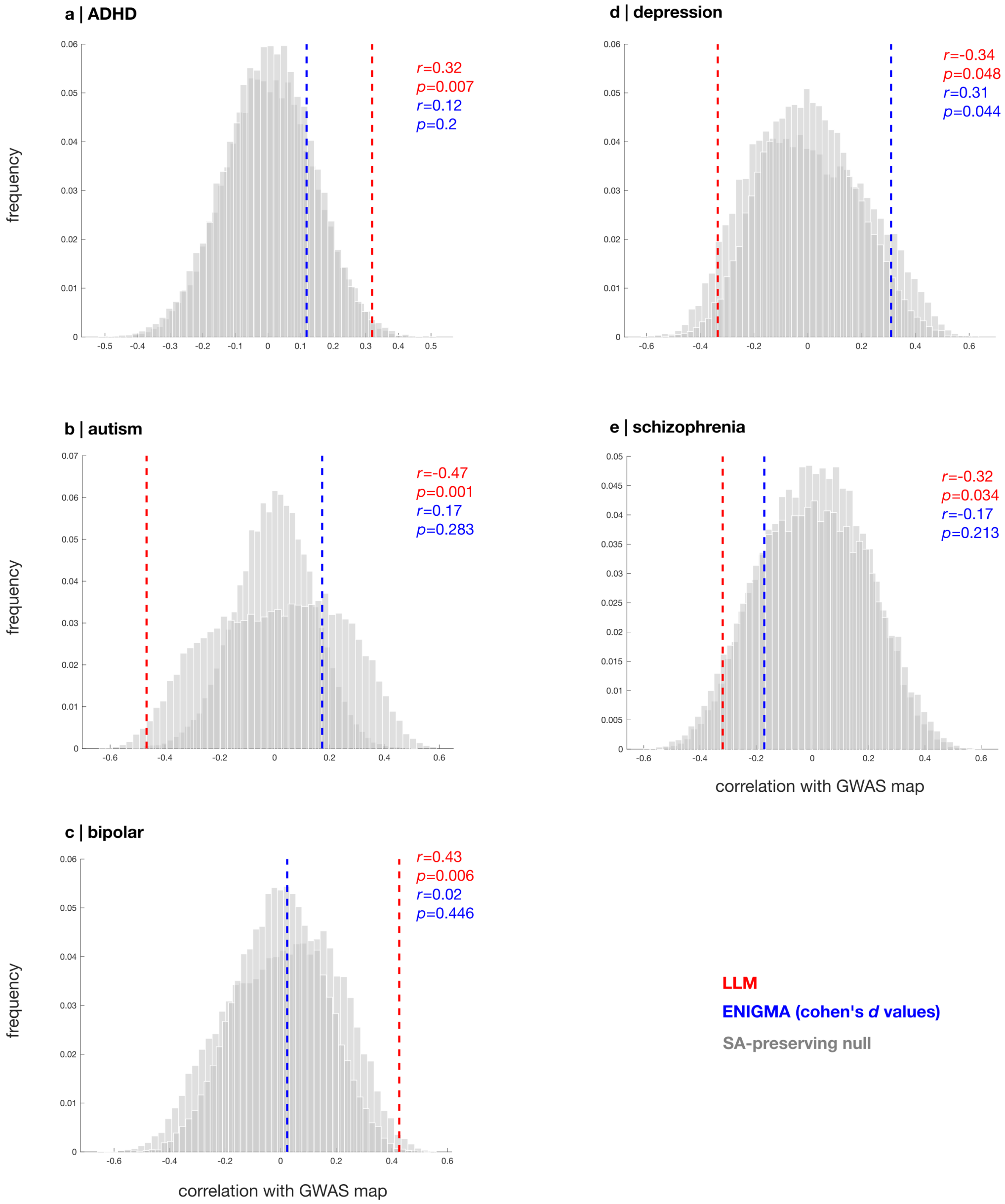


Figure S20. **LLM-derived maps outperform ENIGMA maps of cortical thickness change (Cohen's *d*) for recapitulating risk-gene maps from GWAS | (a)** ADHD **(b)** Autism. **(c)** Bipolar. **(d)** Depression. **(e)** Schizophrenia. Red vertical bar: empirical correlation between risk-gene expression map and LLM map. Blue vertical bar: empirical correlation between risk-gene expression map and ENIGMA map (case-control effect size). Histograms show the distribution of null values obtained from a population of surrogate maps with preserved spatial autocorrelation.

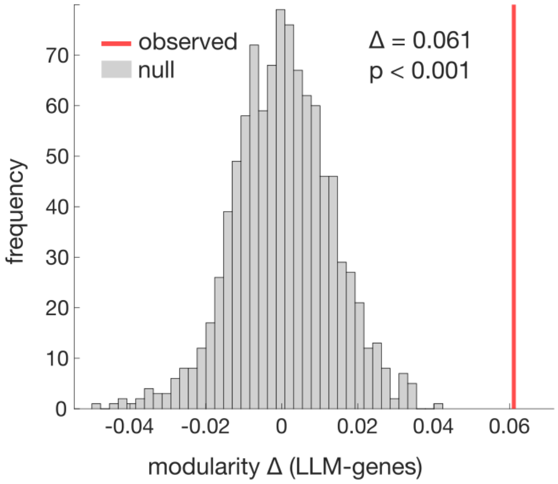


Figure S21. **Clustering of disease maps from LLMs aligns with symptom dimensions better than transcriptomically derived maps of molecular risk |** Modularity is computed using the Newman-Girvan algorithm (252, 253). The observed difference in modular structure is significantly greater than what would be observed by chance, as assessed using a label-permutation null model whereby the difference in modularity score is recomputed after randomly rearranging the matrix (5,000 nulls).

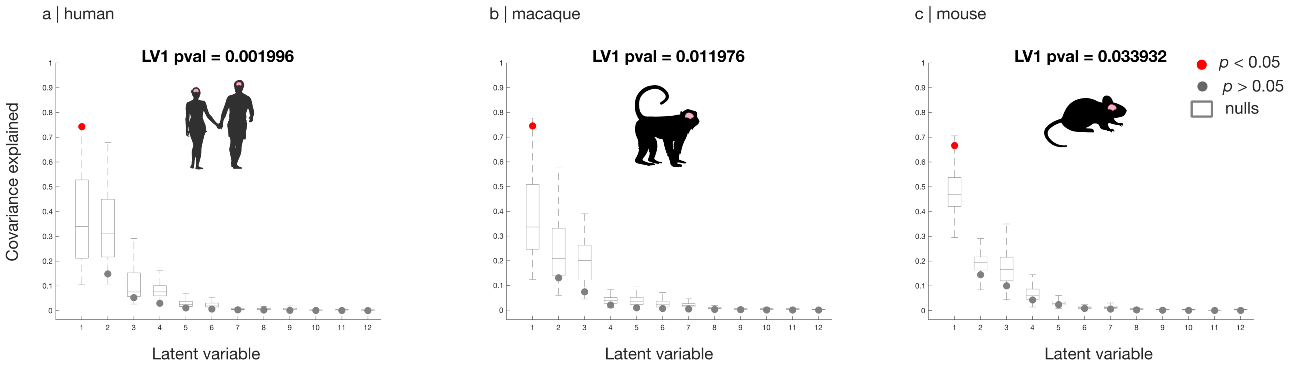


Figure S22. **AI-generated species-specific cognitive maps are significantly associated with species-specific transcriptomic patterns | (a)** Significant PLS LV1 in human. **(b)** Significant PLS LV1 in macaque. **(c)** Significant PLS LV1 in mouse. In each species, significance is assessed against a null distribution of species-specific spatial autocorrelation-preserving null maps (Methods).

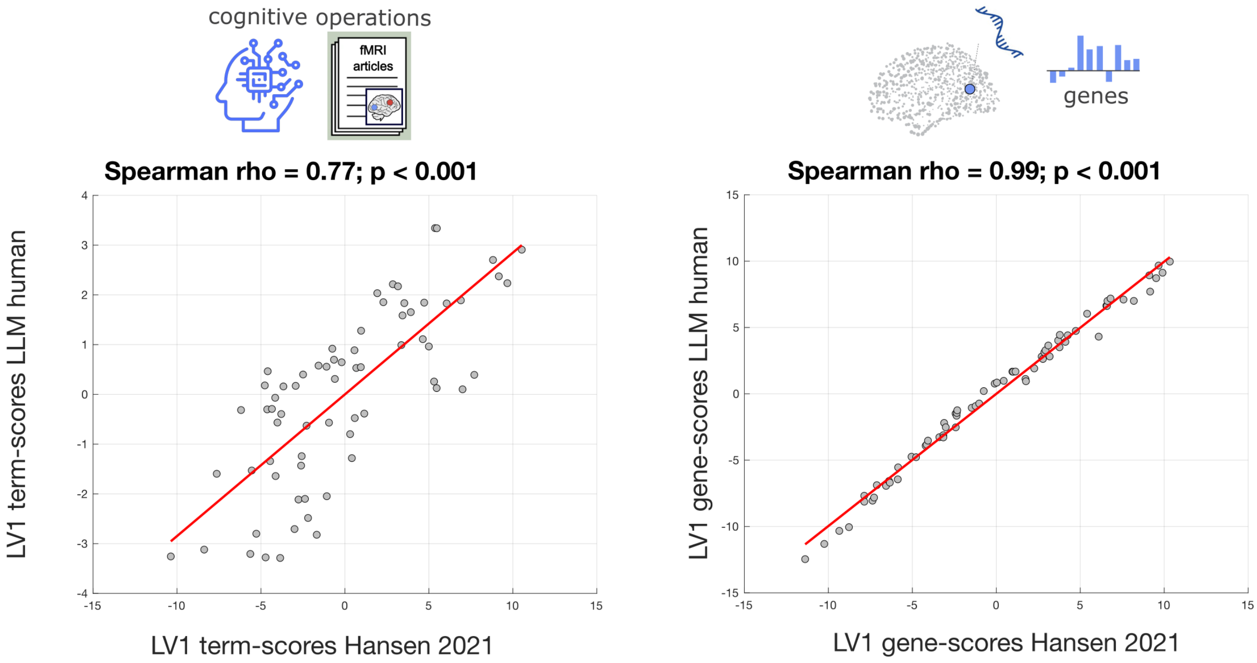


Figure S23. **AI-generated cognitive maps recapitulate gene-cognition associations obtained using NeuroSynth | (Left)** Significant correlation between PLS LV1 term scores obtained from NeuroSynth (as per (131); abscissa) and from AI experts (ordinate). **(Right)** Significant correlation between PLS LV1 gene scores obtained from NeuroSynth (as per (131); abscissa) and from AI experts (ordinate). Each data-point is one brain regions from the Desikan-Killiany atlas.

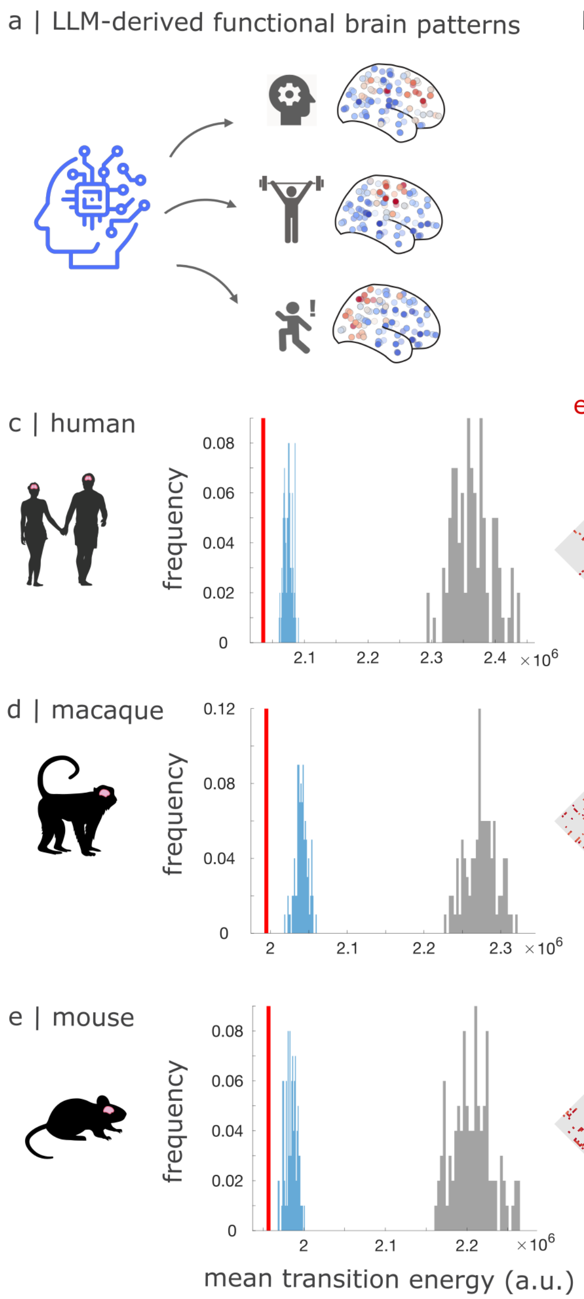


Figure S24. **AI experts generalise the energy efficiency of human brain networks to macaque and mouse | (a)** We use LLMs to obtain brain maps pertaining to 17 representative cognitive operations in human, macaque, and mouse. **(b)** We apply principles of network control theory with species-specific connectomes to compute the energy required to transition between each pair of cognitive brain maps. **(c)** Consistently across all three species, the mean transition energy across all pairs is significantly lower for the real connectome than for geometry-preserving rewired networks, which in turn outperform degree-preserving nulls.

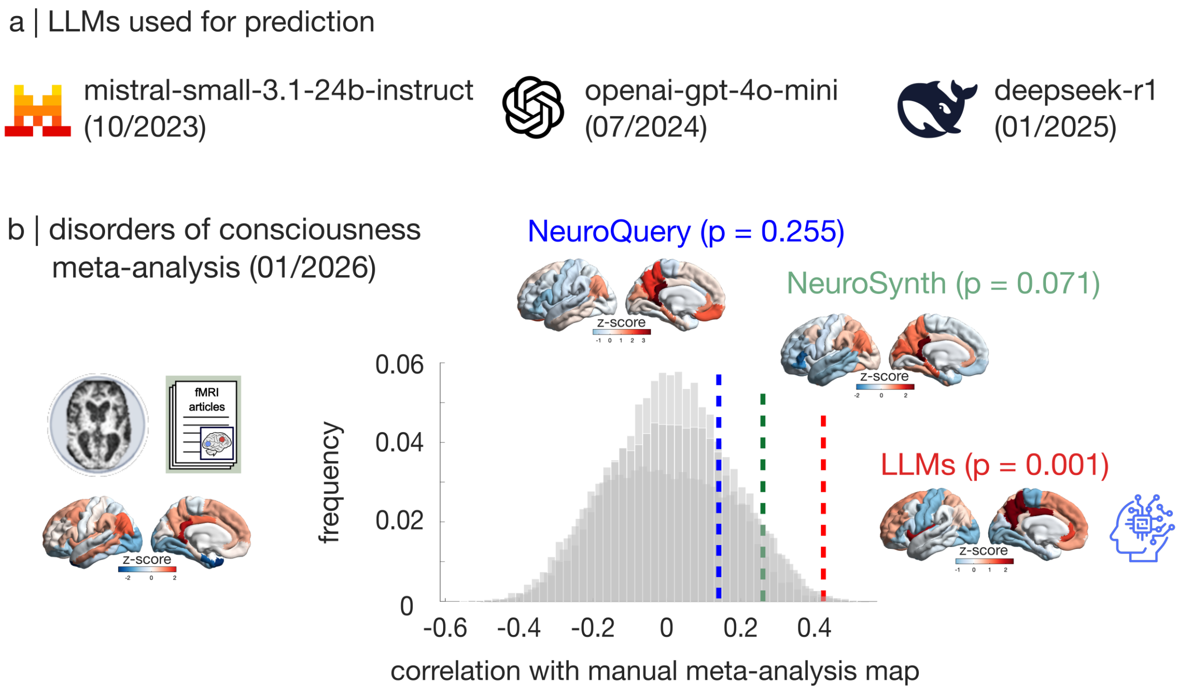


Figure S25. **LLM-generated brain maps predict unseen manual meta-analyses |** **(a)** LLMs used for prediction. Knowledge cutoff dates ranging from October 2023 to January 2025, prior to the manual meta-analysis becoming available online. **(b)** LLM-generated map of 'consciousness' is significantly spatially correlated with the manual meta-analysis of disorders of consciousness from (136) (Spearman's $r = 0.42, p = 0.001$), outperforming the map of 'consciousness' obtained from both NeuroQuery (Spearman's $r = 0.14, p = 0.255$) and from NeuroSynth (Spearman's $r = 0.26, p = 0.071$). Significance of correlations is assessed against a population of null maps with preserved spatial autocorrelation. Brain maps are z-scored across regions, and displayed on the lateral and medial surface of the left cortical hemisphere.

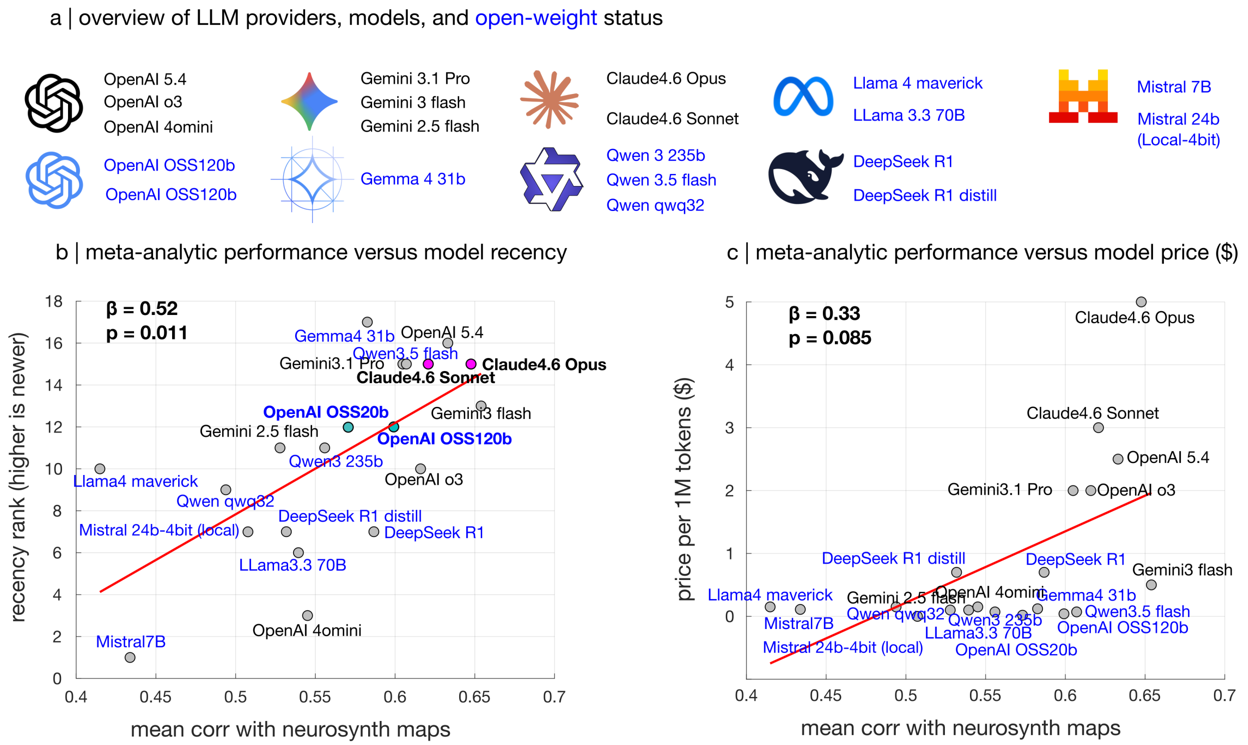


Figure S26. **Benchmarking meta-analytic performance against cost and recency across 20 models from 7 leading AI companies | (a)** Overview of LLM providers, models, and open-weight status. **(b)** Meta-analytic performance versus model recency (rank: highest is newest; see Table 1 for details). Performance is quantified as the mean spatial correlation with NeuroSynth maps, for the 8 cognitive domains of Beam et al. (69). **(c)** Meta-analytic performance versus model price ($ per million tokens). Note that `OSS-20b` is outperformed by the larger `OSS-120b`, and a very similar performance difference is also observed between `Claude-4.6-Sonnet` and the larger `Claude-4.6-Opus`, showing that model size does matter for performance when comparing models of the same family and recency. However, note also that both `OSS-20b` and `Gemma-4-31b-instruct` outperform the larger but older `llama-3.3-70B` - confirming that advances in training and architecture translate to improved performance.

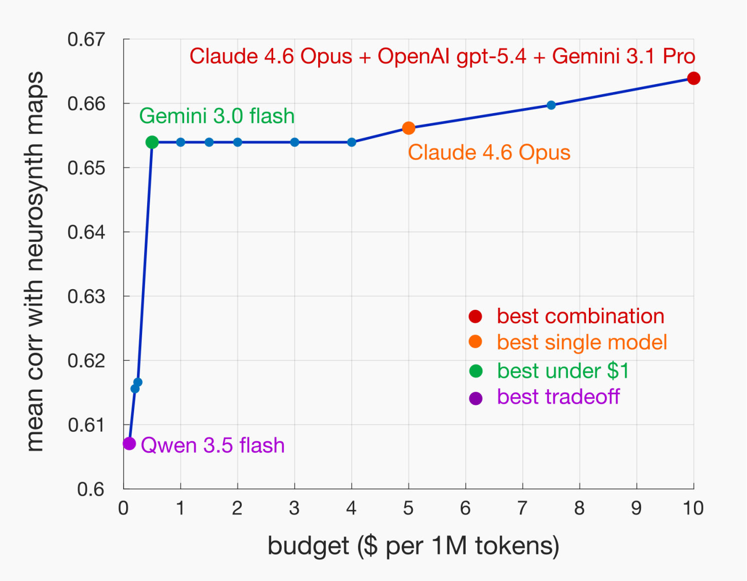


Figure S27. **Maximum achievable meta-analytic performance across budgets |** For all possible combinations of 20 proprietary and open-weight models considered here (Table 1), we ask if aggregating (averaging) their maps improves the match with corresponding NeuroSynth maps. We consider models whose cost per 1M tokens adds up to $0.10, $0.20, $0.25, $0.50, $1.0, $1.50, $2.0, $3.0, $4.0, $5.0, $7.50, or $10.0.

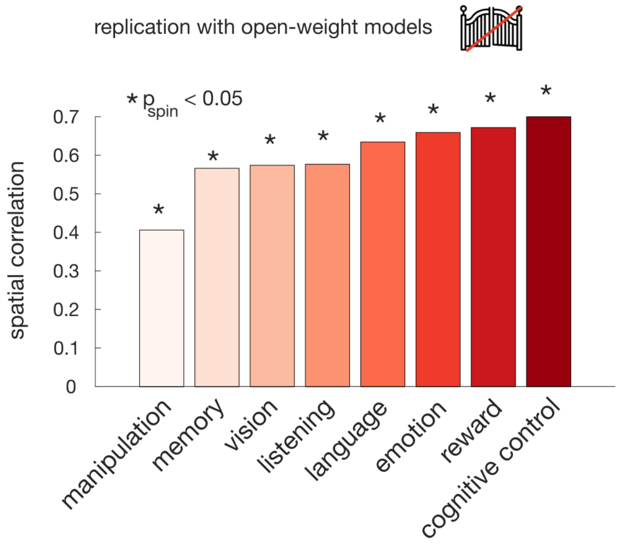


Figure S28. **Brain maps of cognition generated by open-weight LLMs correlate with NeuroSynth maps |** We replicate the main results using three frontier open-weight models: OpenAI's `OSS-120b`, Google's `Gemma-4-31b-instruct`, and Alibaba's `Qwen3-235b-a22b-2507`. Significance of correlations is assessed against a null population of surrogate maps with preserved spatial autocorrelation.

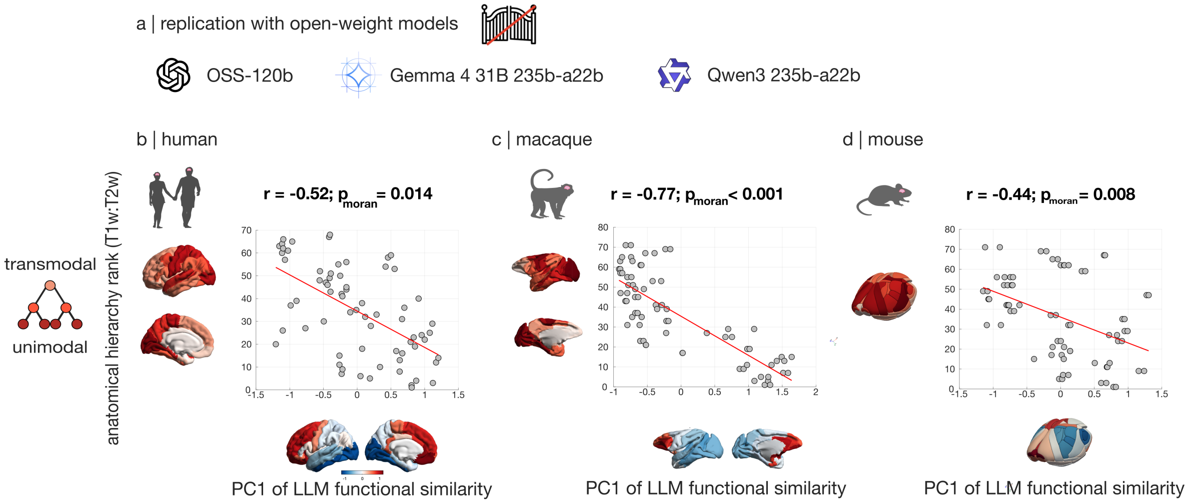


Figure S29. **Cortical hierarchy identified by LLMs replicates with open-weight models in each species |** Significant correlation between the PC1 of LLM-derived functional similarity and hierarchical position estimated using in vivo T1w:T2w ratio, is replicated in each species when using three frontier open-weight models: We replicate the main results using three frontier open-weight models: OpenAI's `OSS-120b`, Google's `Gemma-4-31b-instruct`, and Alibaba's `Qwen3-235b-a22b-2507`. **(a)** Human. **(b)** Macaque. **(c)** Mouse. Brain maps are displayed on the lateral and (for macaque and human) medial surface of the left cortical hemisphere.Significance of correlations is assessed against a null distribution of species-specific surrogate maps with preserved spatial autocorrelation.

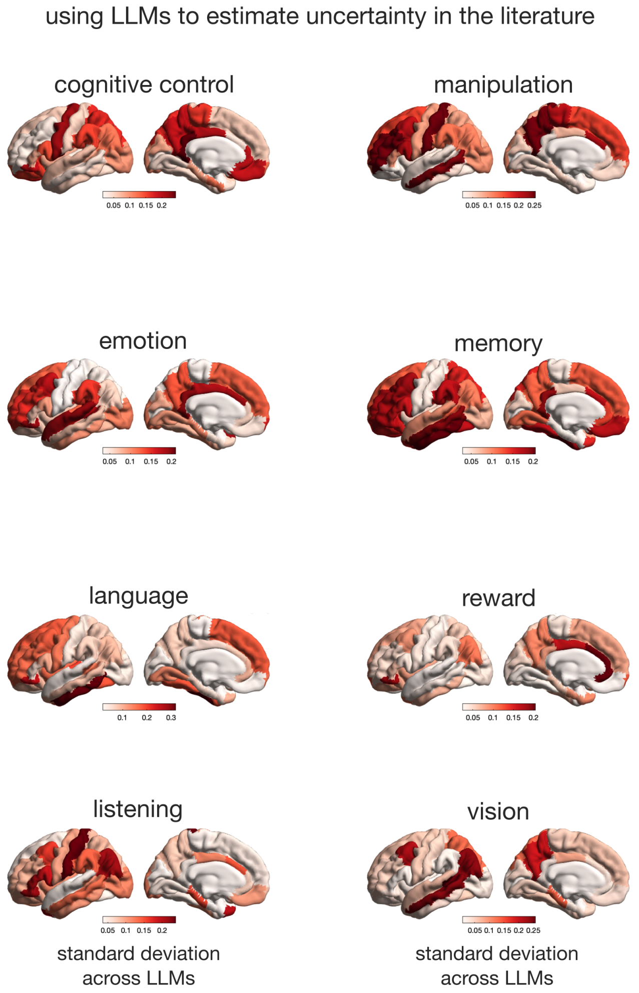


Figure S30. **Regional variability across LLMs for different cognitive functions |** For 8 representative domains of cognition, each brain region of the Desikan-Killiany atlas shows the divergence (standard deviation) between the values attributed by six different LLMs from the Claude, GPT, OSS, Gemini, Gemma, and Qwen families. Comparison with Fig. 1 shows that within each cognitive function, low variability is observed both for regions that are consistently attributed high region-function association, and for regions that are consistently assigned low region-function association. Brain maps are displayed on the lateral and medial surface of the left cortical hemisphere.

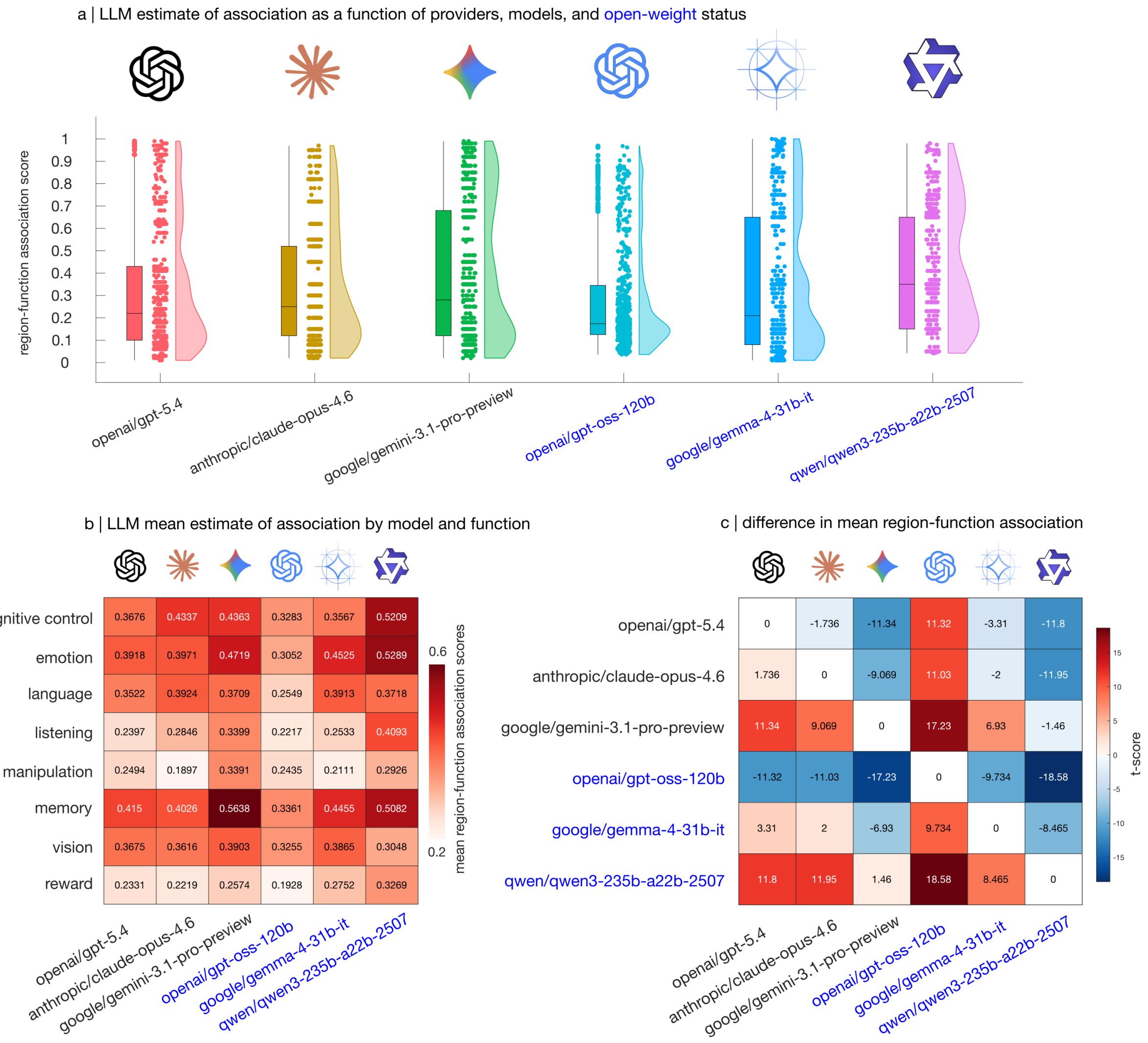


Figure S31. **Comparison of region-function association scores by model and function | (a)** Distribution of LLMs' region-function association scores across all region-function pairs, as a function of LLM model. **(b)** Heatmap shows LLM region-function association scores by LLM model and by 8 representative domains of cognition, averaging across regions. **(c)** Comparison of different LLMs' mean region-function association scores across region-function pairs, from (a). Colorbar indicates the t-score from a paired-samples t-test, such that positive values indicate row $>$ column. Box-plots: center line, median; box limits, upper and lower quartiles; whiskers, $1.5\times$ interquartile range. Each data-point represents the score attributed to one region for one function.

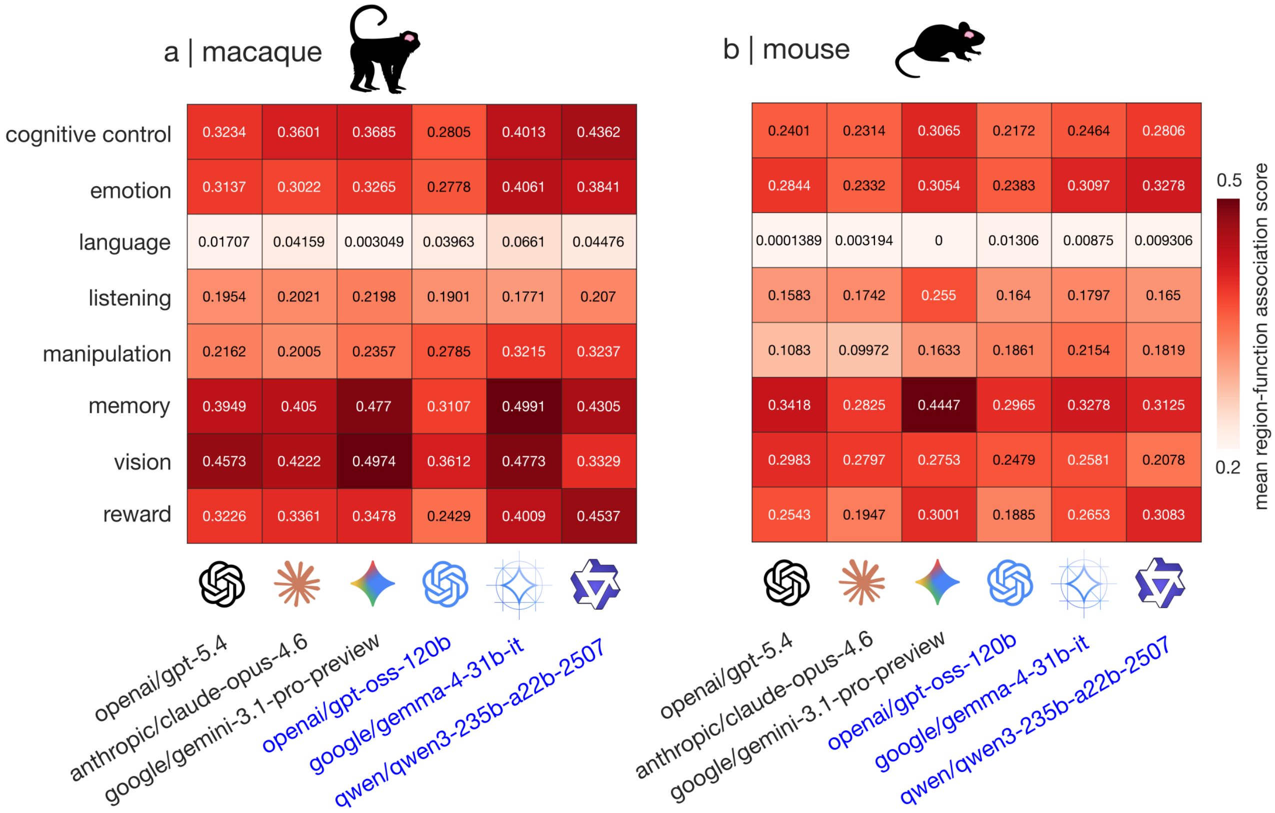


Figure S32. **AI experts consistently assign low probability of language in macaque and mouse | (a)** Heatmap shows LLM region-function association scores by LLM model and by 8 representative domains of cognition, averaged across regions of the macaque brain. **(b)** Heatmap shows LLM region-function association scores by LLM model and by 8 representative domains of cognition, averaged across regions of the mouse brain. All LLMs show a clear drop in estimated region-function association scores for “language”.

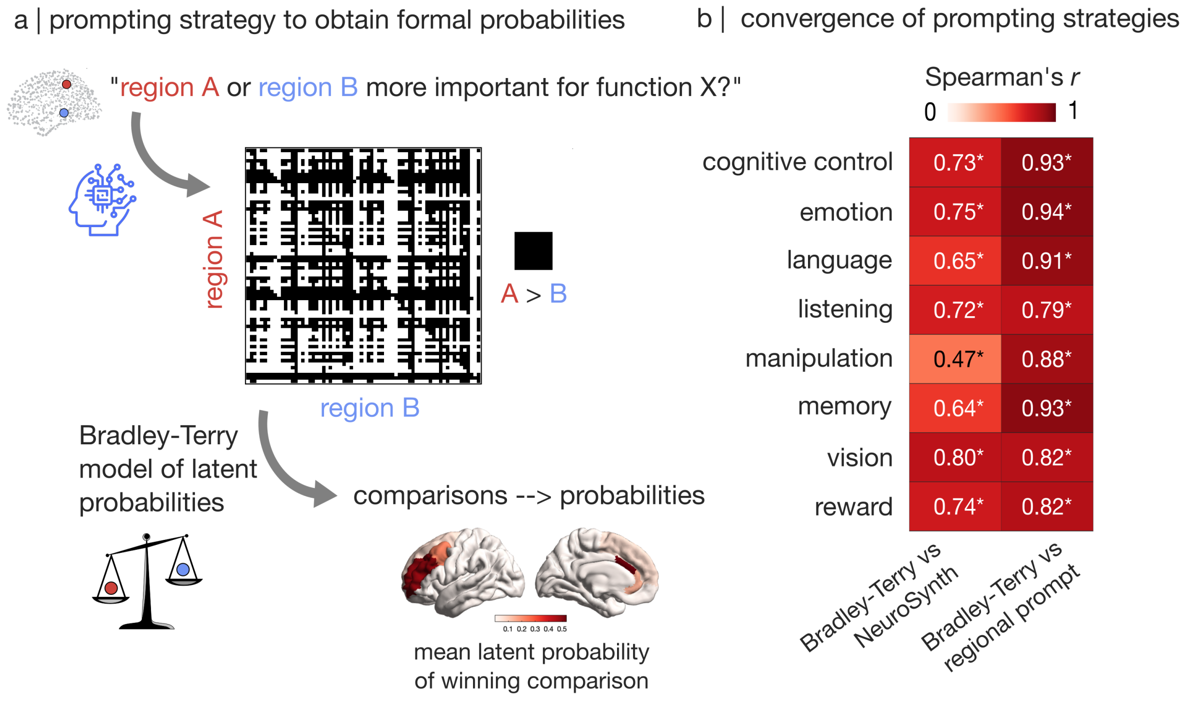


Figure S33. **Brain maps of latent probabilities from pair-wise comparisons are highly consistent with direct region-function association scores and NeuroSynth |** **(a)** For each cognitive function X, we ask each AI expert to decide whether region A or region B is more important. This process is repeated for every possible pair of brain regions. The resulting matrix of pairwise comparisons is provided as input to the Bradley-Terry model (76) (note that the diagonal is ignored) which infers a latent probability for each region, resulting in a brain map. These brain maps are then averaged across LLMs. **(b)** The brain maps of latent probabilities obtained from the Bradley-Terry model are highly and significantly correlated with the corresponding NeuroSynth maps, and also with the brain maps of region-function association scores obtained from the same LLMs. *, $p < 0.05$ against a null distribution of spatial autocorrelation-preserving surrogate brain maps.

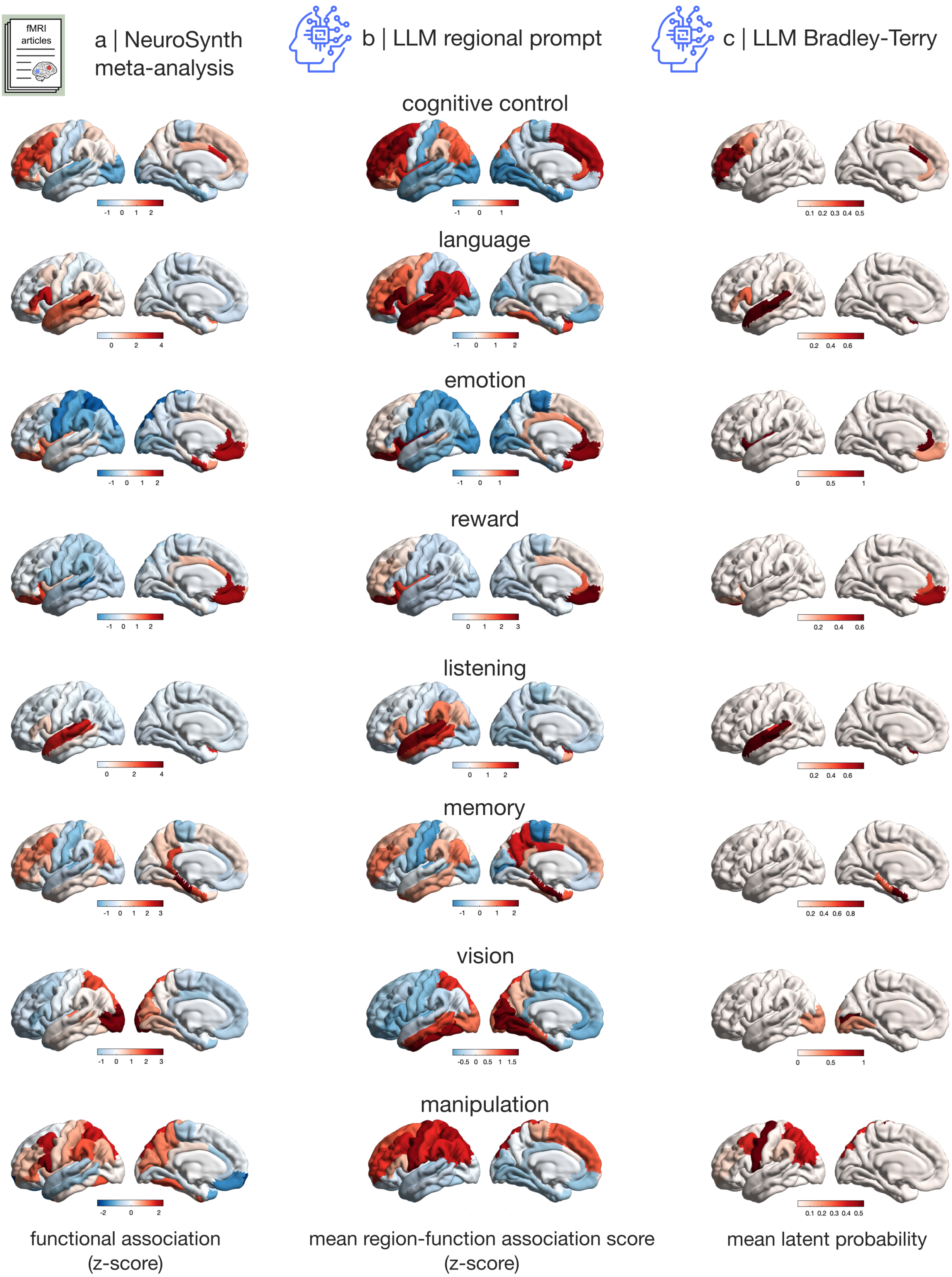


Figure S34. **Brain maps of latent probabilities from pair-wise comparisons |** **(a)** Brain maps from NeuroSynth automated meta-analysis. **(b)** Brain maps of region-function association scores, obtained by asking AI experts to assign a score to each region (maps are z-scored and then averaged across LLMs). **(c)** Brain maps of latent probability for different functions, obtained from the Bradley-Terry model based on direct comparisons between all possible pairs of regions (maps are averaged across LLMs). Brain maps are displayed on the lateral and medial surface of the left cortical hemisphere.

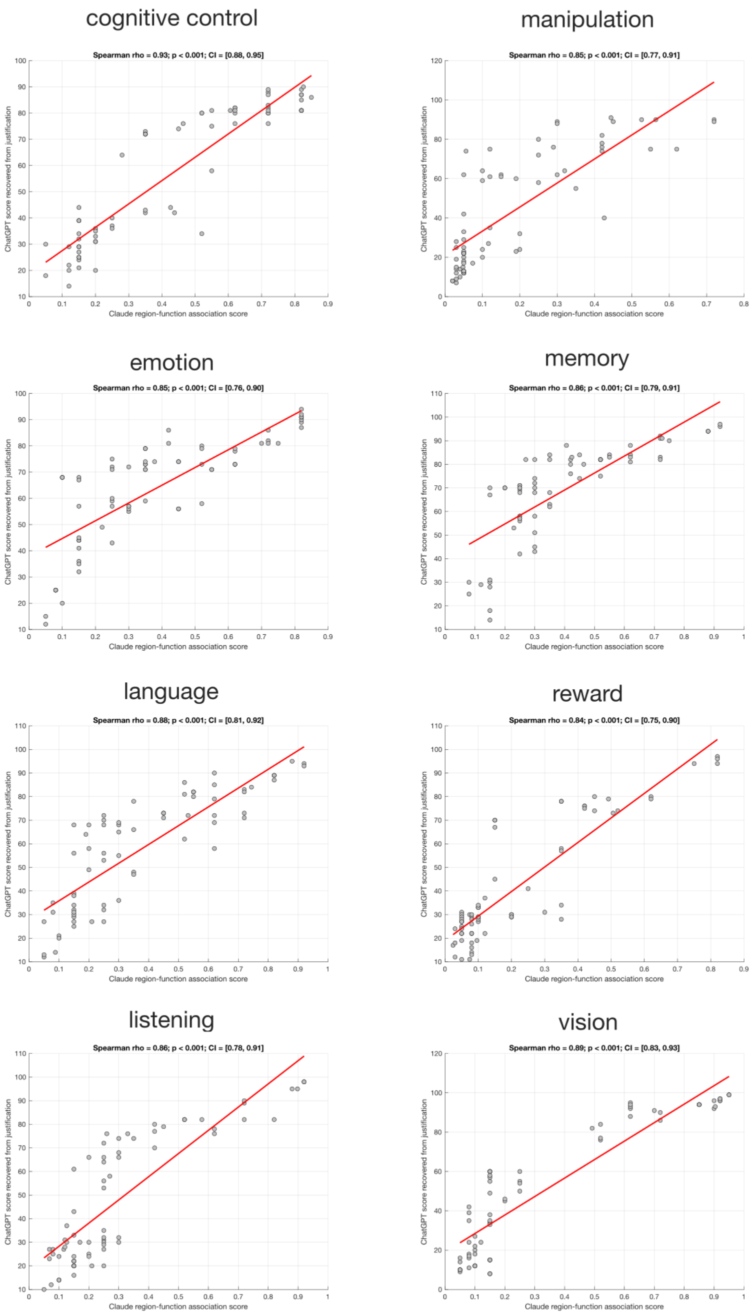


Figure S35. **AI experts' textual justifications are consistent with their region-function association scores |** For each of the 8 fundamental domains of cognition used in the main analysis, we provided ChatGPT-5.5 (not used in the main analyses) with the textual justifications provided by Claude, tasking it to rate the strength of evidence expressed in the statement, from 0 (minimum) to 100 (maximum). We find that the strength of evidence inferred by ChatGPT from text is strongly and significantly correlated with the region-function association score originally provided by Claude alongiside the justifications.

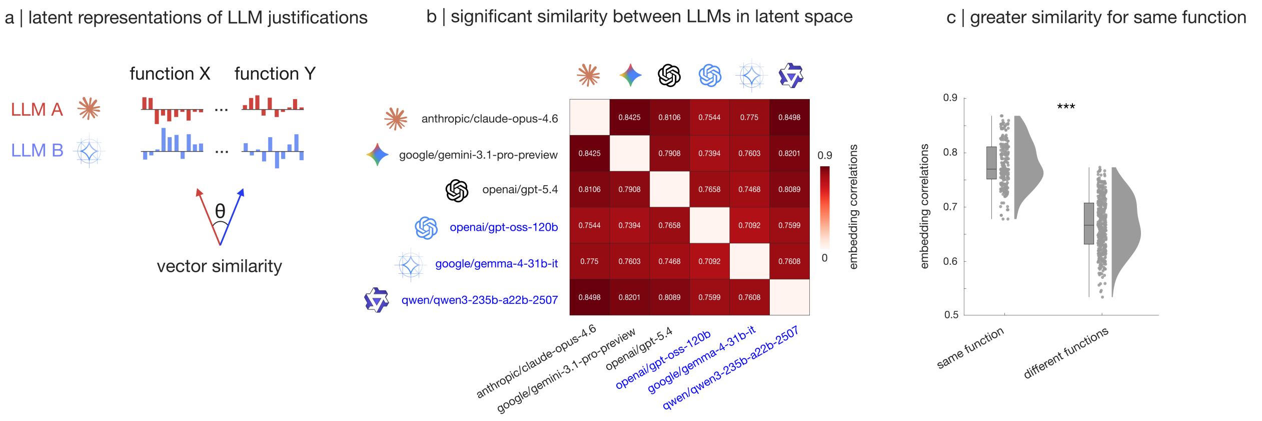


Figure S36. **Quantitative comparison of AI experts' justifications for their region-function association scores | (a)** LLMs are prompted to provide a brief textual justification for the region-function association score that they produce, via the `-justify` flag. Textual justifications for each region and function are then turned into 3072-dimensional text embeddings using the OpenAI API, which can then be compared in embedding space via correlation or cosine similarity. **(b)** Across functions, we observe strong correlations between the embeddings produced by different AI experts (all p < 0.05). **(c)** Across LLMs, the correlations between embeddings are significantly greater when they pertain to the same function, than for different functions. Same-function mean = 0.78, SD = 0.04; different-function mean = 0.67, SD = 0.05; t(658) = 29.53; p < 0.001; Hedge's $g$ = 2.38 from two-sided independent-samples t-test. Box-plots: center line, median; box limits, upper and lower quartiles; whiskers, $1.5\times$ interquartile range. *** $p < 0.001$.

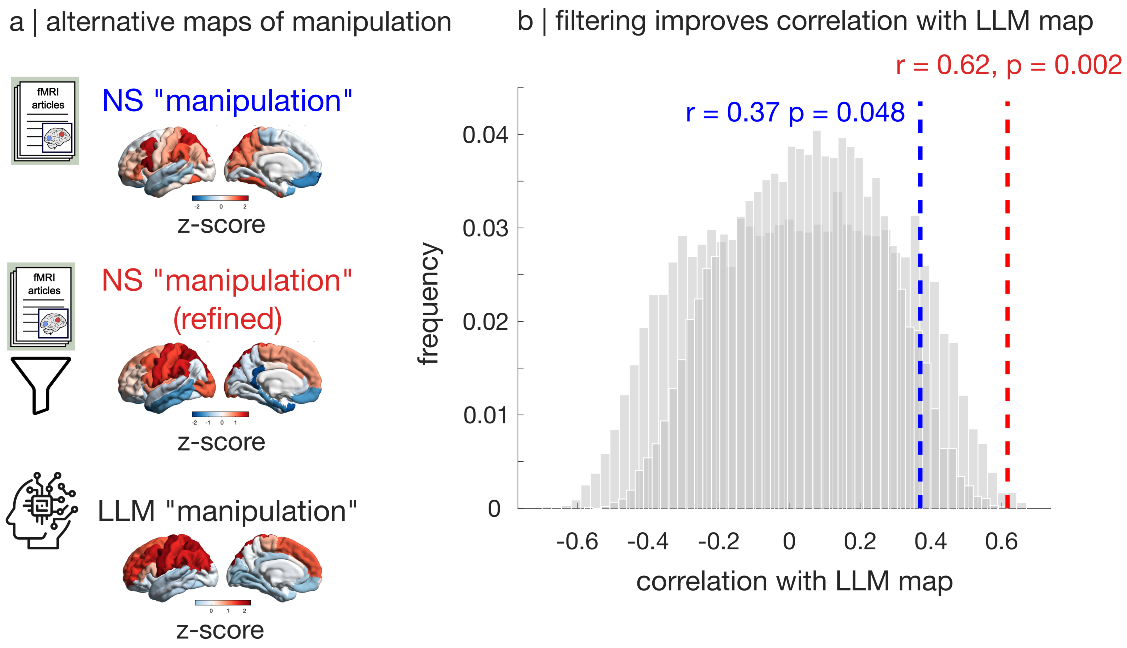


Figure S37. **Filtering out irrelevant meanings of 'manipulation' improves match between NeuroSynth and LLM-generated brain maps of manipulation | (a)** Brain map of 'manipulation' for NeuroSynth (top), NeuroSynth but after removing abstracts that used the term in a way incongruent with the intended meaning of 'object handling' (for example, in the sense of 'experimental manipulation' by the researchers) (middle); and the corresponding brain map generated by automated neuroscience experts (bottom). Brain maps are displayed on the lateral and medial surface of the left cortical hemisphere. **(b)** The spatial correspondence between maps of 'manipulation' generated by NeuroSynth and by LLMs improves drastically after filtering (Spearman's $r = 0.62, p = 0.002$). Significance of correlations is assessed against a population of null maps with preserved spatial autocorrelation.

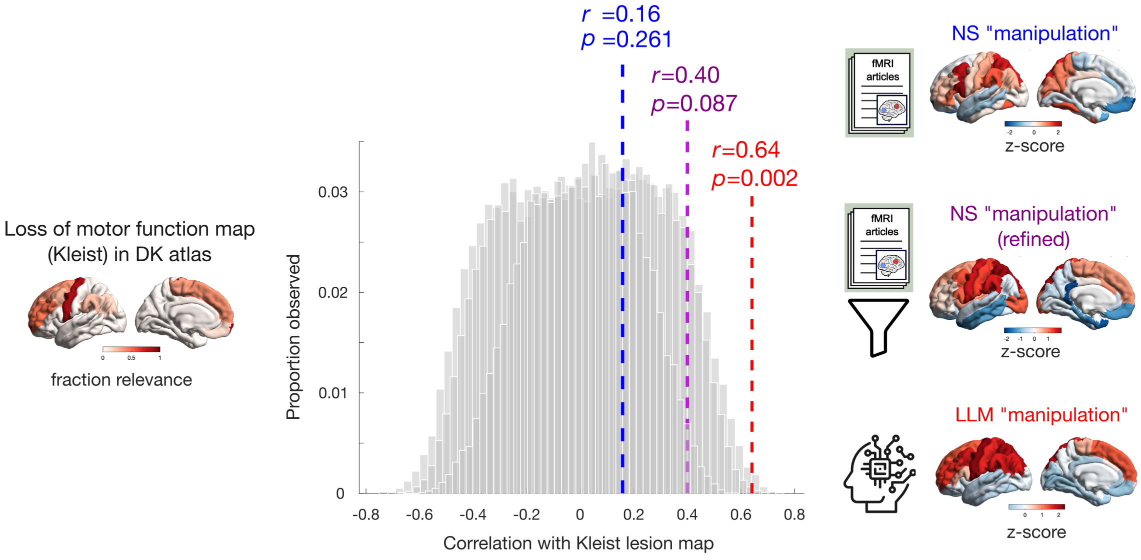


Figure S38. **LLM-generated brain map of manipulation coincides with lesion-induced loss of motor function |** A map of lesion-induced loss-of-function pertaining to motor skill is obtained from the classic work of Karl Kleist, who assigned functions to different Brodmann areas based on the deficits observed following traumatic brain injuries and focal brain lesions (137). We used the version of the Kleist atlas recently digitised and shared in MNI standard space by Pijnenburg et al. (138). The binary MNI-space map was resampled to the Desikan-Killiany atlas and correlated with maps of 'manipulation' generated by NeuroSynth (blue), by NeuroSynth after filtering out irrelevant meanings of 'manipulation' (purple), and by automated neuroscience experts (red). While filtering improves the correlation between the NeuroSynth map and the loss-of-function map, only the LLM-generated map exhibits a statistically significant correlation (Spearman's $r = 0.64, p = 0.002$). Significance of correlations is assessed against a population of null maps with preserved spatial autocorrelation. Brain maps are displayed on the lateral and medial surface of the left cortical hemisphere.

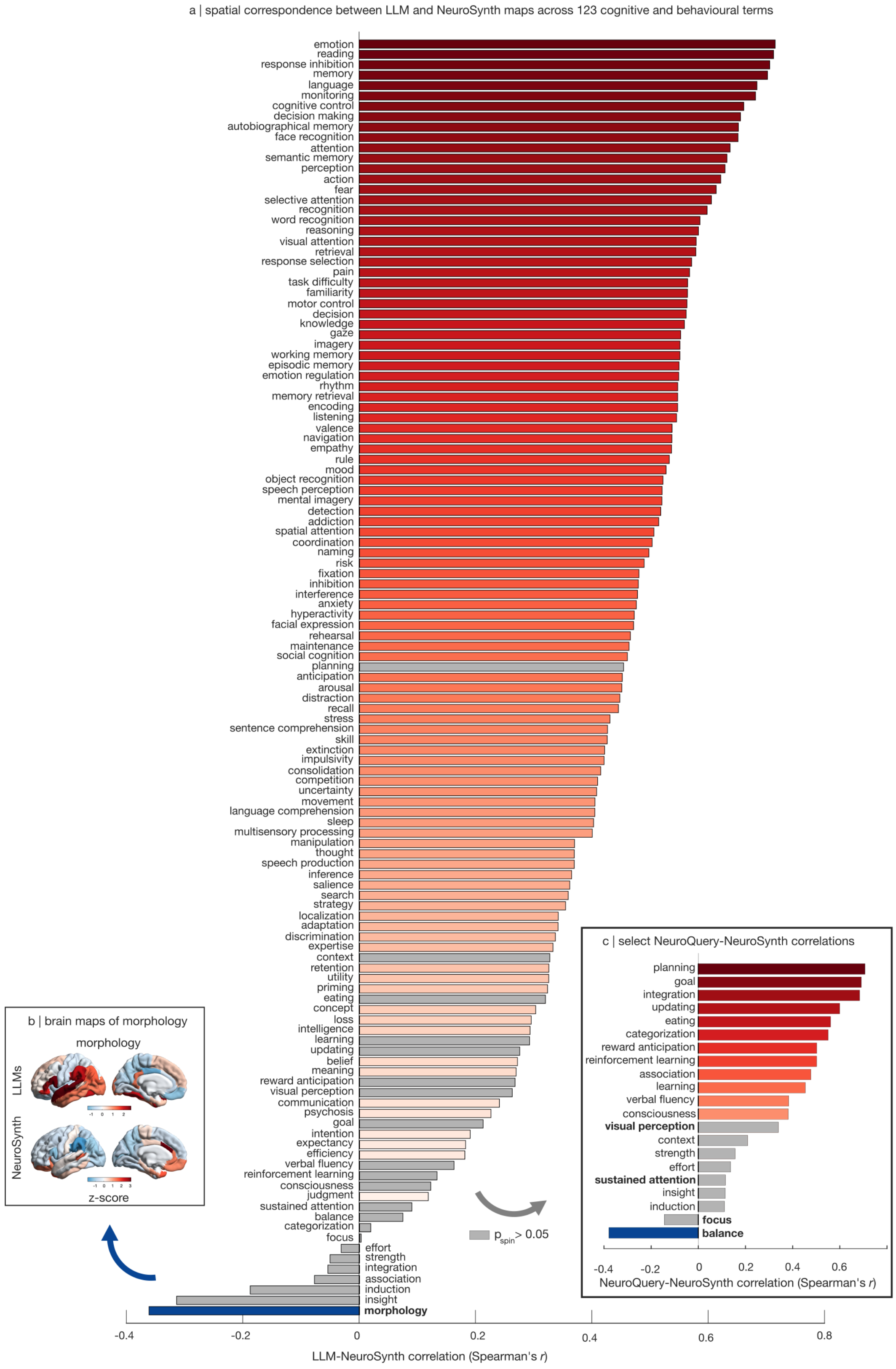


Figure S39. **Spatial correspondence between brain maps from NeuroSynth and from AI experts, for 123 cognitive operations | (a)** Spatial correlation (Spearman's $r$) between cognitive brain maps from AI experts and from NeuroSynth, for 123 cognitive operations. Grey bars indicate non-significant correlation. Significance of correlations is assessed against a population of null maps with preserved spatial autocorrelation. **(b)** Brain maps of 'morphology' from AI experts and from NeuroSynth. **(c)** Spatial correlation (Spearman's $r$) between cognitive brain maps from NeuroSynth and NeuroQuery, for the 21 cognitive operations where LLM-NeuroSynth correlation is non-significant. Grey bars indicate non-significant correlation. Significance of correlations is assessed against a population of null maps with preserved spatial autocorrelation. Brain maps are displayed on the lateral and medial surface of the left cortical hemisphere.

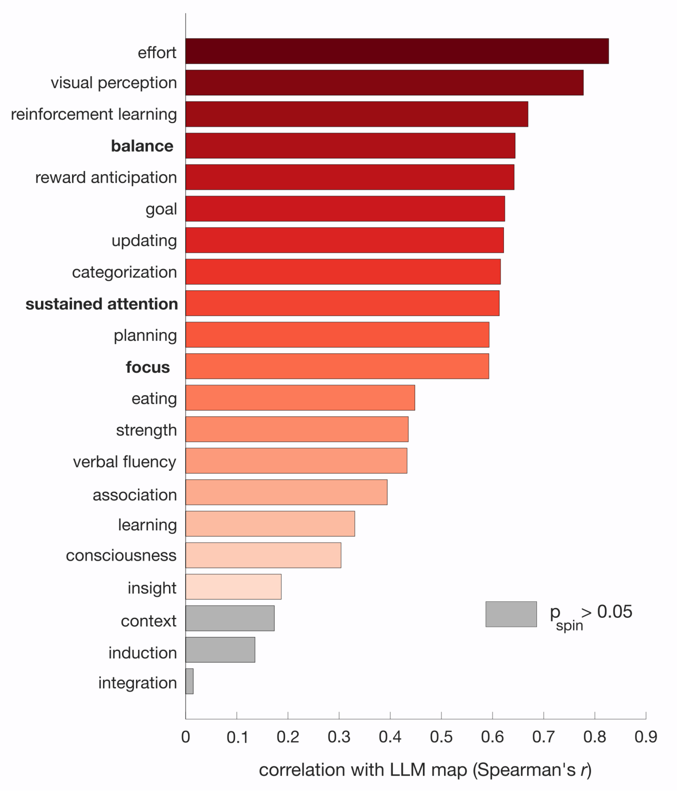


Figure S40. **Spatial correspondence between brain maps from NeuroSynth and from AI experts is improved by refining NeuroSynth abstract selection |** Spatial correlation (Spearman's *r*) between cognitive brain maps from AI experts and from NeuroSynth after refining the abstract selection, for the 21 cognitive operations that initially displayed non-significant correspondence. We see that 18/21 maps are now significantly correlated with the corresponding LLM-derived maps. Grey bars indicate non-significant correlation. Significance of correlations is assessed against a population of null maps with preserved spatial autocorrelation.

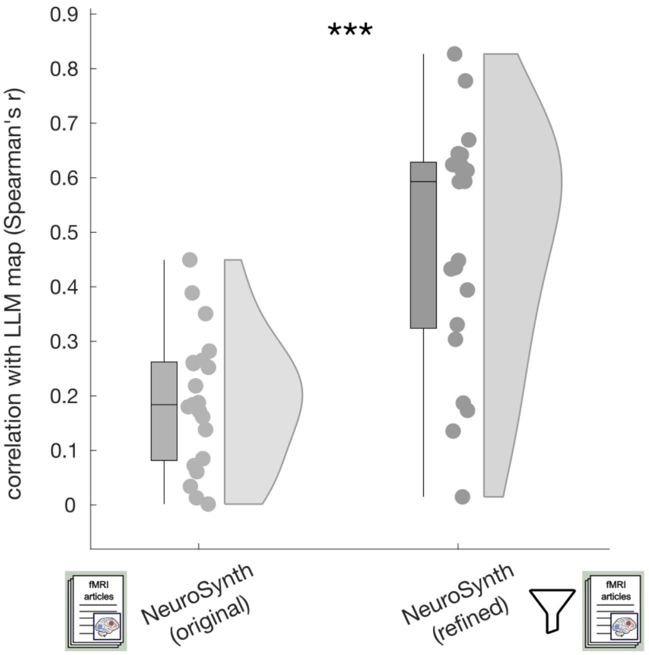


Figure S41. **Refining NeuroSynth abstract selection for inconsistent terms produces significantly greater correlations with corresponding LLM-generated maps |** Ordinate: magnitude of spatial correlation (Spearman's $r$) with the corresponding LLM maps, for the original NeuroSynth maps ($mean = 0.19, SD = 0.12$) and for the refined NeuroSynth maps ($mean = 0.48, SD = 0.22; t(20) = -5.66, p < 0.001$, Hedge's $g = -1.57$ from two-sided paired-samples t-test). Box-plots: center line, median; box limits, upper and lower quartiles; whiskers, $1.5\times$ interquartile range. Each data-point represents one term.

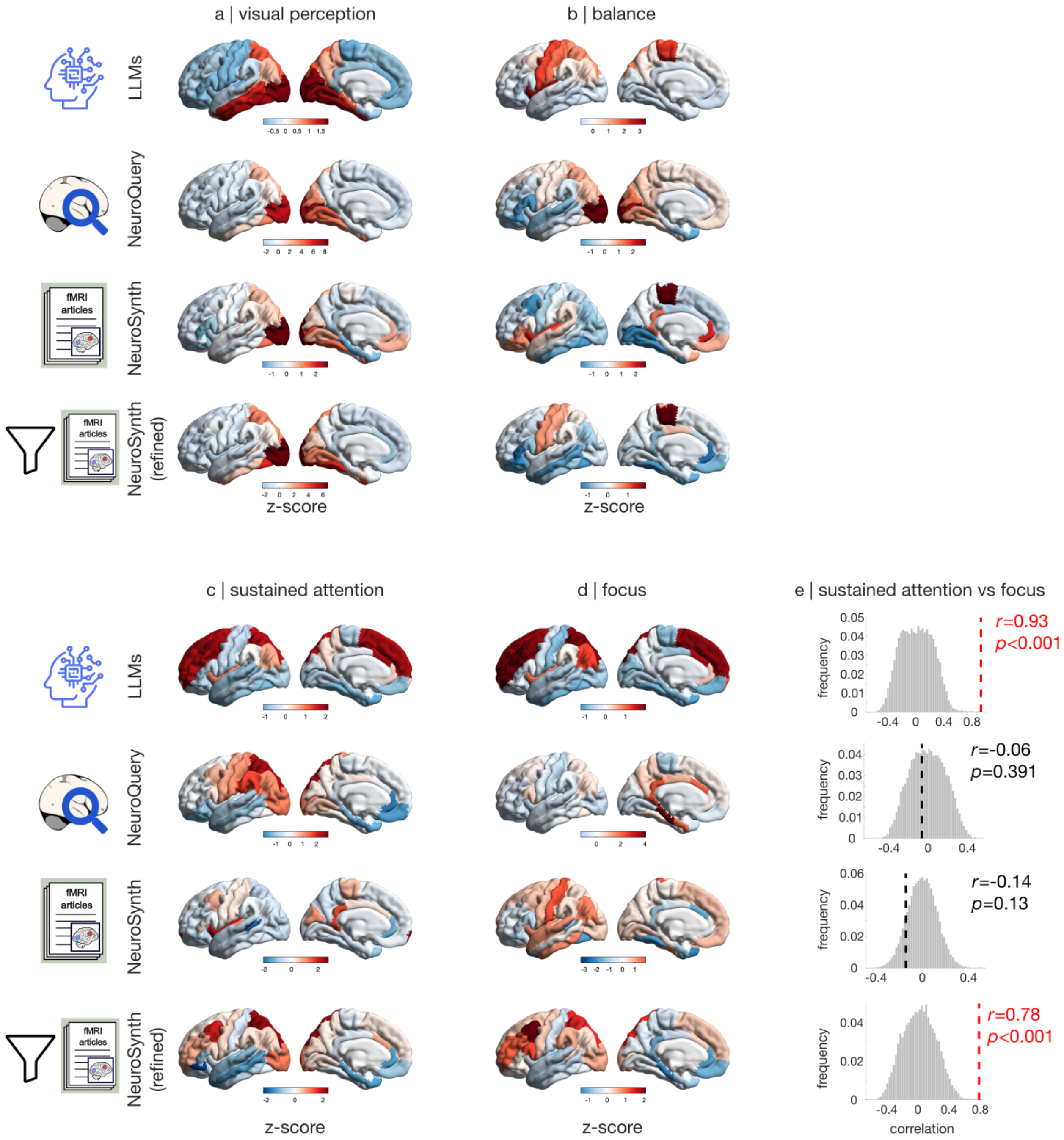


Figure S42. **Brain maps for example terms where AI experts, NeuroQuery, NeuroSynth, and the refined NeuroSynth provide divergent results | (a)** Brain maps of 'visual perception' from LLMs (top), NeuroQuery (second row), NeuroSynth (third row) and NeuroSynth after refining the abstract selection (bottom). **(b)** Brain maps of 'balance'. **(c)** Brain maps of 'sustained attention'. **(d)** Brain maps of 'focus'. **(e)** Brain maps of 'sustained attention' and 'focus' generated by LLMs are significantly correlated (Spearman $r = 0.93, p < 0.001$). No significant correlation between 'sustained attention' and 'focus' maps is observed for either NeuroQuery (Spearman $r = 0.06, p = 0.39$) or NeuroSynth (Spearman $r = -0.14, p = 0.13$), but we do find a significant correlation for NeuroSynth after refining the abstract selection (Spearman $r = 0.78, p < 0.001$), which also brings both maps to resemble the corresponding maps generated by AI experts (significantly so: Fig. S40). Significance of correlations is assessed against a null population of surrogate maps with preserved spatial autocorrelation. Brain maps are z-scored across regions and displayed on the lateral and medial surface of the left cortical hemisphere.

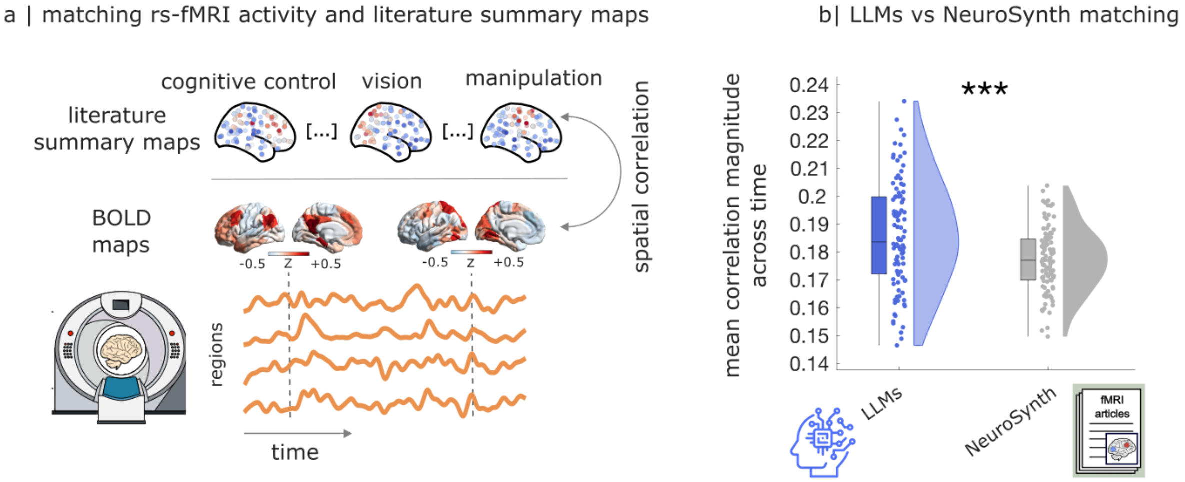


Figure S43. **LLM-generated brain maps of cognition are a better match for human spontaneous fMRI activity than NeuroSynth-generated maps | (a)** At each point in time, the BOLD signal volume is treated as a brain map, and spatially correlated with meta-analytic maps generated by LLMs, or by NeuroSynth. **(b)** Across 100 unrelated volunteers from the Human Connectome Project, the mean magnitude of spatial correlation over time is significantly higher for the LLM-generated maps ($mean = 0.19, SD = 0.02$), than for the NeuroSynth-generated maps ($mean = 0.18, SD = 0.01; t(99) = 6.73, p < 0.001$, Hedge's $g = 0.55$ from two-sided paired-samples t-test). Box-plots: center line, median; box limits, upper and lower quartiles; whiskers, $1.5\times$ interquartile range. Each data-point is one individual.

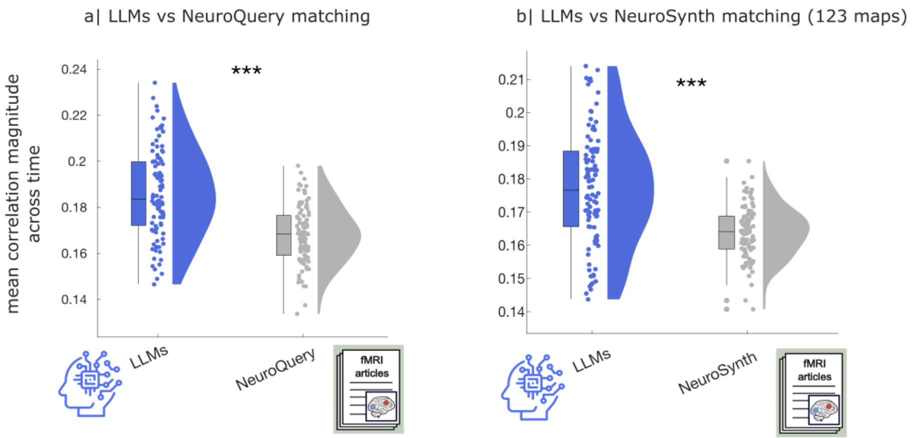


Figure S44. **LLM-generated brain maps of cognition are a better match for human spontaneous fMRI activity than NeuroSynth- or NeuroQuery-generated maps | (a)** Across 100 unrelated volunteers from the Human Connectome Project, the mean magnitude of spatial correlation over time is significantly higher for the LLM-generated maps ($mean = 0.19, SD = 0.02$), than for the NeuroQuery-generated maps ($mean = 0.17, SD = 0.01; t(99) = 18.20, p < 0.001$, Hedge's $g = 1.10$) **(b)** Replication of the NeuroSynth results when using brain maps for 123 cognitive operations (Table 2). LLM-generated maps ($mean = 0.18, SD = 0.02$); NeuroSynth-generated maps ($mean = 0.16, SD = 0.01; t(99) = 10.39, p < 0.001$, Hedge's $g = 0.98$ from two-sided paired-samples t-test). Box-plots: center line, median; box limits, upper and lower quartiles; whiskers, $1.5\times$ interquartile range. Each data-point is one individual.

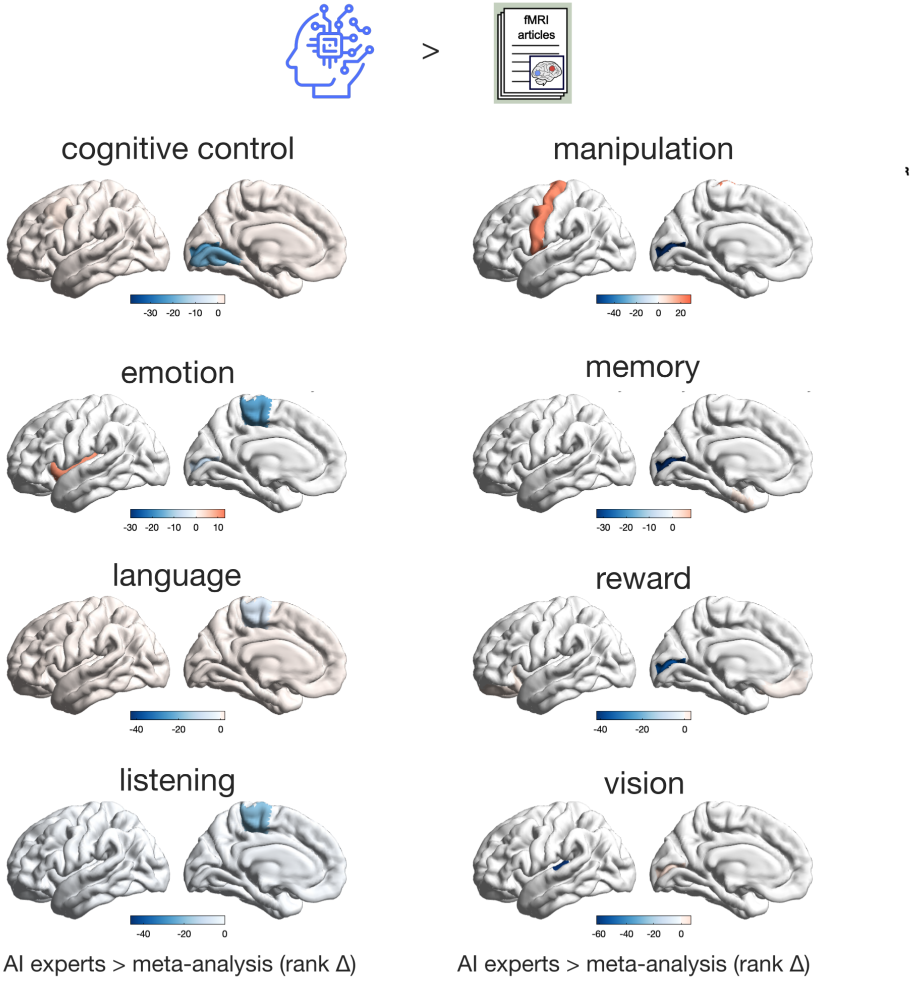


Figure S45. **Systematic differences between LLM-generated and NeuroSynth-generated brain maps |** Significant differences in rank between LLM-generated and NeuroSynth-generated maps, highlighting how NeuroSynth consistently assigns higher ranks to visual and motor regions. Significance is assessed against a population of null maps with preserved spatial autocorrelation. Brain maps are displayed on the lateral and medial surface of the left cortical hemisphere.

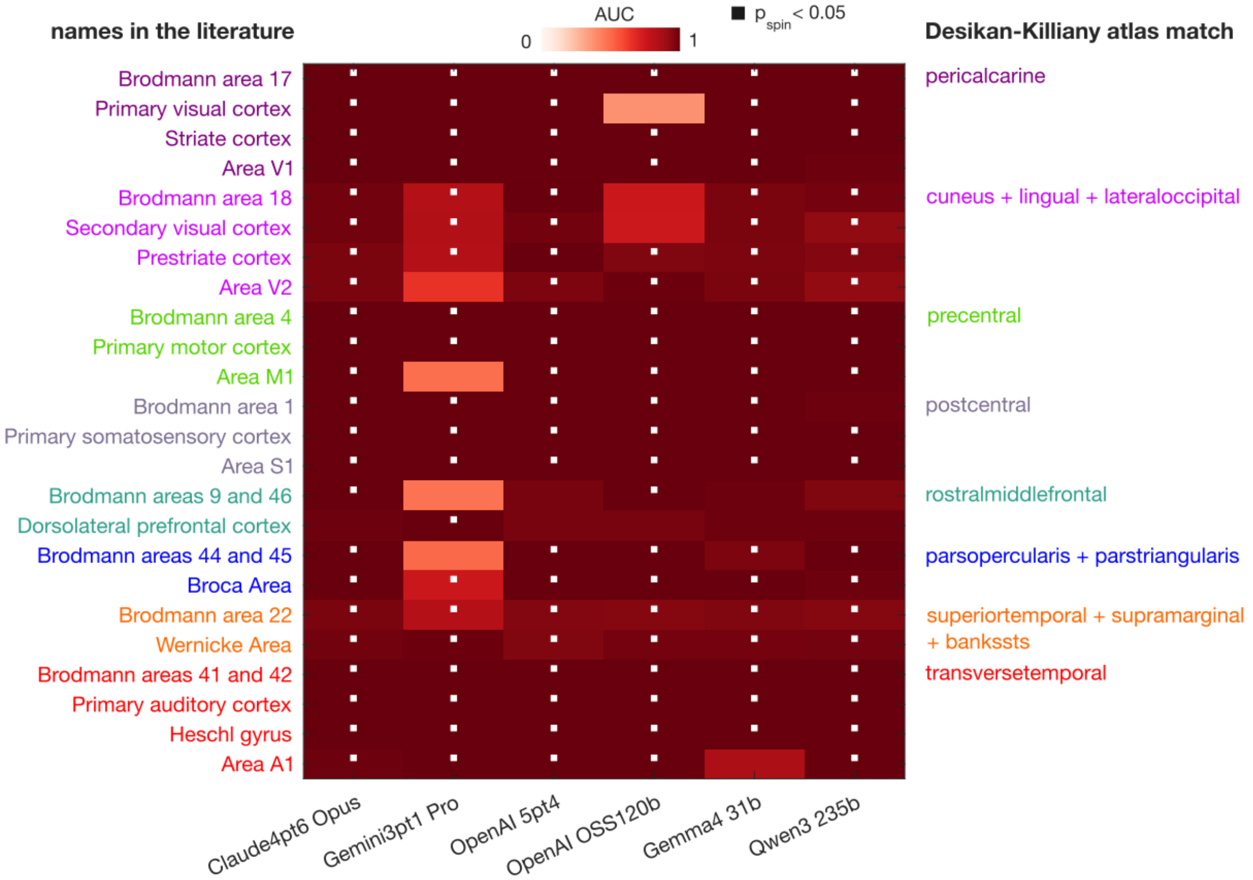


Figure S46. **AI experts can match different names for the same cortical region |** For each region in the Desikan-Killiany atlas, LLMs are asked for the probability that it refers to the same part of the brain as a target region (for example, 'V1'). We repeat this for multiple target regions, each with several synonyms encompassing anatomical, cytoarchitectonic, and functional labels. Terms in the same colour denote the same region. For each target region, we evaluated how well the LLM-derived regional relevance vector matched the corresponding anatomical ground truth. The ground-truth vector was defined over Desikan–Killiany atlas parcels, with entries set to 1 for parcels expected to correspond to the target region and 0 for all other parcels. For example, when the target concept was V1, the *pericalcarine* parcel was labelled as the positive ground-truth region, while all other parcels were labelled as negatives. We then computed the area under the receiver operating characteristic curve (ROC-AUC) between the LLM-provided scores and this binary ground-truth vector. This measures whether the LLM assigns higher scores to anatomically appropriate parcels than to unrelated parcels. Cells with a white dot indicate that the LLM-generated brain map has an AUROC significantly greater than the AUROCs of a null distribution of surrogate maps with preserved spatial autocorrelation.

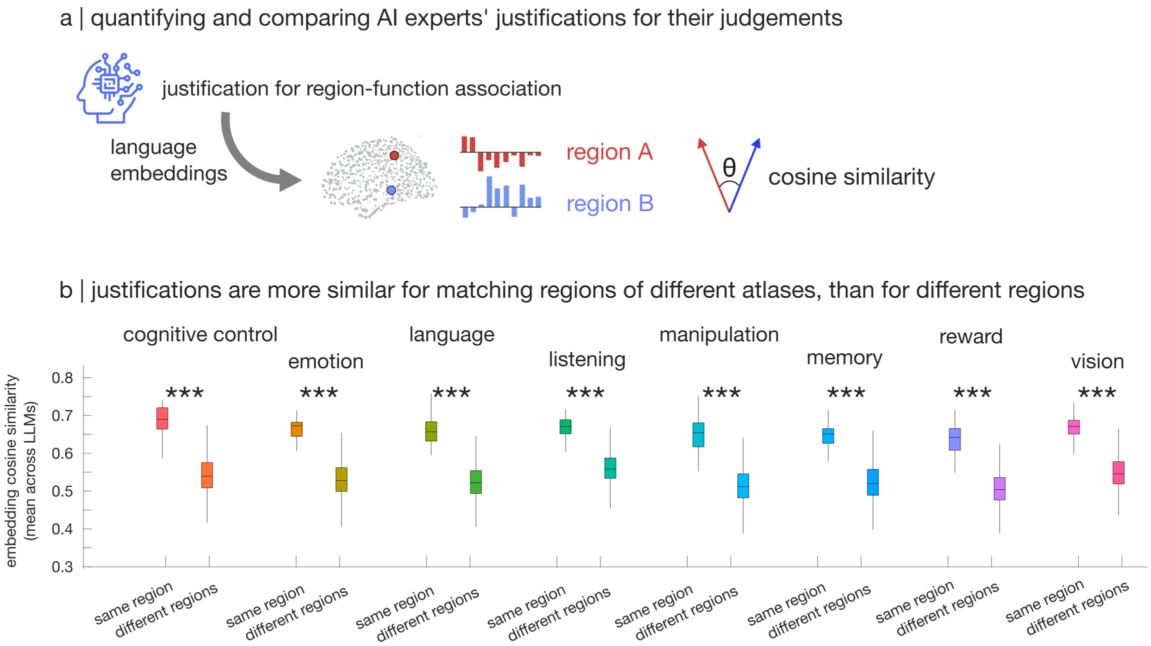


Figure S47. **AI experts' textual justifications are more similar when they pertain to different names of the same cortical area, than different areas | (a)** LLMs are prompted to provide a brief textual justification for the region-function association score that they produce, via the `-justify` flag. Textual justifications for each region and function are then turned into 3072-dimensional text embeddings using the OpenAI API, which can then be compared in embedding space via correlation or cosine similarity. **(b)** Across functions, justifications are consistently more aligned in embedding space when they refer to matching regions of different atlases (for example, 'pericalcarine' in Desikan-Killiany and 'Brodmann area 17' in the Brodmann atlas), than for mis-matched regions. *** $p < 0.001$ from two-sided independent-samples t-tests. Box-plots: center line, median; box limits, upper and lower quartiles; whiskers, $1.5\times$ interquartile range.

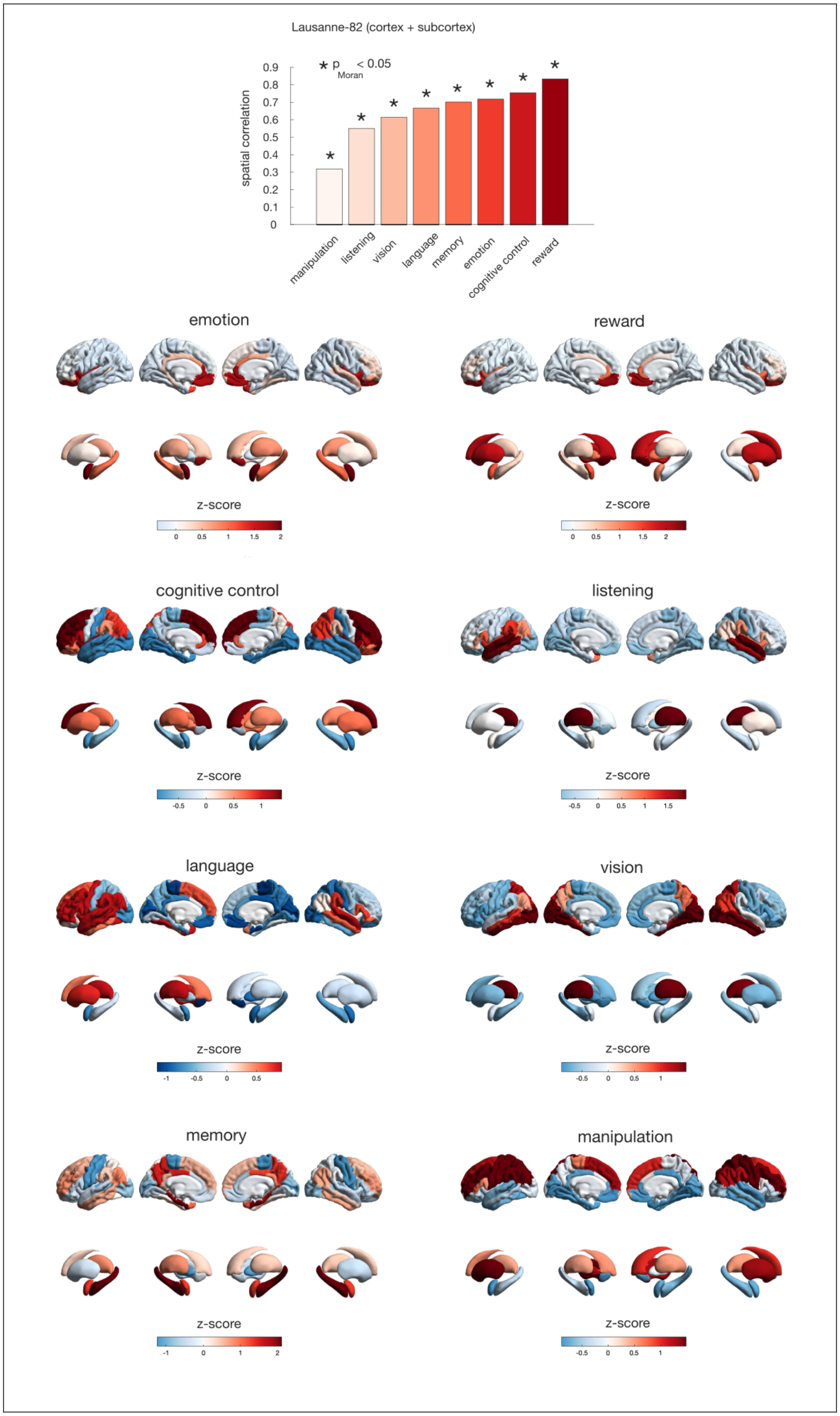


Figure S48. **LLMs generate brain maps of cognition including subcortex | (Top)** Correlation between NeuroSynth meta-analytic maps and LLM-generated maps is replicated when including subcortical structures. Significance of correlations is assessed against a population of null maps with preserved spatial autocorrelation generated using Moran spectral randomisation. **(Bottom)** LLM-generated brain maps with subcortex. Brain maps are displayed on the lateral and medial surface of the left and right cortical hemispheres, with corresponding subcortical views underneath.

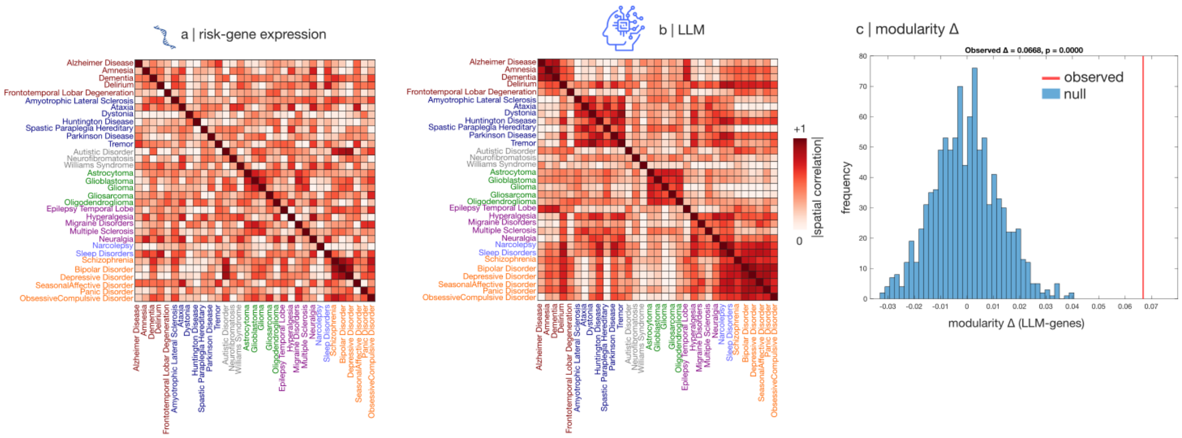


Figure S49. **LLMs recapitulate relationships between disorders better than risk-gene expression, even when including subcortical structures | (a,b)** Similarity (magnitude of spatial correlation) between disease maps from LLM-quantified regional involvement (b) aligns with symptom dimensions significantly better than similarity from transcriptomically derived maps of molecular risk (a). Modular structure is based on assignment into 7 disease categories from the International Classification of Diseases manual: Neurocognitive disorders and Dementias; Neurodegenerative and movement disorders; Neurodevelopmental disorders; Brain tumors; Neurological disorders; Sleep and Wake disorders; and Mood and Behaviour disorders. **(c)** The difference in modular structure between LLM similarity and molecular similarity is significantly greater than chance.

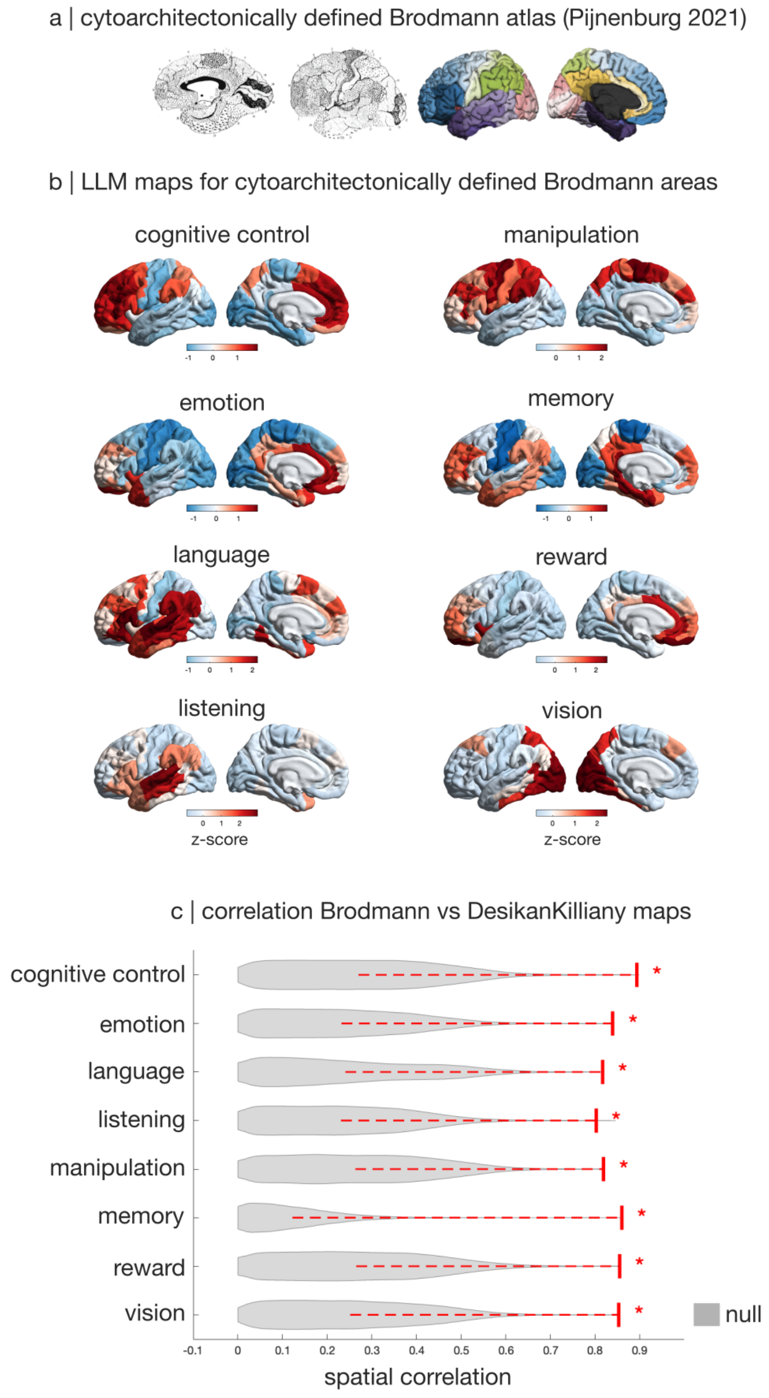


Figure S50. **LLM-generated brain maps of cognition replicate with a cytoarchitectonic atlas |** **(a)** The cytoarchitectonic Brodmann atlas, in original version and the MNI standard-space version digitised and made available by Pijnenburg et al. (138). **(b)** LLM-derived maps of data-driven domains of human cognition from Beam et al. (69) obtained with the Brodmann atlas (e.g., 'Brodmann Area 17' for primary visual cortex). Brain maps are displayed on the lateral and medial surface of the left cortical hemisphere. **(c)** Significant and strong spatial correspondence between LLM-generated maps in the original Desikan-Killiany atlas (from Figure 1), and the Brodmann-atlas maps from panel (b), re-parcellated into the Desikan-Killiany atlas.

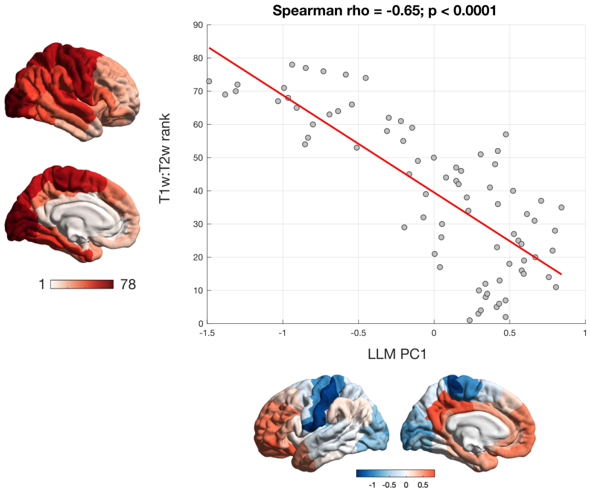


Figure S51. **Cortical hierarchy identified by LLMs replicates with a cytoarchitectonic atlas |** Significant correlation between the PC1 of LLM-derived functional similarity obtained using the cytoarchitectonic Brodmann atlas (abscissa), and hierarchical position estimated using in vivo T1w:T2w ratio (ordinate). Significance of correlations is assessed against a population of null maps with preserved spatial autocorrelation. Brain maps are displayed on the lateral and medial surface of the left cortical hemisphere.

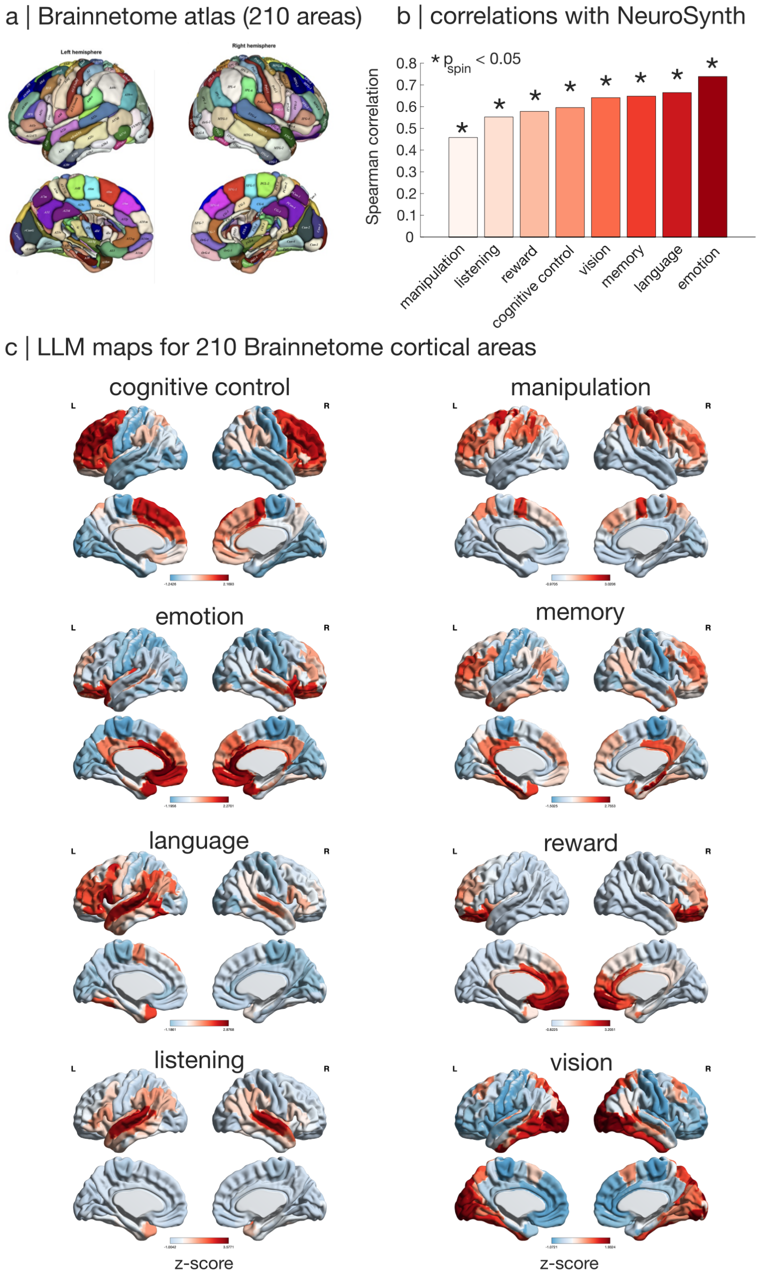


Figure S52. **LLM-generated brain maps of cognition replicate with the multimodal Brainnetome atlas | (a)** The multimodal Brainnetome atlas of Fan et al. (256), with 210 cortical regions. Adapted from (256). **(b)** Significant correlations between NeuroSynth meta-analytic maps and LLM-generated maps in the Brainnetome parcellation. Significance of correlations is assessed against a population of null maps with preserved spatial autocorrelation. **(c)** LLM-generated brain maps for the 8 data-driven domains of human cognition, in the Brainnetome parcellation. Brain maps are displayed on the lateral and medial surface of the left and right cortical hemispheres.

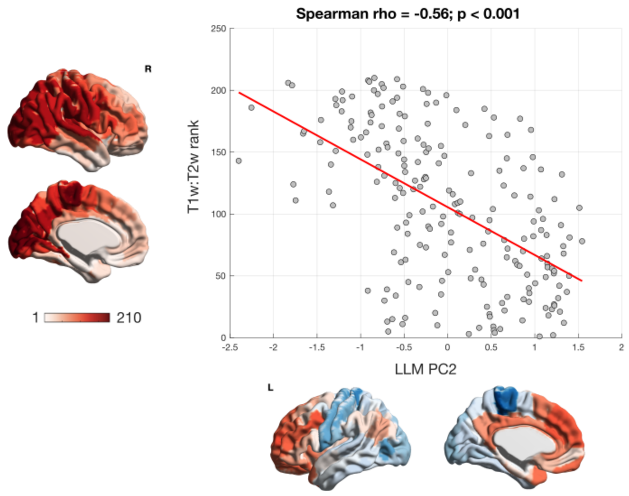


Figure S53. **Cortical hierarchy identified by LLMs replicates with the multimodal Brainnetome atlas |** Significant correlation between the PC1 of LLM-derived functional similarity obtained using the finer-grained Brainnetome parcellation with 210 cortical regions (abscissa), and hierarchical position estimated using in vivo T1w:T2w ratio (ordinate). Significance of correlations is assessed against a population of null maps with preserved spatial autocorrelation. Brain maps are displayed on the lateral and medial cortical surface.

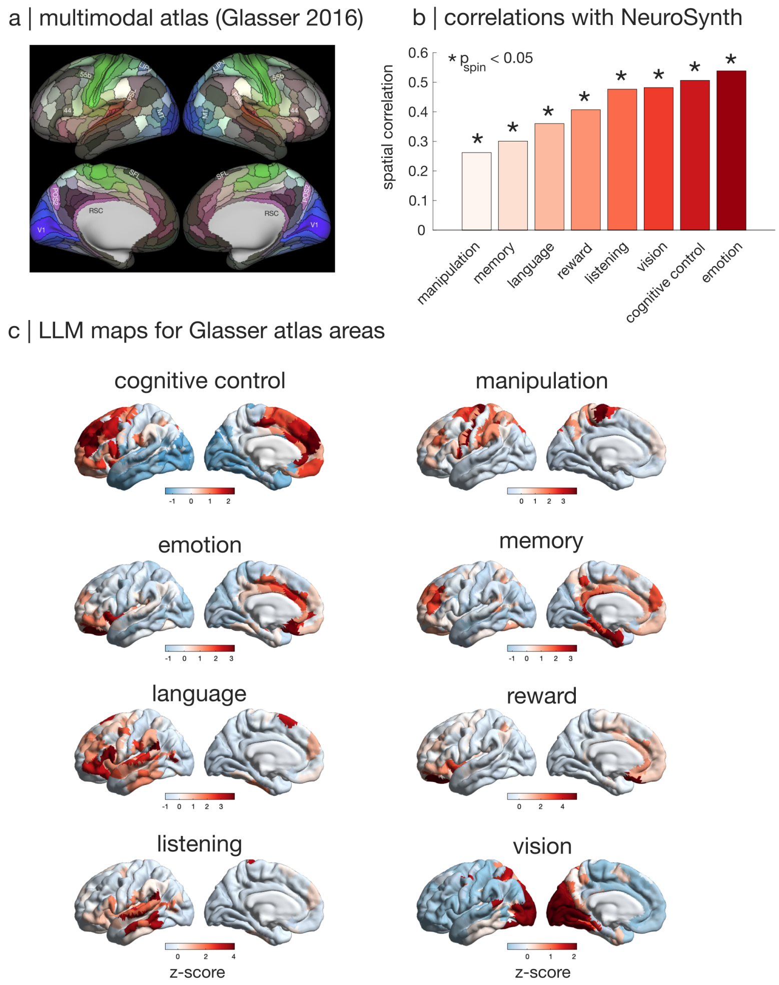


Figure S54. **LLM-generated brain maps of cognition replicate the fine-grained multimodal Glasser atlas | (a)** The multimodal atlas of Glasser et al. (160), with 360 cortical regions instead of 68. Note that more than half of the regions in this parcellation (194 out of 360) were not previously described in the literature. Adapted from (160). **(b)** Significant correlations between NeuroSynth meta-analytic maps and LLM-generated maps in the Glasser parcellation. Significance of correlations is assessed against a population of null maps with preserved spatial autocorrelation. **(c)** LLM-generated brain maps for the 8 data-driven domains of human cognition, in the Glasser parcellation. Brain maps are displayed on the lateral and medial surface of the left cortical hemisphere.

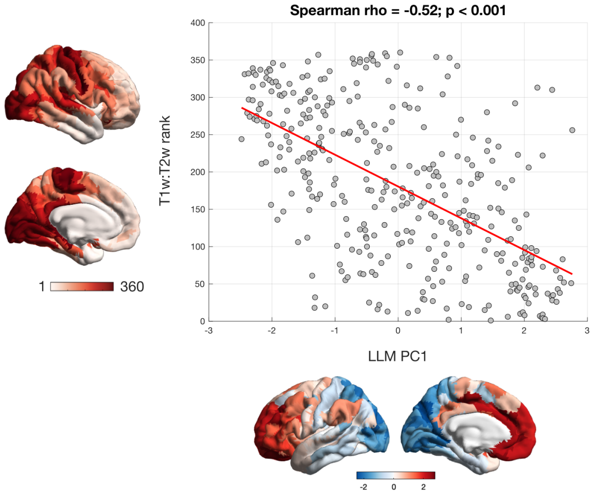


Figure S55. **Cortical hierarchy identified by LLMs replicates the fine-grained multimodal Glasser atlas |** Significant correlation between the PC1 of LLM-derived functional similarity obtained using the finer-grained Glasser parcellation with 360 cortical regions (abscissa), and hierarchical position estimated using in vivo T1w:T2w ratio (ordinate). Significance of correlations is assessed against a population of null maps with preserved spatial autocorrelation. Brain maps are displayed on the lateral and medial cortical surface.

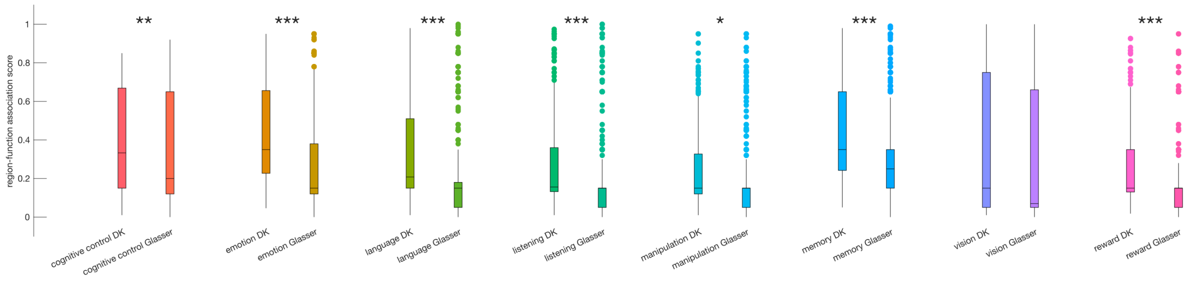


Figure S56. **AI experts assign higher region-function association scores to regions of the Desikan-Killiany than Glasser atlas |** Distributions of region-function association scores for different regions and different LLMs, separately for the 8 cognitive functions used in the main analysis. *, $p < 0.05$; **, $p < 0.01$; ***, $p < 0.001$ from independent-samples permutation-based t-tests (two-sided). Box-plots: center line, median; box limits, upper and lower quartiles; whiskers, $1.5\times$ interquartile range.

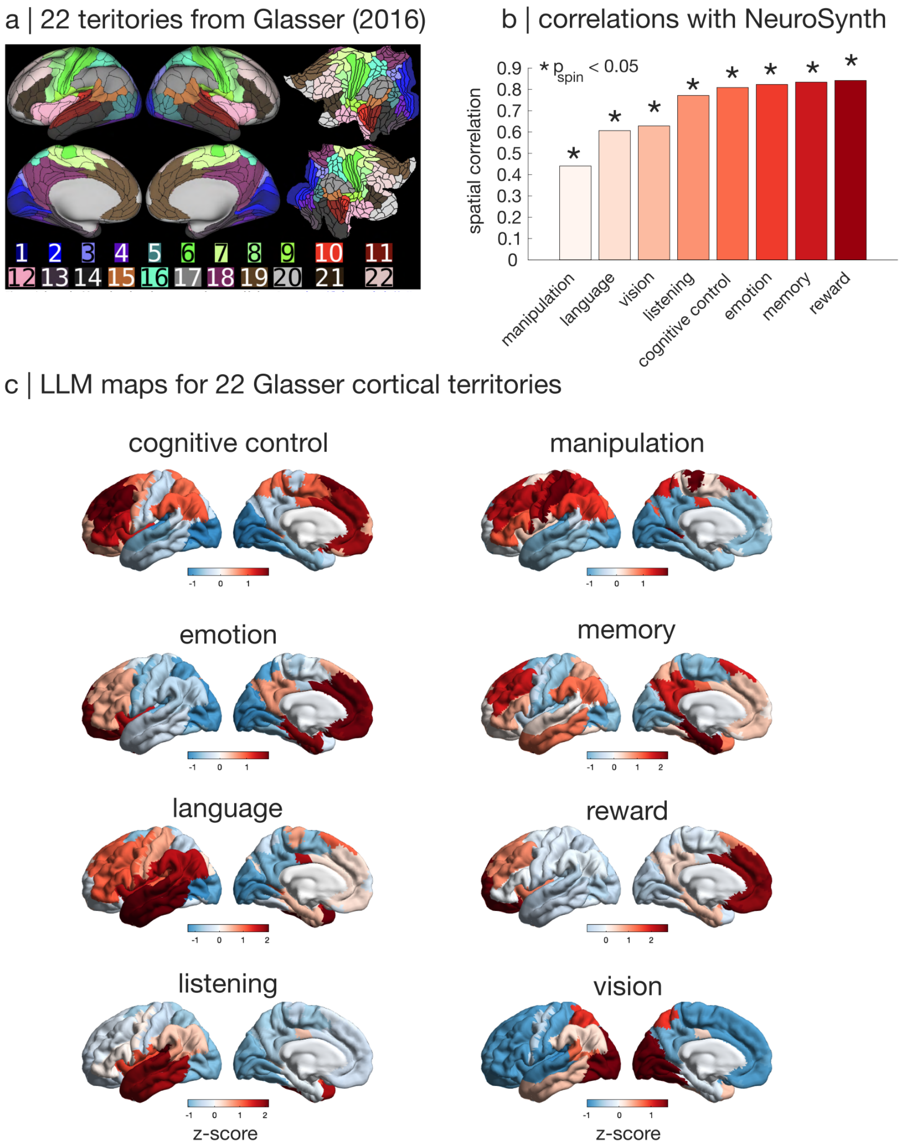


Figure S57. **LLM-generated brain maps of cognition replicate with a coarser-grained subdivision of the Glasser multimodal atlas | (a)** The 180 unique regions of the Glasser multimodal atlas are grouped into 22 cortical territories (as defined in the Supplementary Materials of (160)). Adapted from (160). **(b)** Significant correlations between NeuroSynth meta-analytic maps and LLM-generated maps for this coarser version of the Glasser parcellation. Significance of correlations is assessed against a population of null maps with preserved spatial autocorrelation. **(c)** LLM-generated brain maps for the 8 data-driven domains of human cognition, in the 22 cortical territories defined by Glasser et al. (160). Brain maps are displayed on the lateral and medial surface of the left cortical hemisphere.

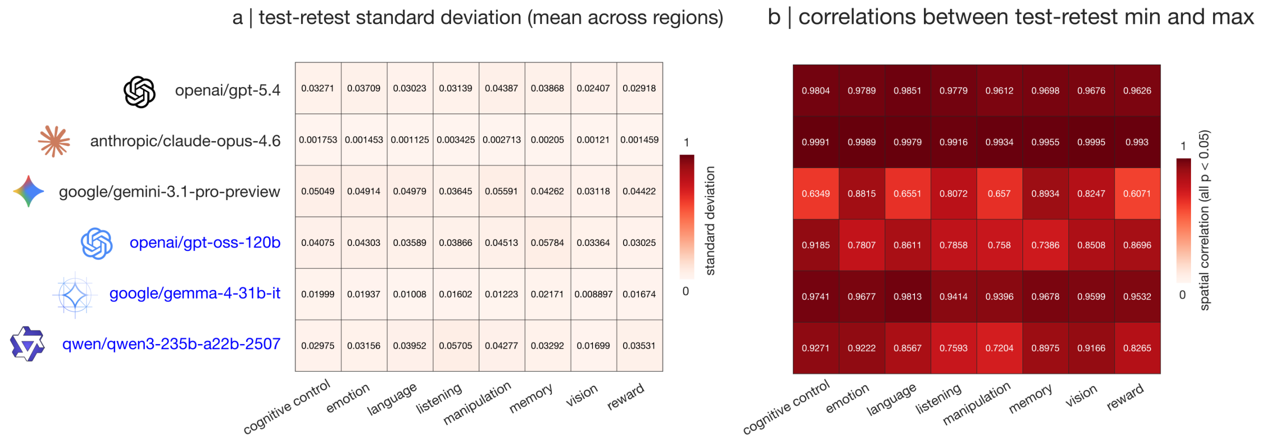


Figure S58. **LLM-generated maps are consistent across different test-retest iterations of the workflow | (a)** Standard deviation of region-function association scores produced by each LLM across 10 iterations, for 8 representative domains of cognition. We consistently observe very low values, approximately ranging between 0.001 and 0.05 on average, indicating high consistency within the same model upon repeated queries. **(b)** Spatial correlations between the two most extreme maps (most conservative and most liberal) generated by each LLM for each function (i.e., the maps of minimum and maximum observed region-function association). All correlations are significant against a null distribution of spatial autocorrelation-preserving surrogate maps.
